# EfgA is a conserved formaldehyde sensor that halts bacterial translation in response to elevated formaldehyde

**DOI:** 10.1101/2020.10.16.343392

**Authors:** Jannell V. Bazurto, Dipti D. Nayak, Tomislav Ticak, Milya Davlieva, Jessica A. Lee, Leah B. Lambert, Olivia J. Benski, Caleb J. Quates, Jill L. Johnson, Jagdish Suresh Patel, F. Marty Ytreberg, Yousif Shamoo, Christopher J. Marx

## Abstract

Normal cellular processes give rise to toxic metabolites that cells must mitigate. Formaldehyde is a universal stressor and potent metabolic toxin that is generated in organisms from bacteria to humans. Methylotrophic bacteria such as *Methylorubrum extorquens* face an acute challenge due to their production of formaldehyde as an obligate central intermediate of single-carbon metabolism. Mechanisms to sense and respond to formaldehyde were speculated to exist in methylotrophs for decades but had never been discovered. Here we identify a member of the DUF336 domain family, named *efgA* for enhanced formaldehyde growth, that plays an important role in endogenous formaldehyde stress response in *M. extorquens* PA1 and is found almost exclusively in methylotrophic taxa. Our experimental analyses reveal that EfgA is a formaldehyde sensor that inhibits translation in response to elevated levels of formaldehyde. Heterologous expression of EfgA in *Escherichia coli* increases formaldehyde resistance, indicating that its interaction partners are widespread and conserved and may include translational machinery. EfgA represents the first example of a formaldehyde stress response system that does not involve enzymatic detoxification. Thus, EfgA comprises a unique stress response mechanism in bacteria, whereby a single protein directly senses elevated levels of a toxic intracellular metabolite and modulates translational activity.

## Introduction

Robust organisms employ various mechanisms for averting cellular damage in the face of stress. In all biological systems, routine cellular processes generate highly toxic metabolites that can inflict damage on macromolecules and metabolites. The response systems that mitigate these endogenous stressors vary from detoxification systems, to neutralize reactive compounds such as hydrogen peroxide, imines, and aldehydes [1–4], to post-damage repair systems such as enzyme-mediated damage reversal [5,6] and targeted molecular degradation systems [7]. Induction of these responses can arise due to direct sensing of the metabolic toxins [8] or by sensing the damaged molecules themselves [5].

Formaldehyde is a ubiquitous metabolic toxin generated in most, if not all, organisms as a byproduct of enzymatic reactions or degradation products of metabolites. Due to its high reactivity with amines and thiols in particular, formaldehyde can damage numerous molecules such as metabolites [9], nucleic acids [10], and proteins [11–13]. To date, the only known formaldehyde-specific stress response systems involve enzymatic detoxification [2,14–16]. In bacteria, various formaldehyde detoxification pathways exist, including the widely conserved glutathione- (GSH-) dependent pathway as well as pathways dependent on pterins or sugar phosphates [2]; some species employ multiple pathways [17]. Thus far there is a single example of a formaldehyde sensor, FrmR, a transcriptional repressor that directly binds formaldehyde and controls expression of the detoxification pathway in *Escherichia coli* [18].

Methylotrophs are diverse organisms that can use reduced one-carbon (C_1_) compounds (e.g., methane, methanol) or multi-carbon compounds lacking carbon-carbon bonds (e.g., trimethylamine) as sole sources of carbon and energy. Methylotrophs are of practical importance in cycling C_1_ compounds like methane in the environment, consuming methylated compounds that affect microbiome-gut interactions [19–21], and converting C_1_ substrates to valuable products in industrial settings [22]. Due to their metabolic capacity, methylotrophs face the unique challenge of managing high fluxes of formaldehyde as a central metabolic intermediate.

Even for methylotrophs, where the metabolic pathways for the production and consumption of formaldehyde have been identified, no genes have been identified that allow cells to sense and respond to formaldehyde. The alphaproteobacterium *Methylorubrum extorquens* (formerly *Methylobacterium* [23] is the most extensively studied facultative methylotroph. During growth on methanol, *M. extorquens* uses periplasmic methanol dehydrogenases to oxidize methanol to formaldehyde, which is presumed to diffuse through the inner membrane or possibly undergo active transport [24] (Fig 1). In the cytoplasm formaldehyde condenses with the C_1_ carrier, dephospho-tetrahydromethanopterin (dH_4_MPT) [25,26], a reaction catalyzed by formaldehyde-activating enzyme (Fae, EC: 4.2.1.147) [27]. Through a series of dH_4_MPT intermediates, C_1_ units are oxidized to formate. This branchpoint metabolite is then either further oxidized to CO_2_ or assimilated into biomass via a tetrahydrofolate-linked pathway and, subsequently, the serine cycle [28]. The dH_4_MPT pathway plays the dual role of formaldehyde oxidation for C_1_ growth and detoxification, whereby mutants with a defective dH_4_MPT pathway exhibit methanol sensitivity due to an inability to detoxify formaldehyde produced intracellularly [25]. Indeed, at the level of gene expression, Fae and the enzymes of the dH_4_MPT pathway are present at high levels during growth on either C_1_ or multi-C growth substrates [29–31]. These data have led to the picture that methylotrophs, which can have intracellular fluxes of 100 mM/min [27] and intracellular concentration of ~ 1 mM [32] during growth on methanol, simply have sufficiently high, constitutive level of formaldehyde oxidation capacity to prevent toxic buildup, perhaps obviating formaldehyde sensing and an associated response.

**Figure 1.**
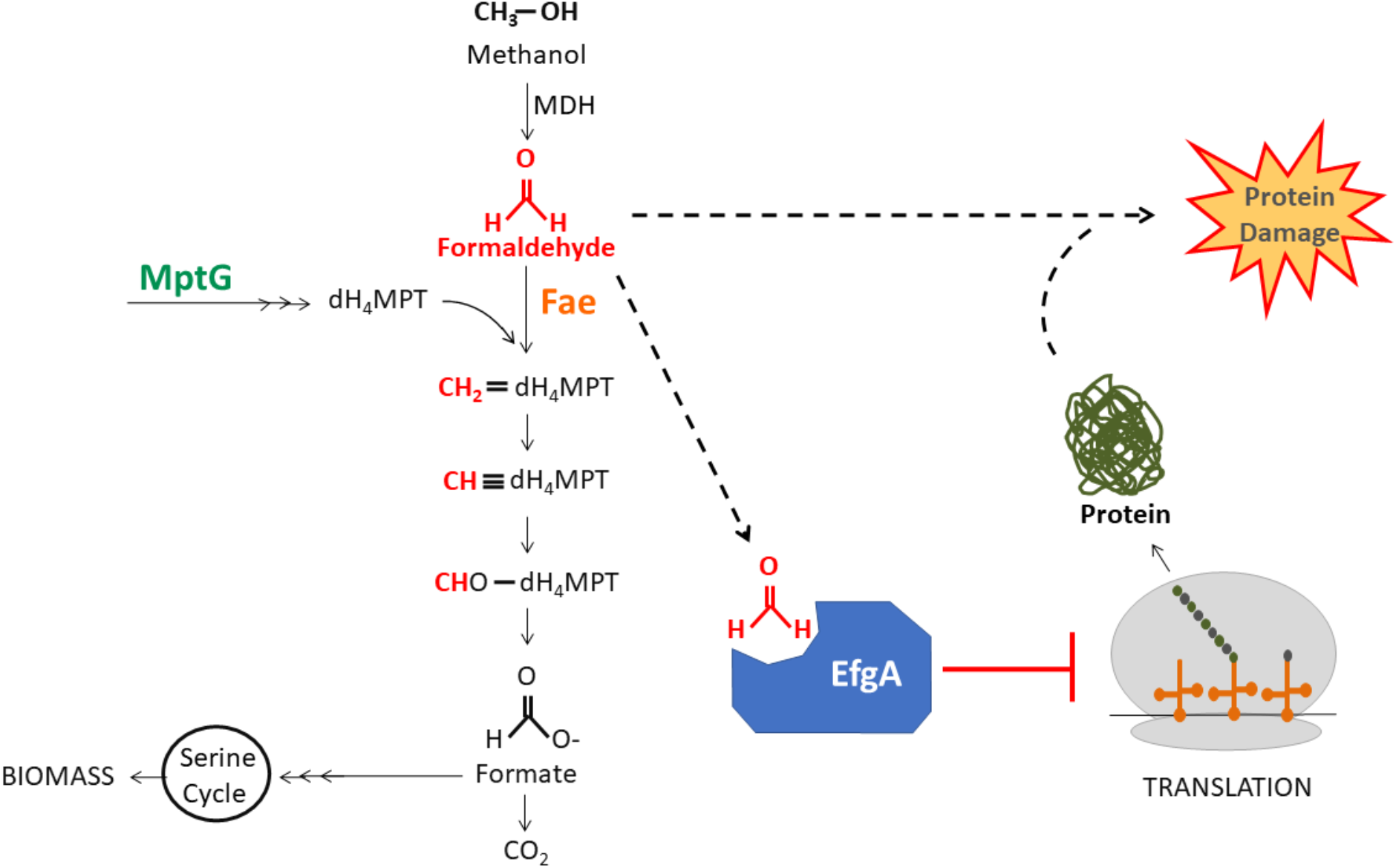
Methanol utilization pathway in *M. extorquens*. Methanol is oxidized to formaldehyde by methanol dehydrogenase (MDH). Fae then condenses free formaldehyde and dephosphotetrahydromethanopterin (dH_4_MPT). MptG is required for dH_4_MPT biosynthesis. The pathway branches at formate which can be further oxidized to CO_2_ or routed to the assimilation pathways (e.g., serine cycle). Alternatively, free formaldehyde can bind EfgA and lead to cessation of translation. Our working model is that EfgA prevents formaldehyde-induced protein damage.

Recent work demonstrating that formaldehyde stress tolerance is responsive to the environment suggests that cells can sense this toxic intermediate. Formaldehyde tolerance was found to be phenotypically heterogeneous across genetically identical individuals of *M. extorquens* PA1, whereby some cells could not even survive exposure to 1 mM formaldehyde when it was provided in the growth medium [33]. This is consistent with the paradoxical finding that many methylotrophs are unable to directly grow on formaldehyde even though they rapidly generate it during growth on more reduced C_1_ compounds [25,33]. Although some of the cells in populations were surprisingly sensitive to formaldehyde, other rare cells could grow at normal rates under conditions where the vast majority rapidly die. The distribution of formaldehyde tolerances was found to rapidly shift upwards in response to formaldehyde stress and the distribution would relax downwards in its absence. Critically, differential death and growth could not explain the distribution shifts, arguing that cells have yet undiscovered systems to sense and respond to formaldehyde and do not rely solely upon consistently high levels of dH_4_MPT pathway enzymes for resistance.

Here we have employed experimental evolution to select for growth on formaldehyde and have uncovered multiple loci encoding genes that can impact formaldehyde resistance. Experimental evolution has unique advantages compared to the classical approach of examining mutants defective in a process [34,35], including the ability to invoke the role of essential genes and gain-of-function mutations, both of which we observed herein. The simplest mechanistic hypothesis for increased resistance would be that – like antibiotic resistance mediated by enzymatic modification – evolved resistance would be mediated by an increase in formaldehyde oxidation. Thus, it came as a surprise that none of the loci with beneficial mutations are known to be related to formaldehyde oxidation or any other known methylotrophy gene [36]. Instead, we identified several novel loci, most commonly a gene of unknown function that we name *efgA* (enhanced formaldehyde growth). We explored the role and function of EfgA through a combination of X-ray crystallography, molecular modeling, mutational analysis, and biochemical characterization, revealing that EfgA is a sensor that directly binds formaldehyde and leads to an arrest in protein translation. EfgA is beneficial to cells when confronted with elevated levels of internally produced formaldehyde and, through phylogenetic analyses and heterologous expression, we show that EfgA function is broadly conserved in methylotrophs. Furthermore, EfgA-mediated formaldehyde protection is transferable to non-methylotrophs. Our findings represent the first characterized formaldehyde stress response in methylotrophs and demonstrate a unique strategy where a single protein senses a toxic metabolite and leads to inhibition of protein translation.

## Materials and methods

### Bacterial strains, media, and chemicals

*Methylorubrum* (reclassified from *Methylobacterium* [23]) strains used in this study are derived from *M. extorquens* PA1 [37,38] lacking cellulose synthesis genes to optimize liquid growth measurements [39]. Thus, the genotype referred to herein as ‘wild-type’ (CM2730) is more accurately ΔcelABC.

*Escherichia coli* strains used in the physiological studies were derivatives of BW23474 [40] while those used for cloning and protein overexpression were derivatives of *E. coli* NEB 10-beta and BL21 (DE3) (Stratagene), respectively.

All strains used in this study are described in Tables 1 and S1.

**Table 1.**
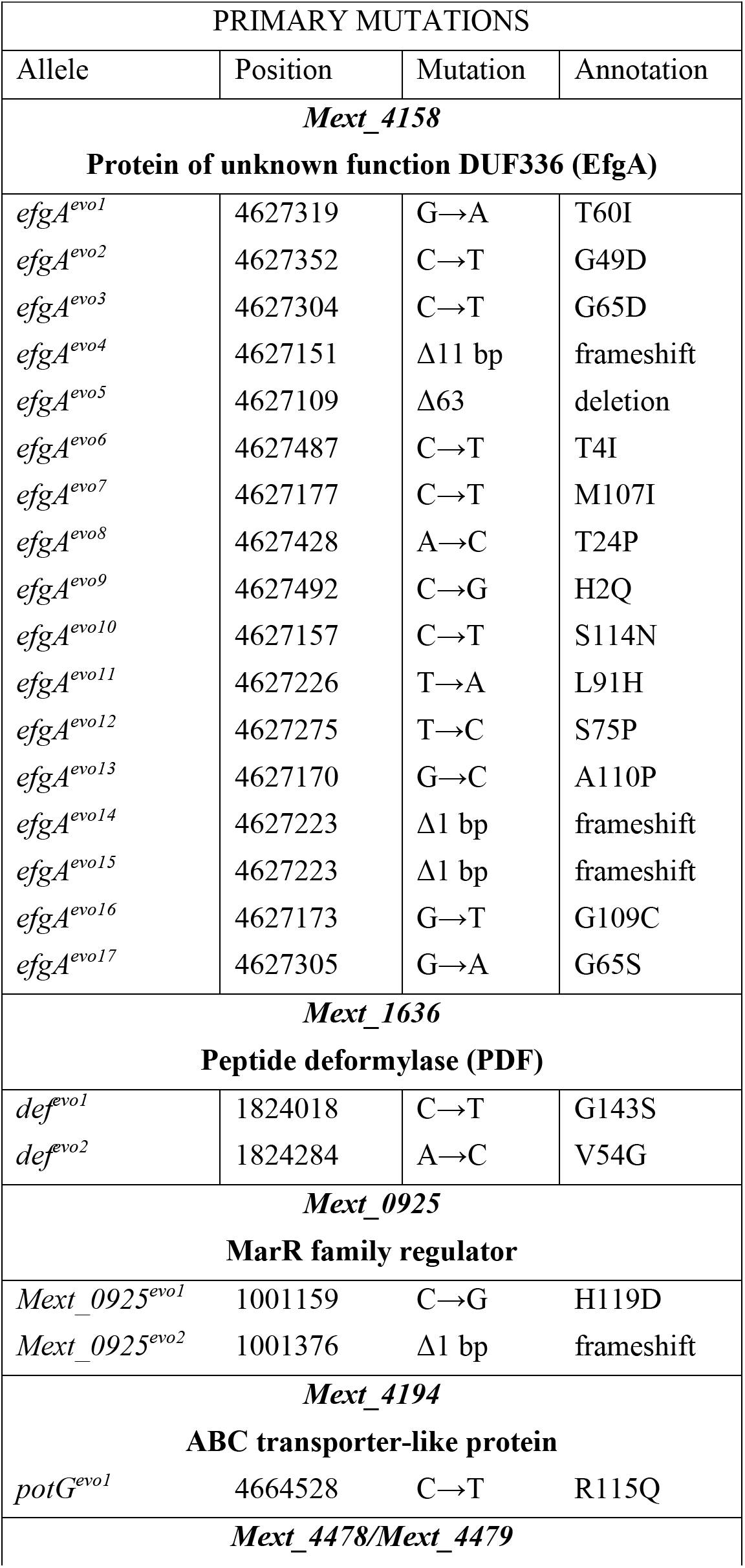
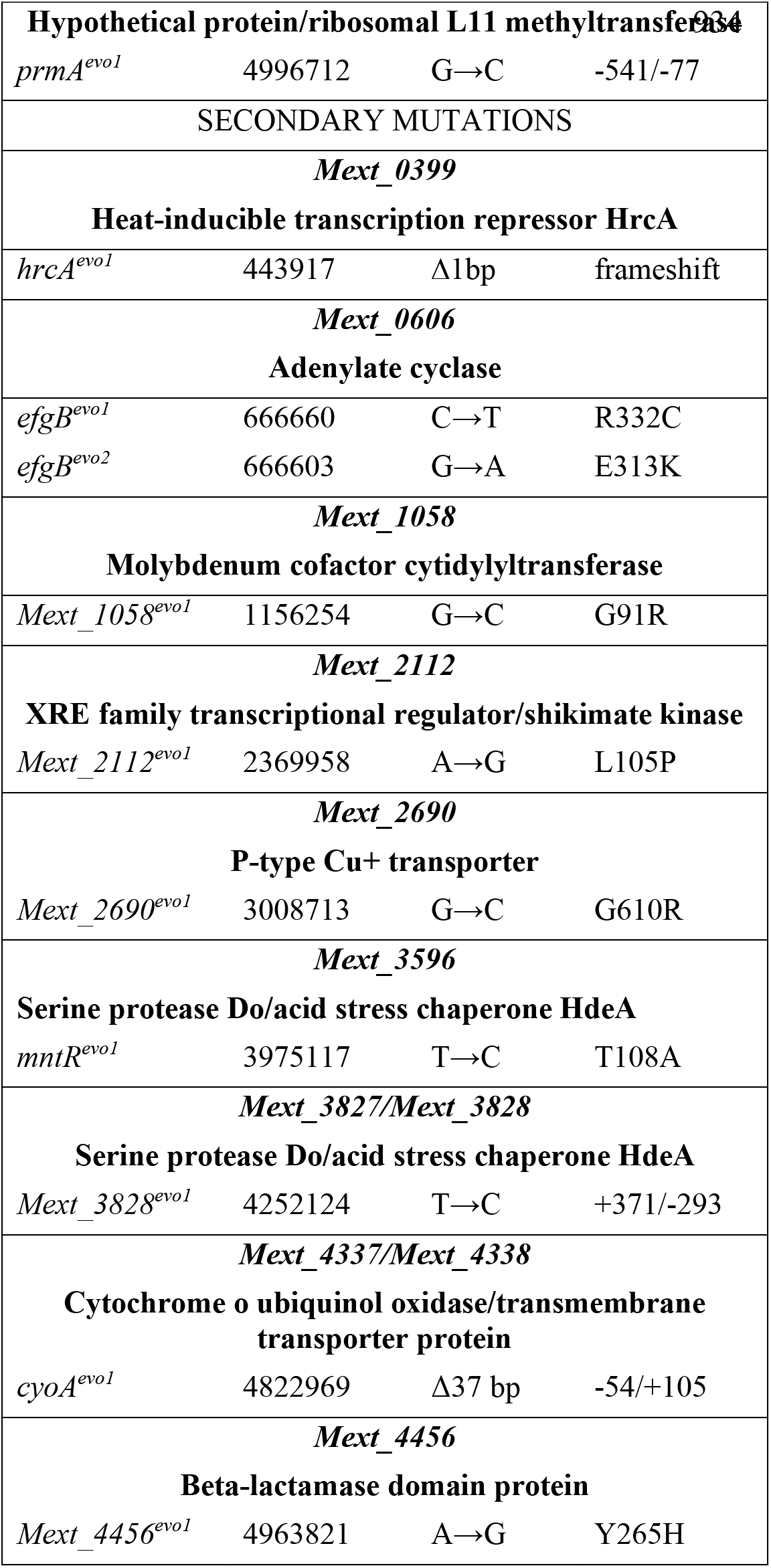
Evolved beneficial alleles

Growth experiments for *M. extorquens* were performed in a modified Hypho medium [41] or *Methylobacterium* PIPES (MP) medium [39] with 3.5 mM succinate, 15 mM methanol, or 2, 4, 5, 6, 8, 10 mM formaldehyde as a sole carbon source. For growth on solid medium, Bacto Agar (15 g/L, BD Diagnostics) was added and the concentrations of succinate or methanol were increased to 15 or 125 mM, respectively, or BD Difco nutrient agar was used.

Formaldehyde stock solutions (1 M) were prepared by boiling 0.3 g paraformaldehyde and 10 mL of 0.05 N NaOH in a sealed tube for 20 m stocks were kept at room temperature and made fresh weekly (growth experiments) or daily (*in vitro* binding experiments). When present in the media, compounds were at the following final concentrations: kanamycin (50 μg/mL); tetracycline (12.5 μg/mL), trimethoprim (10 μg/mL), streptomycin (100 μg/mL), sucrose (50 g/L), cumate (30 μg/mL). Stock solutions of glyoxal, acetaldehyde, butyraldehyde, glutaraldehyde, propionaldehyde were prepared in water or ethanol.

Growth experiments for *E. coli* were performed in MOPS medium [42] with 2 mM glucose, 0.7-1.1 mM formaldehyde, and 0.5 mM L-rhamnose for induction.

Chemicals were purchased from Sigma Aldrich, VWR or ThermoFisher Scientific.

### Growth analyses

Starter cultures of *M. extorquens* (2 mL) were growth in biological triplicate by inoculating media with individual colonies. Cultures were grown in Hypho or MP liquid medium with shaking (250 rpm on platform shaker or 70 rpm in a New Brunswick TC-7 culture roller drum) during incubation at 30 °C. Early stationary-phase cultures (24 h for succinate, 36 h for methanol) were then subcultured (1/64) into relevant media for growth measurements. Cell density was determined by monitoring absorbance with a Spectronic 200 (Thermo Scientific) or a SmartSpec Plus (Bio-Rad) at 600 nm. To determine cell viability (CFU/mL), a 100 μL aliquot of culture was used to harvest cells by centrifugation. The supernatant was discarded and the cell pellet was resuspended into MP medium (no carbon). Cell suspensions were then serially diluted (1/10 dilutions, 200 μL total volume) in 96-well polystyrene plates with MP medium (no carbon) and 10 μL aliquots of each dilution were spotted to MP medium plates (15 mM succinate) in technical triplicate. Plates were inverted and incubated at 30 °C until colony formation was apparent (4-6 d), at which point colonies were counted. Technical triplicates were averaged for each biological replicates and biological replicates were averaged.

Starter cultures for growth analyses of *E. coli* were initiated from freezer stocks of WM8637 or WM8653 into 5 mL tubes at 37 °C shaken at 250 rpm. After growth to stationary phase overnight, these were subcultured (1/500) into MOPS medium with 2 mM glucose. After overnight growth, both cultures were diluted to an OD_600_ = 0.02 into MOPS with 2 mM glucose with or without 0.5 mM rhamnose. A volume of 640 μL was pipetted into Costar 3548 48-well plates (Corning) and grown without lids at 37 °C in Synergy H1 plate readers (BioTek) with double-orbital shaking at 425 rpm and a 3 mm orbit and readings taken every 15 minutes (OD_600_ with 100 ms delay and 8 measurements per data point). After 2.33 h to establish exponential growth, the dispenser of the Synergy H1 was used to automatically deliver formaldehyde at 225 μL/sec from a 32 mM stock made in H_2_O (volume added ranged from 14-22 μL) straight into the wells to final concentrations of 0.7, 0.9, or 1.1 mM. Data were analyzed by first subtracting the average of the blank wells used at the corners of the plate. The data shown are from non-edge wells on the plate; consistent trends with shifted times of recovery were found for the daily duplicates in the edge wells. The entire experiment was repeated on three additional days (each with duplicates) from separate starter cultures.

### Genomic context of *efgA* and *efgB* in other organisms

To examine synteny of *efgA* and *efgB* the genomic context of each gene was examined in organisms with closely related homologs using GeneHood (https://gitlab.com/genehood/genehood-cli) and the MiST3 (Microbial Signal Transduction Database) [43]. Figures were generated using the GeneHood software.

### Experimental evolution

From individual colonies, three independent cultures of CM2730 were grown in 10 mL liquid Hypho medium in batch culture in 50 mL flasks (sealed) supplemented with 15 mM methanol. Upon reaching stationary phase, a 156 μL aliquot (1/32 inoculum) was transferred to fresh medium containing decreased methanol and increased formaldehyde concentrations (Fig S1). For the first 60 generations the concentration of formaldehyde increased with each transfer. For the next 90 generations the concentration of formaldehyde was kept constant (20 mM). Final populations (150 generations) were plated to a series of solid media containing: i) nutrient agar, 15 mM methanol, 3.5 mM succinate, 15 mM formate, 5 mM formaldehyde, ii) Hypho, 4 mM succinate, iii) Hypho, 100 mM methanol, and iv) Hypho, 30 mM formate. Colonies that arose were streaked for isolation on the same respective medium and were further characterized.

A second round of experimental evolution used 25 individual colonies and only involved four transfers after an initial round of growth at 15 mM methanol: i) 10 mM methanol, 1 mM formaldehyde, ii) 5 mM methanol, 2.5 mM formaldehyde, iii) 5 mM methanol, 5 mM formaldehyde, and iv) 5mM formaldehyde.

### Sequence acquisition and phylogenetics

#### Maximum likelihood phylogenetic tree of EfgA

The amino acid sequence of EfgA was used as a query with PSI-BLAST [44] using an 1E^−10^ cutoff in RefSeq [45]. From each of the 5172 matches, the DUF336 sequence was extracted for analysis; sequences > 30% identity were analyzed with CD-HIT [46], allowing us to eliminate all sequences with 90% or greater identity. The average amino acid size was then calculated for the database, and any sequence longer 1σ was removed prior to alignment. The formaldehyde-bound crystal structure of EfgA 6C0Z, was used as a query for homologous structures, which were then added into the sequence database manually. The 5172 sequences were then aligned with MUSCLE [47] using default parameters. The alignment file was then analyzed with FastTree 2.1 [48] with the LG + CAT [49], WAG + CAT [50], and JTT + CAT [51] models with and without gamma distribution. The LG + CAT model generated the best LogLk (−584712.707) and bad splits (15/5168). The phylogenetic reconstruction was then analyzed and annotated with iTOL [52] with marker positions for known structures added to the tree. The final phylogenetic data is available at TreeBASE (http://purl.org/phylo/treebase/phylows/study/TB2:S27073).

#### Maximum likelihood phylogenetic tree of EfgB

The nucleotide sequence of EfgB was used as a query with BLASTN [44] using a filter to eliminate all sequences with 90% or greater identity; the top 100 matches (all >65% identity) were used to assess phylogeny. The evolutionary history was inferred by using the maximum likelihood method and the general time reversible model [53]. The bootstrap consensus tree was inferred from 500 replicates [54]. The percentage of replicate trees in which the associated taxa clustered together in the bootstrap test (500 replicates) are shown next to the branches [54]. Initial tree(s) for the heuristic search were obtained automatically by applying Neighbor-Join and BioNJ algorithms to a matrix of pairwise distances estimated using the Maximum Composite Likelihood (MCL) approach, and then selecting the topology with superior log likelihood value. This analysis involved 97 nucleotide sequences. There were a total of 1578 positions in the final dataset. Evolutionary analyses were conducted in MEGA X [55]. The final phylogenetic data is available at TreeBASE (http://purl.org/phylo/treebase/phylows/study/TB2:S27073).

### Genetic approaches in *M. extorquens* and *E. coli*

Allelic exchange was used to introduce changes into relevant genetic loci as previously described [56]. Due to challenges with identifying a tetracycline concentration that would reliably differentiate between strains with and without the resistance marker in *M. extorquens* PA1, we constructed a kanamycin resistant version of the allelic exchange vector. A 2 kb region of pCM433 [56] encoding cat and most of *tet* was excised using *Eco*RV. The remaining vector backbone, with the exception of a 0.9 kb region containing *bla* was PCR amplified and joined via Gibson assembly (HiFi DNA Assembly, New England Biolabs) with a 1.3 kb PCR product containing *kan* PCR amplified from pCM66 [57] to generate pPS04 (Fig S2). The complete sequence of pPS04 is available in GenBank (Submission 2392028, pending) and has been deposited at Addgene (pending).

Deletions of *efgA*, eliminated 404 bp from the ORF of *_4158* (21-424/435); deletions of *efgB, fmt*, and *fmt-def* eliminated the entire coding region(s) of *_0606* (1401 bp), *_1635* (930 bp), and *_1635-_1636* (1456 bp), respectively.

Inducible expression vectors derived from pLC290 were used to express *efgA* from *M. extorquens* PA1 and *Mfla_1444* from *Methylobacillus flagellatus* KT in *M. extorquens* (pDN147 and pDN162, respectively). Vectors include 30 bp upstream sequence of each gene including respective native ribosomal binding sites.

The WM8655 strain was generated to express *efgA* from *M. extorquens* PA1 from the rhamnose-inducible P*_rhaS_* promoter in *E. coli*. The *efgA* coding sequence plus 30 bp at the 5’ end was amplified using primers with 30 nt overlaps to permit Gibson assembly (HiFi DNA Assembly, New England Biolabs) into pAH120 [40] that had been digested with *XbaI* and *NdeI*, generating pDN380. An empty control strain, WM8637, was first generated by electroporation of pINT-ts into BW23474 [40]. pDN380 was introduced via electroporation into WM8637 to generate WM8655. Constructs were confirmed by analytical PCR and sequencing of the *efgA* locus.

All vectors were designed using SnapGene software. The Gibson assembly kit from New England Biolabs was used to construct vectors from restriction enzyme-digested, linearized vector backbone and PCR-generated inserts. For *E. coli*, transformations were performed using standard (WM8637 and WM8655) or manufacturer’s (BL21 (DE3)) protocols for chemical transformation. For complementation of *M. extorquens*, triparental conjugations were performed using pRK2073 [58].

### Formaldehyde quantification

Formaldehyde concentrations in the culture media were measured as previously described [59]. Supernatant from a 100 μL aliquot of culture was isolated by centrifugation (14,000 x g). In technical triplicate, 10 μL of the supernatant or 100 μL of 0.1X supernatant (diluted with MP medium, no carbon) was combined with 190 or 100 μL Nash reagent B (2 M ammonium acetate, 50 mM glacial acetic acid, 20 mM acetylacetone), respectively, in 96-well polystyrene plates. Reaction plates were incubated (60 °C, 10 min), cooled to room temp (5 min) and absorbance was read at 432 nm on a Wallac 1420 VICTOR Multilabel reader (Perkin Elmer). Formaldehyde standards were prepared daily from 1 M formaldehyde stock solutions and a standard curve was alongside all sample measurements.

### Crystallization and structure of *M. extorquens* EfgA in complex with formaldehyde

#### Expression and purification of EfgA from *M. extorquens*

The gene encoding EfgA (including the thrombin cleavable 6XHis tag) was transformed into *E. coli* strain BL21 (DE3) (Stratagene) for overexpression from pDN79. The protein was overexpressed by growing cells in Luria Bertani broth (LB) medium to an A_600_ of 0.6 at 37 °C and subsequent induction with 0.5 mM IPTG for 14 h at 16 °C. The cell pellet was collected by centrifugation and resuspended in buffer A (50 mM Tris pH 7.5, 1 M NaCl, 20 mM imidazole, 0.3 mM DTT, 0.2 mM PMSF and Complete protease inhibitor cocktail tablet, EDTA-free (Roche Diagnostics Corp, Indianapolis, IN, US)). Lysis was performed by sonication. The lysate was then centrifuged for 60 m at 24,000 rpm at 4 °C. The supernatant was then applied onto a Hi Trap affinity (5 ml) (Ni^2+^) column (GE Healthcare Life Sciences) The column was washed with 10 column volumes of buffer A, and the protein was then eluted with increasing concentrations of imidazole from 20 to 500 mM. The fractions containing the protein of interest were pooled, dialyzed against 50 mM Tris pH 7.5, 0.3 mM DTT, 0.5 mM EDTA and purified over a Q-XL Sepharose column (GE Healthcare) using a 0.1-1 M NaCl gradient. The fractions containing EfgA were pooled and concentrated in an Amicon centrifugal filter concentrator with a 10 kDa cutoff membrane (Millipore). The concentrated protein was then further purified by size-exclusion chromatography using a Superdex-200 column (GE Healthcare, HiLoad 16/60) equilibrated with buffer (50 mM Tris pH 8.0, 250 mM NaCl). Again, the fractions containing the protein were pooled and concentrated with an Amicon centrifugal filter concentrator with a 10 kDa cutoff membrane. The purity of the protein was analyzed with 15% SDS–PAGE by using ImageJ software and was determined to be greater than 95%.

#### Crystallization and structure

Crystals of *M. extorquens* EfgA were obtained by sparse matrix screening at 15 mg/mL at 4 °C and 10 °C. Preliminary results were followed by optimization of the successful condition manually using the sitting drop vapor diffusion method. The best quality crystals were grown in 0.2 M potassium fluoride, 2.2 M ammonium sulfate ((NH_4_)_2_SO_4_) at 10 °C. The EfgA-formaldehyde crystals suitable for data collection were grown at 10 °C in 0.2 M KNO3, 2.2 M (NH_4_)_2_SO_4_, 16.6 mM formaldehyde using the hanging drop vapor diffusion method. Diffraction data sets for *M. extorquens* EfgA were collected at 100 K at 1.65 Å resolution at the Argonne National Laboratory’s Advanced Photon Source (ANL APS) beamline 21-ID-G on a MarMosaic 300 CCD detector. Diffraction data for EfgA in complex with formaldehyde were collected at 100 K at 1.85 Å resolution at Argonne National Laboratory’s Advanced Photon Source (ANL APS) beamline 21-ID-D on a MarMosaic 300 CCD detector. X-ray diffraction data were processed using HKL2000 [60]. The crystal of apo-protein belonged to the space group P22121 with the unit cell parameters a=71.82, b=72, c=105.18, α=β=γ=90°. *M. extorquens* EfgA in complex with formaldehyde crystallized in space group P21221 with the unit cell parameters a=72.06, b=72.06, c=104.56, α=β=γ=90°. The 3-dimensional structures were determined by molecular replacement using *Klebsiella pneumoniae* protein OrfY (Pfam DUF336) domain as the search model in Phaser-MR [61]. The molecular replacement for EfgA-formaldehyde complex was further confirmed by the initial (2Fo-Fc) map generated using Coot [62] that clearly indicated electron density for the formaldehyde that was not included in the original search model. The structure was refined using the Phenix suite [63] and Coot [62]. Ramachandran plots and root-mean-square deviations (rmsd) from ideality for bond angles and lengths were determined using a structure validation program, MolProbity [64]. A summary of data collection and refinement statistics are listed in Table 2.

**Table 2.**
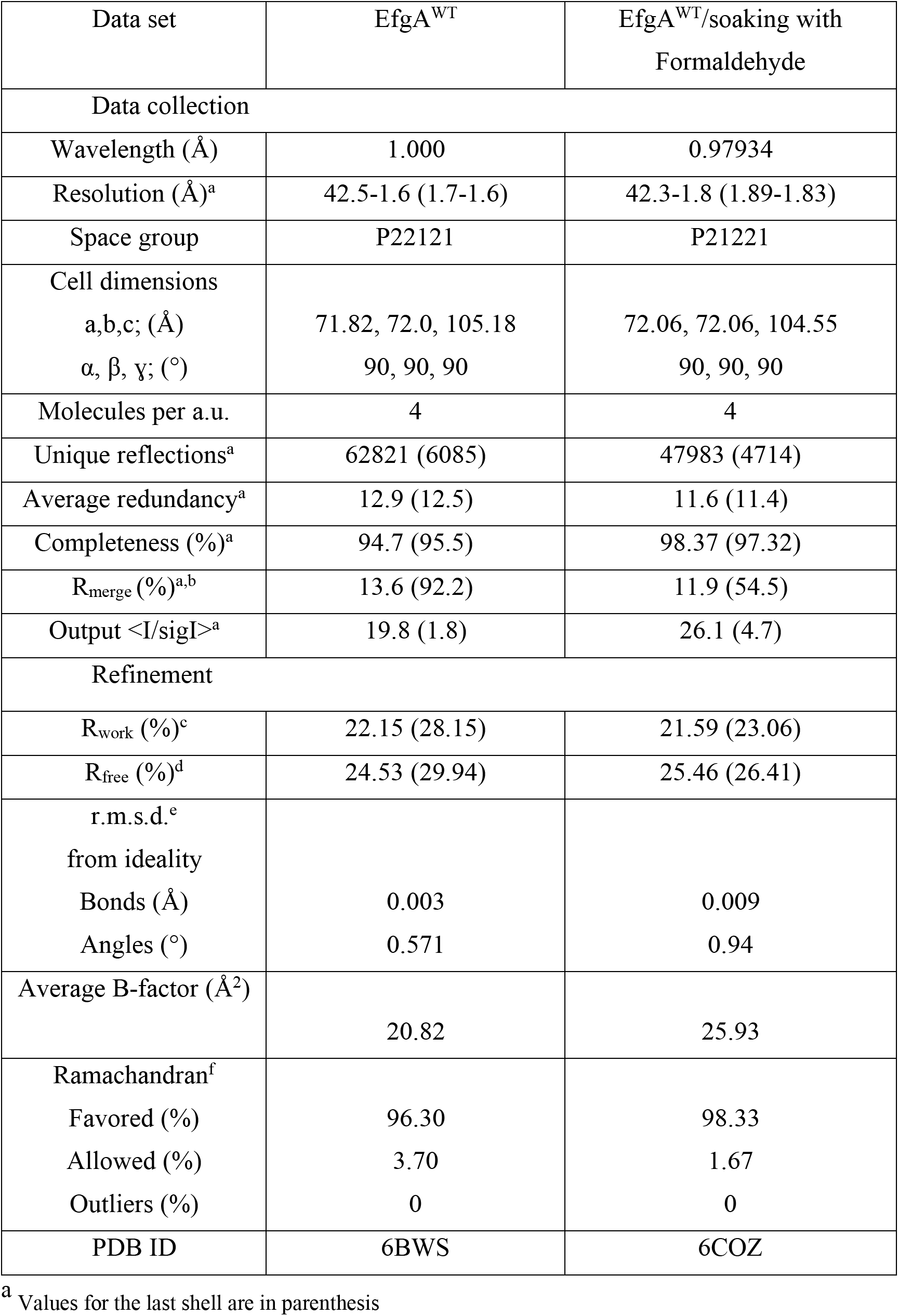
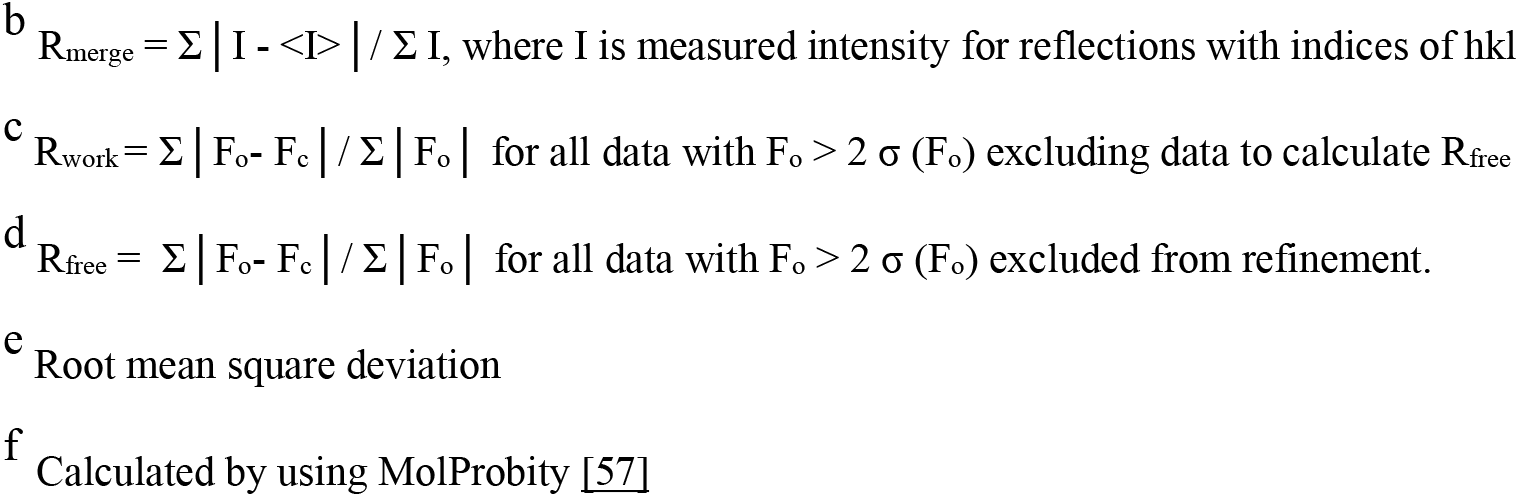
Data collection and refinement statistics

| Data set | EfgA<sup>WT</sup> | EfgA<sup>WT</sup>/soaking with Formaldehyde |
| --- | --- | --- |
| Data collection |  |  |
| Wavelength (Å) | 1.000 | 0.97934 |
| Resolution (Å)<sup>a</sup> | 42.5-1.6 (1.7-1.6) | 42.3-1.8 (1.89-1.83) |
| Space group | P22121 | P21221 |
| Cell dimensions |  |  |
| a,b,c; (Å) | 71.82, 72.0, 105.18 | 72.06, 72.06, 104.55 |
| α, β, γ; (°) | 90, 90, 90 | 90, 90, 90 |
| Molecules per a.u. | 4 | 4 |
| Unique reflections<sup>a</sup> | 62821 (6085) | 47983 (4714) |
| Average redundancy<sup>a</sup> | 12.9 (12.5) | 11.6 (11.4) |
| Completeness (%)<sup>a</sup> | 94.7 (95.5) | 98.37 (97.32) |
| R<sub>merge</sub> (%)<sup>a,b</sup> | 13.6 (92.2) | 11.9 (54.5) |
| Output <I/sigI><sup>a</sup> | 19.8 (1.8) | 26.1 (4.7) |
| Refinement |  |  |
| R<sub>work</sub> (%)<sup>c</sup> | 22.15 (28.15) | 21.59 (23.06) |
| R<sub>free</sub> (%)<sup>d</sup> | 24.53 (29.94) | 25.46 (26.41) |
| r.m.s.d.<sup>e</sup><br>from ideality |  |  |
| Bonds (Å) | 0.003 | 0.009 |
| Angles (°) | 0.571 | 0.94 |
| Average B-factor (Å<sup>2</sup>) | 20.82 | 25.93 |
| Ramachandran<sup>f</sup> |  |  |
| Favored (%) | 96.30 | 98.33 |
| Allowed (%) | 3.70 | 1.67 |
| Outliers (%) | 0 | 0 |
| PDB ID | 6BWS | 6COZ |

<sup>a</sup> Values for the last shell are in parenthesis
<sup>b</sup> $R_{\text{merge}} = \Sigma |I - \langle I \rangle| / \Sigma I$ , where $I$ is measured intensity for reflections with indices of $hkl$
<sup>c</sup> $R_{\text{work}} = \Sigma |F_o - F_c| / \Sigma |F_o|$ for all data with $F_o > 2 \sigma(F_o)$ excluding data to calculate $R_{\text{free}}$
<sup>d</sup> $R_{\text{free}} = \Sigma |F_o - F_c| / \Sigma |F_o|$ for all data with $F_o > 2 \sigma(F_o)$ excluded from refinement.
<sup>e</sup> Root mean square deviation
<sup>f</sup> Calculated by using MolProbity [57]

### *In silico* predictions of formaldehyde binding site and folding and binding stabilities of EfgA variants

To predict the location of the formaldehyde binding pocket on the EfgA tetramer, and to estimate folding and binding stabilities of EfgA variants, classical molecular dynamics (MD) simulations were carried out. These simulations used the apo EfgA tetramer X-ray crystal structure (PDB ID: 6BWS) and EfgA monomer (chain A of EfgA tetramer) and were performed using GROMACS v2018 [65]. The AutoDock Vina program [66] was then used to dock formaldehyde to snapshots of the EfgA tetramer for 100 snapshots extracted from the MD simulations to determine the pockets most heavily populated with high scoring poses. EfgA monomer and tetramer snapshots were analyzed using FoldX software (MD+FoldX approach) to estimate folding and binding stabilities of EfgA variants.

#### Molecular dynamics simulation of EfgA monomer and tetramer

Both EfgA tetramer and EfgA monomer structures were subjected to atomistic MD simulations using the same protocol. MD simulations were performed using AMBER99SB*-ILDN [67] forcefield. The EfgA structure was placed in a cubic box of TIP3P water and the net charge was neutralized by adding Na^+^ and Cl^−^ ions at a concentration of 0.15 M. Protonation states for all ionizable residues were automatically assigned for neutral pH. The system was then minimized using the steepest descent algorithm for 10,000 steps. The subsequent equilibration process was to perform 1 ns simulation with the positions of all heavy atoms in the complex harmonically restrained to allow equilibration of the water molecules around the proteins, followed by another 1 ns simulation with no restraints. During equilibration, the temperature and the pressure of the system was set to 300 °K and 1 atm respectively using the Berendsen algorithm [68]. Production simulations were then carried out for 100 ns with pressure maintained using Parrinello-Rahman barostat [69] and temperature was controlled using the v-rescale thermostat [70]. Particle mesh Ewald [71] was used to treat electrostatics with a real-space cutoff of 1.2 nm. Van der Waals interactions were cut off at 1.2 nm with the Potential-shift-Verlet method for smoothing interactions. The LINCS algorithm [72] was applied to constrain all bonds to their ideal lengths and timestep of 2 fs was used. During the 100 ns production simulation snapshots were saved every 1 ns giving 100 snapshots of EfgA tetramer and EfgA monomer to be used for docking calculations and FoldX analysis.

#### Docking of formaldehyde to EfgA tetramer

Each of the 100 EfgA tetramer snapshots obtained during MD simulations was used to dock formaldehyde with the AutoDock Vina software [66]. Generally, docking programs allow the ligand to be completely flexible during the conformational search with only a few restricted side chains on the protein assigned as flexible. Use of 100 snapshots from MD simulation allows us to at least partly overcome this limitation.

The 3-D coordinates of formaldehyde were obtained from PubChem (https://pubchem.ncbi.nlm.nih.gov) and assigned Gasteiger partial charges using Autodock Tools (http://mgltools.scripps.edu/). Ligand docking was then carried out by creating a grid box of size 80 Å × 60 Å × 60 Å, centered on the geometric center of the EfgA tetramer, with a grid spacing of 1 Å. All regions of the tetramer protein complex were included in the search for the most favorable interactions of the ligand. The input exhaustiveness parameter for the docking was set to 400. The number of top docking orientations with high docking scores was fixed to 20. This docking protocol was applied to all 100 snapshots and the X-ray crystal structure of the EfgA tetramer, yielded 2020 (101 snapshots × 20 top docking conformations) conformations of formaldehyde bound to the EfgA tetramer. Highly populated docking clusters were then identified using VolMap plugin built in VMD software [73].

#### Predicting folding and binding stabilities of EfgA variants

A mutation of EfgA can affect the folding of a monomer and/or the formation of the tetramer. In order to determine how amino acid mutations, including the known variants, alter stabilities (ΔΔG values) for EfgA folding and formation of a tetramer we calculated ΔΔG values of folding and binding using our previously successful MD+FoldX approach [74–76]. This involves analyzing MD snapshots with FoldX software [77]. MD snapshots of the EfgA monomer and tetramer were analyzed using the same protocol reported in our previous study [76]. Briefly, each snapshot was subjected to the RepairPDB command six times in succession to minimize and obtain convergence of the potential energy. For each snapshot, all possible 19 single mutations in the monomer/tetramer at each amino acid site were then generated using BuildModel command. Lastly, the folding stability of the EfgA monomer due to each mutation was estimated using Stability command, and the binding stability of the EfgA tetramer was estimated using AnalyseComplex command. For each mutation, we then estimated ΔΔG of folding and binding by averaging the FoldX results across all individual snapshot estimates. This process led to a total of 2,546 (134 EfgA residues × 19 possible mutations at each site) ΔΔG values for both folding and binding.

### Comparison of structural homologs

Structures for EfgA homologs (OrfY – 2A2L, HbpS – 3FPV, Ybr137w – 4CLC, DESPIG_02683 – 4NKP, PduOC – 5CX7, and EfgA – 6BWS and 6C0Z) were acquired from PDB (https://www.rcsb.org/) and aligned with PyMol v2.3 [78].

### EfgA:ligand binding assays

#### Expression and purification of EfgA

Recombinant EfgA containing a C-terminal 6X-His tag was expressed in *E. coli* BL21(DE3) housing pET28 vectors (Novagen). Single colonies from LB supplemented with 50 μg/mL kanamycin were used to generate 25 mL overnight cultures grown at 37 °C with continuous shaking at 250 RPM. Cultures were diluted to an OD_600_ = 0.05 in a 1 L flask containing 800 mL of LB kanamycin then placed at 37 °C. When cultures reached OD_600_ ~ 0.52-0.60, they were induced with 1 mM isopropyl β-D-1-thiogalactopyranoside (IPTG) and grown for 4 hours at 37 °C. Cells were then harvested by centrifugation at 10,000 x g for 20 m at 4 °C. Harvested cells were washed with Buffer A (50 mM Na_2_HPO_4_, 300 mM NaCl, 35 mM imidazole at pH 8.0), then harvested again and resuspended in 15 mL Buffer A. Cells were lysed via French Pressed (Thermo Fisher Scientific) in a pre-chilled cell at 20,000 psi. Cell lysate was cleared by centrifugation at 50,000 x g for 2 hours at 4 °C followed by filtration through 0.22 μm PTFE filters and stored at –80 °C prior to purification.

Protein lysates were thawed at 4 °C and centrifuged briefly to ensure no precipitants were present prior to column chromatography. An NGC FPLC (Bio-Rad) was used to purify the C-terminal 6X-His EfgA with 1 mL Ni-NTA columns (Bio-Rad). Columns were equilibrated with 10 mL of Buffer A at 1 mL/min prior to loading lysates. Lysates were loaded at 0.35 mL/min followed by 10 mL of Buffer A at 1 mL/m. An isocratic phase was generated by passing 10 mL of Buffer B (50 mM Na_2_HPO_4_, 300 mM NaCl, 500 mM imidazole at pH 8.0) gradually through the column in reverse phase at 0.5 mL/min with samples being collected every 0.5 mL. Degassed H_2_O (10 mL) was run at 1.0 mL/min until conductivity and absorbance (280 nm) were zero. Samples with high 280 nm values were collected during the isocratic phase of Buffer B.

Fractions from FPLC purification were treated with Laemmli buffer [79] with 10 mM 1,4-dithiothreitol (DTT) and heated for 5 m at 85 °C then analyzed on a 4-15% discontinuous sodium dodecyl sulfate polyacrylamide gel electrophoresis (SDS-PAGE) gel with a 6% stacking gel run at ambient temperature at a constant 100 V. Samples that showed high purity were pooled and stored with 9% glycerol (v/v) and then quantitated with a Bradford assay [80] prior to storage at – 80 °C.

#### Microscale isothermal calorimetry (mITC)

Binding of formaldehyde and EfgA was measured via mITC using an Affinity ITC - LV (TA Instruments - Waters LLC). An isothermal buffer for experimental titrations of EfgA was designed to minimize binding between ligands and buffer (50 mM Na_2_HPO_4_, 300 mM NaCl, pH 8.0). Pooled samples of EfgA or EfgA variants were buffer exchanged 1:128 fold with mITC buffer using 3K MWCO Amicon cellulose acetate filters at 4000 x g for 30 m in a swinging bucket rotor. Protein samples were quantitated via Bradford assay and normalized to 2 mg/mL using mITC buffer. Initially, protein:ligand concentrations were used in varying ratios to determine working assay parameters. Proteins were then diluted in mITC buffer to a final concentration of 50 μM. Formaldehyde (1 M) was prepared from paraformaldehyde in milliQ-H_2_O and used within 24 hours. Formaldehyde stock was serially diluted to 25 mM in mITC buffer. Methanol, formate, and acetaldehyde were all prepared in the same way as formaldehyde to minimize difference between ligand preparation. Prior to use, protein samples and buffer were degassed for 10 m at 650 mm Hg; ligands were degassed for only 5 m to minimize vapor loss.

All experimental runs were performed with 400 μL mITC buffer in the reference and 400 μL sample cells. Between runs, the sample cell and titration syringe were washed 10 times with degassed H_2_O and 10 times with degassed mITC buffer. The run protocol was 20 injections of 2 μL of 25 mM ligand (with the first being a 0.3 μL throw-away titration) every 200 s with a stir speed of 125 RPM, and 25 °C. Prior to any run or data collection, a slope (μW/h) difference of 0.30 and standard deviation (μW) of 0.03 was required between reference and sample cell.

Isotherms of buffer:ligand were subtracted from protein:ligand data prior to calculation of binding energies. The blank μcal energy was subtracted from the total, which acted as the heat of dilution of ligand into protein.

#### Microscale thermophoresis (MST)

MST experiments were performed on a Monolith NT.115 system (Nanotemper Technologies, San Francisco, CA, USA). A solution of unlabeled formaldehyde was serially diluted in reaction buffer (10 mM Na_2_HPO_4_, 1.8 mM KH_2_PO_4_, 2.7 mM KCl, 137 mM NaCl, 0.05% (v/v) Tween-20) to which an equal volume of Alexa-647 labeled EfgA was added to a final concentration of 20 nM. The samples were loaded into standard treated capillaries (Nanotemper) using 70% LED and 80% IR-laser power. Laser on and off times were set at 30 s and 5 s, respectively. The resulting Kd values are based on an average from three independent MST measurements. Temperature of MST experiments were 20 °C and 30 °C. Data analysis was performed using Nanotemper Analysis software, v.1.5.41.

The raw MST traces for each individual experiment were transformed and fit according to published methods by fitting the normalized fluorescence (Fnorm) to the Hill equation:

(F_norm_=(F_norm,max_-F_norm,min_)x[c_A_]^n^/([c_A_]^n^ + K_d_^n^), where F_norm,max_=maximal normalized fluorescence, F_norm,min_=minimal normalized fluorescence, [c_A_]=concentration of protein, K_d_ = dissociation constant and n=hill coefficient.

### Formaldehyde tolerance distributions

To compare the distribution of formaldehyde tolerance phenotypes among individual cells in populations of WT and Δ*efgA* mutants, formaldehyde tolerance assay plates were prepared as follows: MP medium was prepared with agar, autoclaved, and cooled to 50 °C; then methanol (final concentration: 125 mM) and formaldehyde (to the desired final concentration) were rapidly mixed in, and the agar was poured into 100 mm petri dishes. The dish lids were immediately replaced and plates were cooled on the benchtop. Plates were stored at 4 °C and used within 1 week of pouring. CFU were plated and enumerated as described above.

Cell cultures were grown in MP-methanol medium until stationary phase. They were then subjected to serial 1:10 dilutions in MP to a final dilution of 10^−6^. From each of the seven dilutions, three replicates of 10 μL were pipetted onto each MP-methanol-formaldehyde plate to form spots (total: 21 spots per sample per plate type). The spots were allowed to dry briefly in a laminar flow hood, then lids were replaced and plates were stored in plastic bags and incubated at 30 °C for 4 days before colonies were counted. For each replicate set of seven spots, the two highest-dilution spots with countable colonies were enumerated and summed, then multiplied by 1.1 times the lower of the two dilution factors to calculate the original number of colony-forming units (CFU) in the sample. The frequency of tolerant cells at each formaldehyde concentration was then calculated by expressing the number of CFU at that concentration as a proportion of the CFU measured at 0 mM. For each sample, the mean and standard deviation of the three replicate spot series was calculated. To compare the shape of the curves, we measured the rate of the decline of tolerant cells relative to formaldehyde concentration (log10 cells / mM formaldehyde) by fitting an line to the points where frequencies are consistently statistically different (for WT, the last 4 non-zero values; for Δ*efgA*, the last 4 values).

This method has a limit of detection of 1.65×10^−7^ (an abundance of 34 CFU/mL is necessary to observe 1 cell per 30 μL plated, and the total cell population tested was 2×10^8^ CFU/mL; therefore the least-abundant subpopulation that could be detected, disregarding the effects of Poisson distributions at lower λ, is one with an average frequency of 1.65×10^−7^ within the total population). Although this assay measures the growth of bacterial colonies and not directly that of individual cells, it has been demonstrated to correlate well with single-cell methods of measuring formaldehyde tolerance distribution [33].

### Competition assays

In biological triplicate, strains were acclimated to growth in Hypho supplemented with 15 mM methanol. In each competition, a test strain was mixed 1:1 (by volume) with a fluorescent reference strain expressing mCherry (CM3841, *efgA^evo1^ efgB^evo1^* Δ*hpt::P_tacA_-mCherry*). This mixture was then used to subculture (1/64 inoculum) into 5 mL of identical fresh medium and grown as described above. The frequencies of fluorescent and non-fluorescent cells were quantified at the start (F0, t = 0) and end (F1, t = h when cells reached stationary phase) of the competition experiment using an LSRII flow cytometer (BD, Franklin Lakes, NJ). mCherry was excited at 561 nm and measured at 620/40 nm. For a given sample, at least 1000 cells were counted. An identical set of competitions were set up with 5 mM formaldehyde in place of methanol as the sole carbon source in the growth medium for acclimation and subsequent competition assays. Malthusian fitness values (W) relative to the reference strain were calculated by a previously described equation assuming an average of 64-fold size expansion of mixed populations during competitive growth: W = log(F_1_x64/F_0_)/log((1-F_1_)x64/(1-F_0_)); for a 32-fold size expansion in formaldehyde: W = log(F_1_x32/F_0_)/log((1-F_1_)x32/(1-F_0_)) [41].

### *In vivo* translation assays

Succinate-growth stationary phase cultures of wild-type (CM2730) and the Δ*efgA* mutant (CM3745) were inoculated into MP medium with 15 mM succinate in biological triplicate and grown at 30 °C with shaking. At early exponential phase (OD_600_ = 0.25), 1 mM [^13^CD_3_]-methionine (Met) was added to each culture and mixed to homogeneity. Each culture was divided into three aliquots and treated with i) nothing, ii) 5 mM formaldehyde, or iii) 50 μg/mL kanamycin and immediately returned to incubator. At 0, 20, 40, 60, 90, 180, and 360 min, the optical density was measured and cells from 1 mL of culture were harvested by centrifugation (top speed, 2 m.) for [^13^CD_3_]-Met quantification by modification of a previously described method [81]. For time = 0 m., cells were harvested immediately post [^13^CD_3_]-Met addition, prior to formaldehyde/kanamycin treatment. Cells were washed with 1 mL of MP medium (no carbon) with 5 s vortex and again harvested by centrifugation. Cell pellets were resuspended in 200 μL of 6 M HCl. The suspension was transferred to a new Eppendorf tube, incubated (105 °C, 18 h), and then dried (lid open, 95 °C, ~20 h). Pellets were resuspended in dimethylformamide (DMF) and the suspension was transferred to a new Eppendorf tube. Amino acids were derivatized with the addition of *N*-*tert*butyldimethylsilyl-*N*-methyltrifluoroacetamide with 1% (wt/wt) tertbutyldimethyl chlorosilane (TBDMSTFA), incubated (85 °C, 1 h), and then transferred to GC/MS vials for analysis. GC parameters with minor modifications: pressure, 124.5 psi; total flow, 17.9 mL/min; column flow, 1.35 mL/min; column length, 29 m.

GC/MS analysis showed that the peak area ratio of major peaks 218 m/z (standard Met) and 222 m/z ([^13^CD_3_]-Met) was well-correlated with the presence of each species in a mixture and was thus used to measure [^13^CD_3_]-Met incorporation in cells.

## Results

### Evolution of *Methylorubrum extorquens* on lethal concentrations of formaldehyde identifies two novel loci, *efgA* and *efgB*, with homologs that are stress response sensors

We evolved *M. extorquens* PA1 [38] for robust growth on formaldehyde in order to identify what loci might be involved in sensing and respond to formaldehyde toxicity. The first of these experiments involved using a steady transition from growth on 15 mM methanol to growth on 20 mM formaldehyde in the first 60 generations; this concentration was maintained to generation 150 (Fig S2). Testing growth of individual isolates confirmed growth at these previously lethal concentrations (Fig S3).

Resequencing the genome of a representative isolate from each of the three populations identified nonsynonymous mutations in two genes of unknown function that were both mutated in each lineage, *_4158* and *_0606. _4158* encodes a single domain, 144 amino acid protein with a domain of unknown function (DUF) 336. *_0606* encodes a 466 amino acid protein that encodes a putative adenylate/guanylate cyclase. Neither gene resides in an apparent operon nor are they located near known methylotrophy genes (Fig S4).

Further sequencing in ten additional, randomly picked isolates from each population confirmed the prevalence of nonsynonymous mutations in *_4158* and *_0606* and identified mutations in a few other loci that did not occur in more than one population (Table 1, S1). The repeated occurrence of mutations in *_4158* and *_0606* suggested that these two loci are of particular importance for enhanced formaldehyde growth, thus we renamed them *efgA* and *efgB*, respectively. In a second round of evolution experiments to rapidly obtain growth in 5 mM formaldehyde, isolates from 18 of the 25 populations had mutations in *efgA* (two frameshifts, a 63 bp deletion, the remainder nonsynonymous mutations (Table 1, S1)); no mutations were identified in *efgB*. Subsequent experimentation described below differentiated the respective roles of *efgA* and *efgB* in formaldehyde growth.

### Homologs of EfgA are specifically associated with methylotrophic lineages

Phylogenetic analysis demonstrated that DUF336 domains are encoded in all three domains of life and the representatives with structures are broadly dispersed across the tree (Fig 2A). A few DUF336 family members are encoded in gene clusters of well-characterized metabolic pathways [82–86], and some of these DUF336 proteins are localized within bacterial microcompartments, proteinaceous organelles that can confine catabolic processes that involve the generation of toxic, and often volatile aldehydes [83,84,87]. The role of DUF336 domain in these contexts, however, has remained elusive. The only comprehensive studies on bacterial DUF336 function were with HbpS in the Gram-positive bacterium *Streptomyces reticuli* [88–95]. HbpS localizes to the extracellular face of the cytoplasmic membrane where it senses environmental heme and, in turn, initiates a signaling cascade that mitigates oxidative stress. This raises the possibility that EfgA, despite low levels of sequence similarity to other DUF336 domains (23% identity to HpbS) may also play a role in sensing.

**Figure 2:**
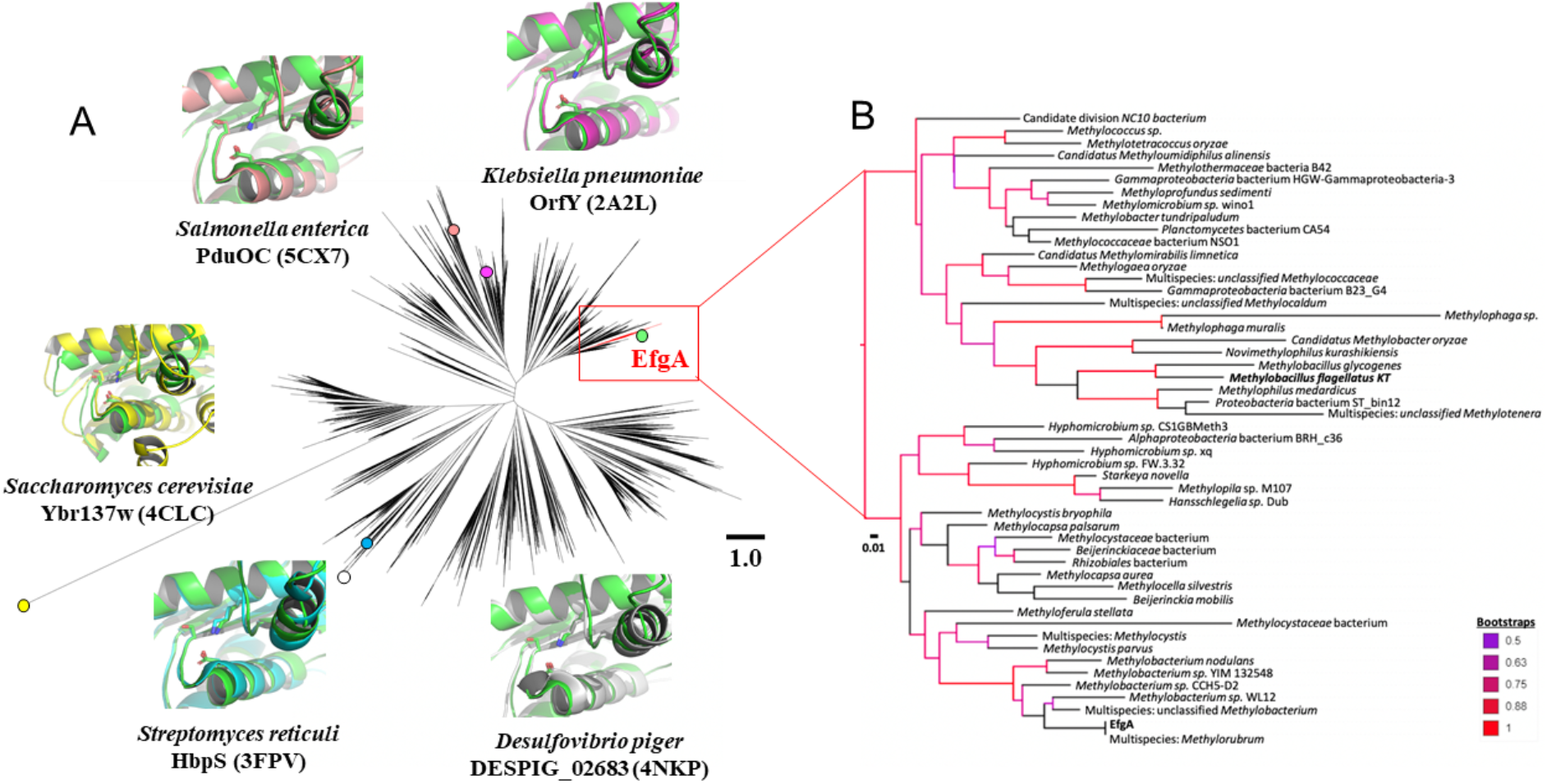
Phylogenetic analysis of the DUF336 superfamily indicates close homologs to EfgA are present in a broad array of methylotrophs. (A) The evolutionary relationship of the DUF336 region was compared to all current amino acid sequences and structures currently on NCBI via maximum likelihood. Solved structures of EfgA homologs are represented on the tree with their respective PDB accession codes and overlaid onto EfgA structure (green). The red colored sequences represent members of the EfgA clade. (B) Expanded view of the EfgA clade with bootstrap values above 50 highlighted by color according to the key. Bolded names indicate experimentally verified EfgA-like function. Scale bars for both (A) and (B) indicate substitutions per residue. The phylogenetic data is available at TreeBASE (http://purl.org/phylo/treebase/phylows/study/TB2:S27073).

Phylogenetic analyses suggest that EfgA is linked to methylotrophy. Focusing upon close homologs of EfgA, we noted that these were found in a highly supported clade whose members are almost exclusively characterized methylotrophs (Fig 2B). These 51 sequences originate from a broad phylogenetic range including *alpha*-, *beta*-, and *gammaproteobacteria*, and NC10 (e.g., *Methylomirabilis oxyfera*) clades. However, we noted that some methylotrophic groups, such as those in the genus *Bacillus*, do not encode an EfgA homolog within this clade.

### Loss of EfgA function necessary and sufficient for growth on formaldehyde

In order to understand how *efgA* is involved in formaldehyde growth, we generated a series of mutants and characterized their phenotype in growth media containing varying concentrations of formaldehyde as the primary carbon and energy source. Introduction of *efgA^evo^* alleles from evolved isolates into the wild-type background was sufficient to enable robust growth in medium containing up to 5 mM formaldehyde (Fig 3, Fig 4). Conversely, restoration of the *efgA*^WT^ allele in an evolved isolate abolished formaldehyde growth, demonstrating the necessity of *efgA^evo^* for formaldehyde growth (Fig 3). Like the evolved alleles, a Δ*efgA* in-frame deletion allele in the wild-type background indicated that the EfgA function must be eliminated to permit growth on formaldehyde. These data, as well as the spectrum of mutations obtained for the *efgA^evo^* alleles indicate that these are loss-of-function mutations.

**Figure 3.**
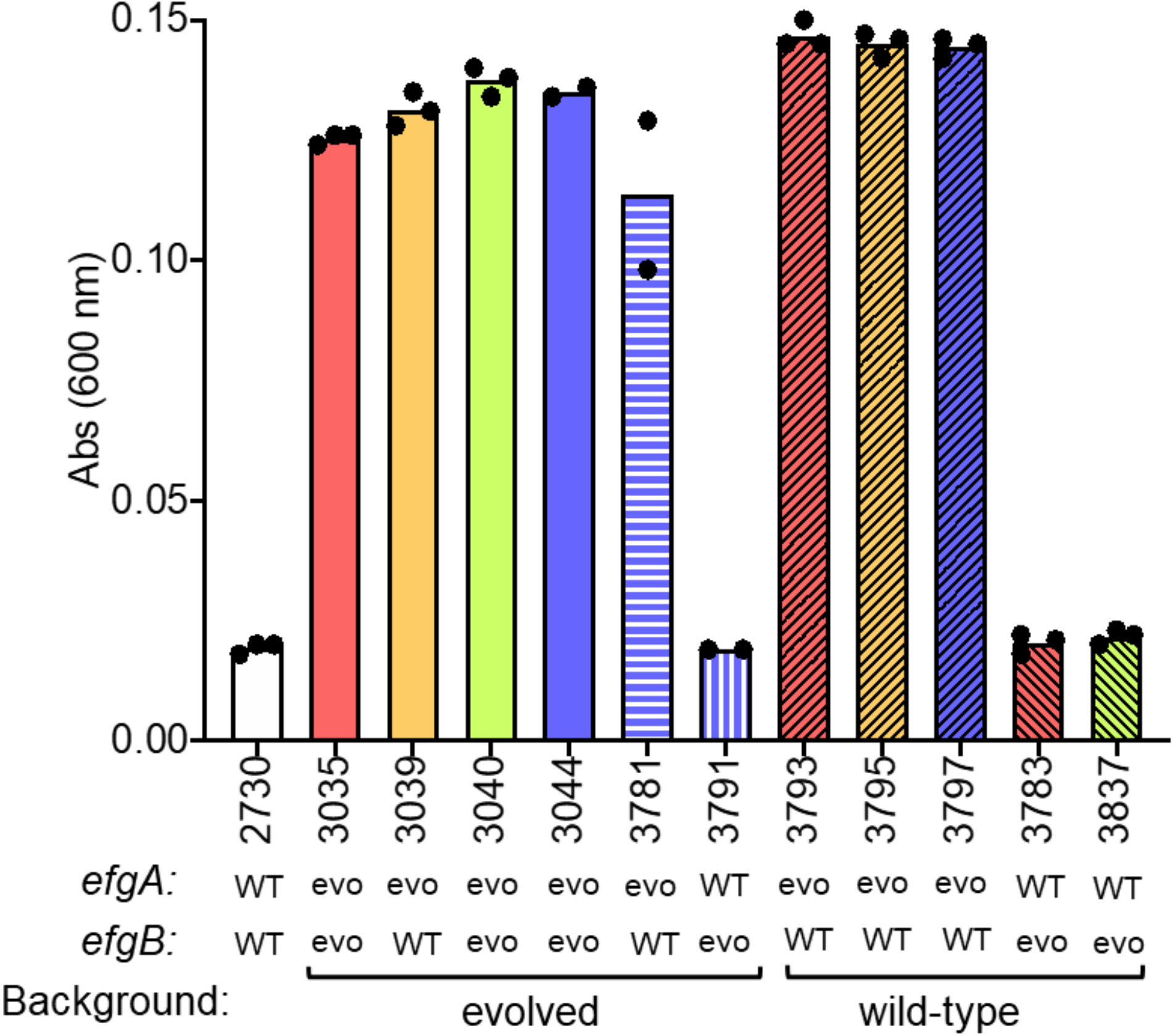
Evolved alleles of *efgA* are necessary and sufficient to confer growth on formaldehyde. Final yields (24 h) of strains grown in liquid MP medium with 5 mM exogenous formaldehyde provided as a sole source of carbon and energy are shown. The wild-type (CM2730) was grown alongside isolates evolved to grow on 20 mM formaldehyde: CM3035 (pink), CM3039 (orange), CM3040 (green), CM3044 (violet). Genetic derivatives of CM3044 where the WT allele replaced the evolved (“evo”) allele are shown with violet and white horizontal (*efgB^WT^*, CM3781) or vertical (*efgA^WT^*, CM3791) bars. Genetic derivatives of wild-type were made by introducing distinct *efgA^evo^* alleles (CM3793, CM3795, and CM3797) or *efgB^evo^* alleles (CM3783, CM3837) present in formaldehyde-evolved isolates; colors are consisten with evolved isolate that alleles were isolated from.

**Figure 4.**
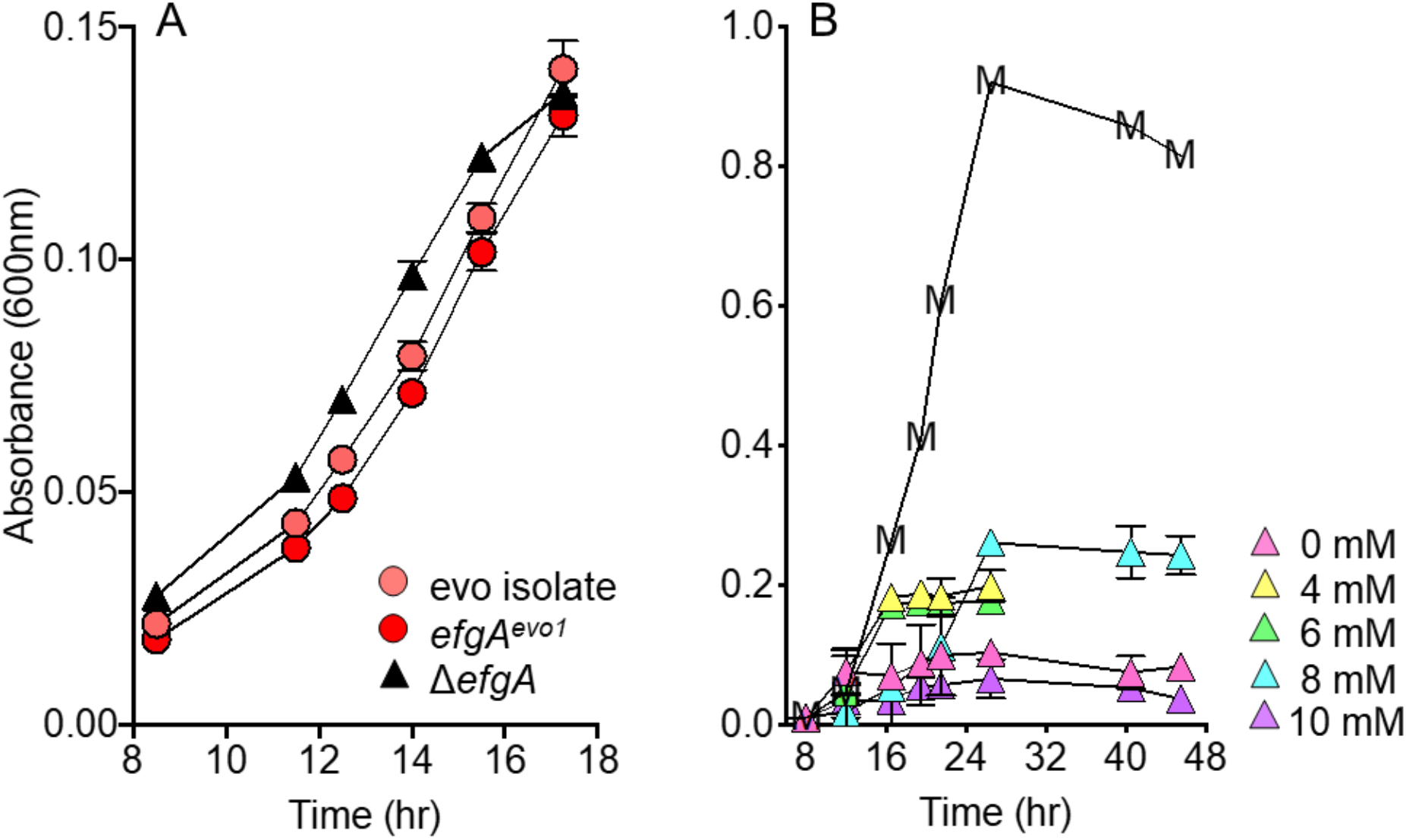
Evolved *efgA* allele or deletion of *efgA* recapitulates formaldehyde growth. (A) Evolved isolate (CM3035, pink), reconstructed *efgA^evo1^* mutant (CM3793, red), and Δ*efgA* mutant (CM3745, black triangles) were grown in liquid Hypho medium with 5 mM exogenous formaldehyde as a sole source of carbon and energy. (B) The Δ*efgA* mutant strain was grown in liquid MP medium with 0, 4, 6, 8, or 10 mM exogenous formaldehyde as a sole source of carbon and energy. For comparison, representative growth with 15 mM methanol (no formaldehyde) is shown (‘M’). Error bars represent the standard error of the mean of three biological replicates.

Further phenotypic analysis of the Δ*efgA* in the wild-type background showed that it enabled growth in medium containing 4-8 mM formaldehyde at initial growth rates comparable to those seen on methanol (Fig 4). The Δ*efgA* strain also exhibited increased formaldehyde resistance in the presence of an alternative growth substrate (Fig S5). Correspondingly, introduction of a second copy of *efgA*^WT^ in the chromosome of wild-type resulted in increased sensitivity to formaldehyde in the presence of an alternative growth substrate (Fig S6). These data show that formaldehyde growth was due to increased formaldehyde resistance and *efgA* plays a key role in the cell’s response to formaldehyde.

### EfgB plays a secondary role in formaldehyde resistance

*efgB* encodes a putative adenylate/guanylate cyclase, a group of regulatory proteins that synthesize cyclic nucleotide second messengers. Adenylate/guanylate cyclases are well-known to exist in essentially all organisms and have been associated with a wide variety of phenotypes, ranging from catabolite repression, induction of virulence, and stress response [96–98]. EfgB displays low identity to CyaA of *E. coli* (6.5%) with its most closely related homologs found in other members of the Rhizobiales with diverse physiologies (Fig S7), rather than methylotrophic organisms from a wide variety of phylogenetic groups.

Genetic analyses with *efgB* alleles indicate that EfgB also plays a role in growth on formaldehyde. Consistent with *efgA^evo^* alleles being necessary and sufficient for formaldehyde growth, the introduction of *efgB^evo^* alone into the wild-type background did not confer growth on 5 mM formaldehyde (Fig 3). When *efgB*^WT^ was introduced into an evolved strain to replace the *efgB^evo1^* allele, no detectable change in growth was observed in medium containing up to 5 mM formaldehyde (Fig 3), however, when Δ*efgB* was introduced into any of the evolved strains growth at higher concentrations of formaldehyde was decreased (Fig S8). These data indicate that the *efgB^evo^* alleles are gain-of-function mutations and led us to hypothesize that they could increase formaldehyde resistance in a genomic background that had already achieved formaldehyde growth.

To more clearly elucidate the role of EfgB, we further examined different *efgB* alleles in the Δ*efgA* and wild-type backgrounds. In the Δ*efgA* background, growth analysis showed that, at higher concentrations of formaldehyde (6-10 mM) the state of the *efgB* allele affected growth in the order *efgB^evo^>efgB^WT^*>Δ*efgB* (Fig 5). In the wild-type background, strains with *efgB^evo^* alleles exhibited modest resistance to formaldehyde, but only when tested at the low concentration of 2 mM formaldehyde in the presence of a primary growth substrate (Fig S9). These data indicate that the *efgB* alleles did not impact growth in the absence of formaldehyde stress but do increase formaldehyde resistance.

**Figure 5.**
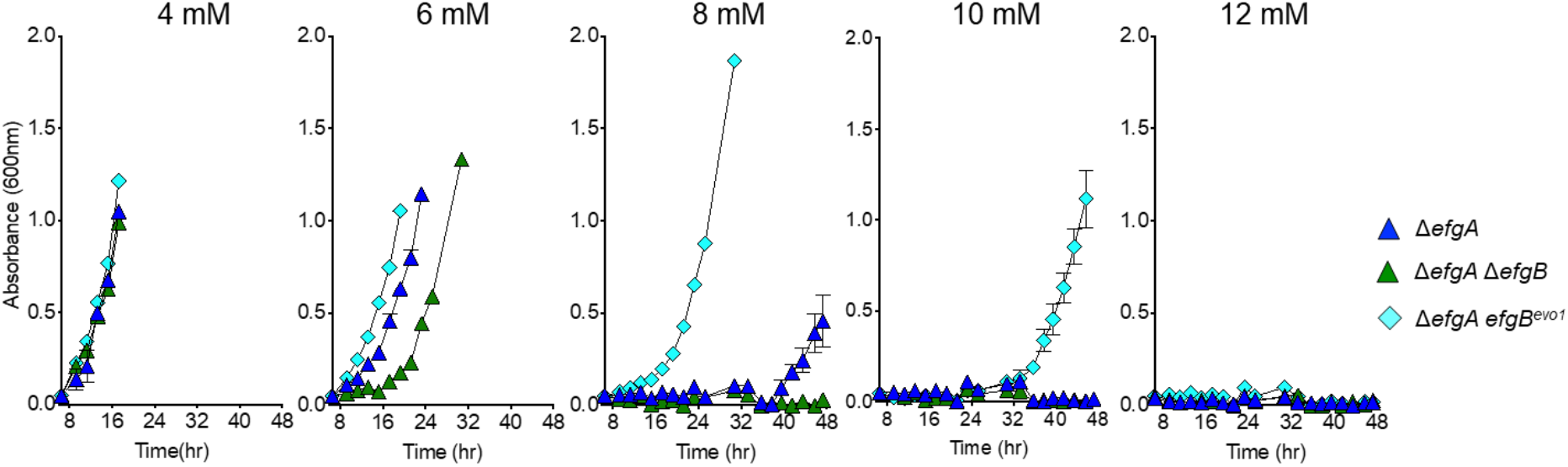
Formaldehyde resistance of *efgA efgB* double mutants indicates *efgB^evo1^* is a gain-of-function allele. The Δ*efgA* (blue triangles), Δ*efgA* Δ*efgB* (green triangles), and Δ*efgA efgB^evo1^* mutants (cyan diamonds) were grown in liquid MP medium with 15 mM succinate and 4, 6, 8, 10, or 12 mM exogenous formaldehyde. Error bars represent the standard error of the mean for three biological replicates.

### EfgA has a formaldehyde-specific role in methylotrophy

Phenotypic analyses of the Δ*efgA* mutant indicate the impact of EfgA is specific to formaldehyde stress. During growth on methanol or succinate alone, the Δ*efgA* mutant phenotype was indistinguishable from wild-type with regard to lag time, growth rate, and final yield (Fig S5). Inhibitory levels of several other aldehydes (glyoxal, acetaldehyde, glutaraldehyde, butyraldehyde, propionaldehyde) did not exhibit a differential effect on Δ*efgA* strains compared to wild-type (Fig S10). These data suggest that EfgA has a formaldehyde-specific role in the cell.

Analysis of *efgB* mutants indicate that EfgB is involved in a broad stress response and not limited to formaldehyde stress. None of the *efgB* alleles tested affected growth on methanol or succinate alone (Fig S9). In contrast to the results with the Δ*efgA* strain, *efgB^evo^* alleles conferred modest resistance to a number of aldehydes in addition to formaldehyde (Fig S11A). This broader stress resistance extended beyond aldehydes, as *efgB^evo^* alleles provided resistance to heat shock as well as a few antibiotics (Fig S11B, C).

Taken together, our data indicate that EfgA has a formaldehyde-specific role in the cell whereas EfgB is involved in resistance to multiple stressors. The specific association of homologs of EfgA, but not EfgB, with methylotrophy further corroborates this formaldehyde-specific role. Our data definitively show that EfgA plays a primary role in formaldehyde resistance unlike EfgB, which plays a secondary role. Correspondingly, we focused our efforts to uncover the biochemical function and role of EfgA.

### The crystal structure of EfgA and molecular dynamics simulations suggest a formaldehyde-binding pocket

To further develop hypotheses regarding the biochemical function of EfgA we determined the structure of an N-terminal His-tagged derivative. EfgA diffracted to 1.65 Å resolution (Protein Data Bank (PDB): 6BWS). The tertiary structure of individual protomers is comprised of an antiparallel β-sheet flanked by four antiparallel α-helices in a mixed topology (Fig S12). The packing of the monomers is consistent with a homotetrameric quaternary structure (Fig 6A). No cofactors or metals copurified with EfgA. With these structural data in hand, we employed a MD+FoldX approach [76] to rationalize the *efgA^evo^* alleles that emerged. This analysis suggested that 11 of the 14 non-synonymous *efgA* mutations either decrease the stability of monomers or of the interactions between them (all but H2Q, M107I, S114N; Fig S13).

**Figure 6.**
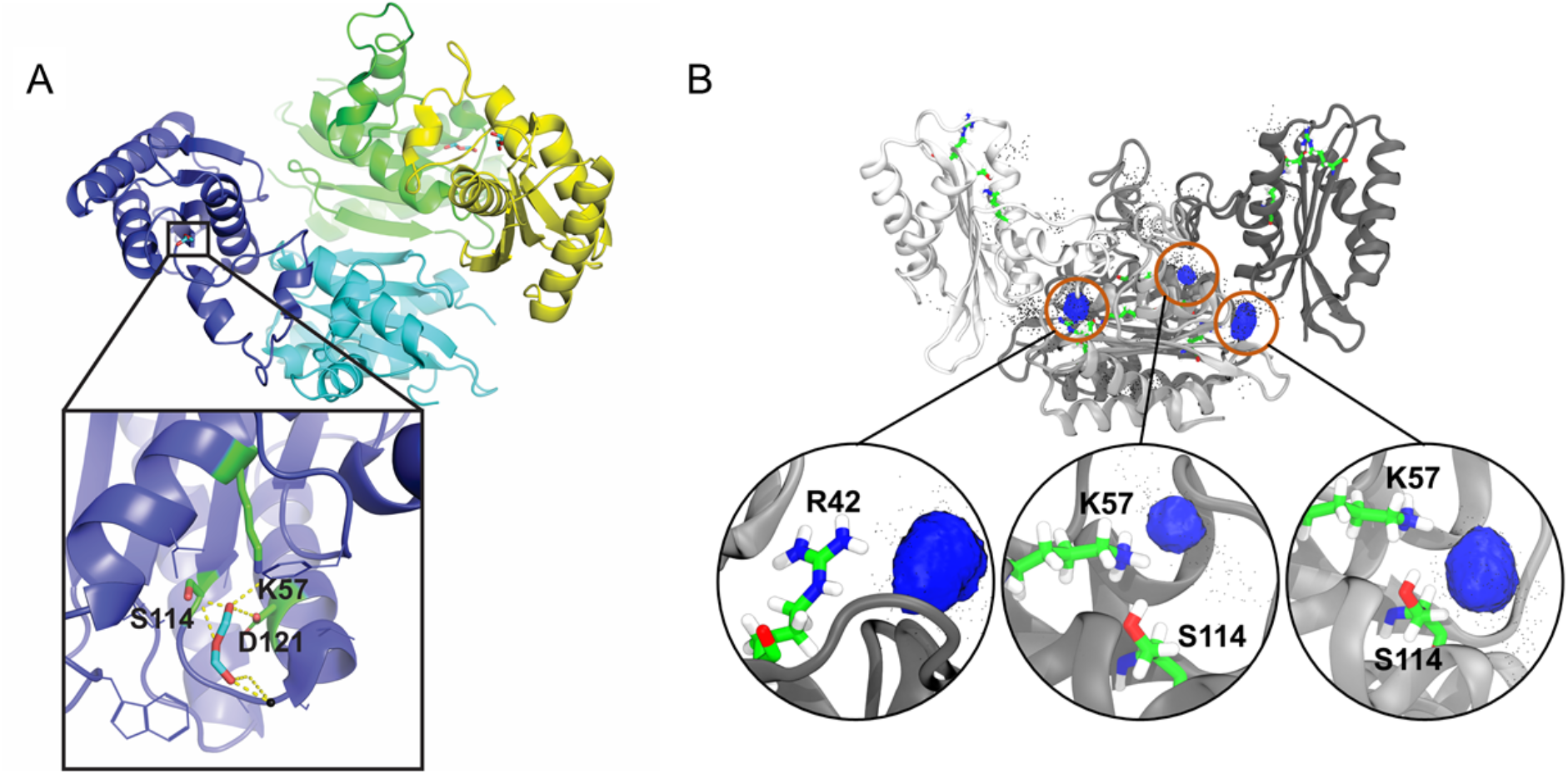
X-ray crystal structure of EfgA and predicted formaldehyde binding sites. (A) EfgA tetramer with each chain highlighted in different colors. Inset: enlarged view of the EfgA binding pocket showing three crucial amino acids, K57, D121, and S114, potentially involved in H-bond interaction with formaldehyde in the crystal structure. (B) EfgA tetramer with each chain highlighted in shades of grey with all 2020 docked poses of formaldehyde and two key amino acids, K57 and S114. Each pose is represented by a single carbon atom (black points) of formaldehyde. Blue clusters demonstrate highly populated regions. Orange circles show an enlarged view of each distinct site. Residues at each site are elementally color-coded, with carbon=green, oxygen=red, nitrogen=blue, and hydrogen=white.

Search for a structural homolog via DALI revealed that the C_α_ positions of the EfgA monomer have a root-mean-square deviation of 1.95 Å (with 127 atoms aligned) for the C_α_ positions of the monomeric chain of HbpS from *S. reticuli* (PDB: 3FPV, Fig 2A). HbpS and EfgA only share 36% amino acid identity and EfgA lacks the twin-arginine translocation signal sequence present in HbpS (Fig S14). We did not expect HbpS and EfgA to have identical functions but were intrigued that EfgA may also function as a stress sensor, specifically hypothesizing that it may sense formaldehyde.

X-ray crystallography of formaldehyde-soaked crystals indicated a specific site where formaldehyde may bind. The structure of the protein with formaldehyde as a potential complexed ligand resolved at 1.83 Å resolution (PDB: 6C0Z). The overall structure was largely unchanged from the apo-protein, but the difference maps comparing the apo and formaldehyde complexes, indicated new electron density in the formaldehyde-soaked crystals. The additional electron density was localized to a specific binding pocket of each monomer. In three EfgA monomers, new densities were modeled as oxydimethanol and in the fourth monomer, the electron density was modeled as formate (Fig 6A). Both oxydimethanol and formate are derivatives of formaldehyde and are structurally homologous. Potential hydrogen bonds were observed between ligands and amino acids that corresponded to S114, D121, and K57 in the native EfgA protein. (Fig 6A inset).

Consistent with the structural data from formaldehyde-soaked crystals, molecular dynamics simulations identified the identical pocket as the likely site of formaldehyde binding. Unbiased docking calculations were performed to dock formaldehyde to snapshots obtained from 100 ns molecular dynamics (MD) simulation initiated from the X-ray crystal structure of apo EfgA tetramer. In total, 2020 docked poses were captured and regions of high-density poses were found to correspond with the binding pockets identified during formaldehyde crystal soaks (Fig 6B). In addition to the primary interaction (K57, D121, and S114), these calculations suggest a feasible interaction at R42. The potential importance of the K57-D121-S114 pocket is emphasized by the fact that one of the loss-of-function *efgA^evo^* alleles (EfgA^S114N^) modifies one of these three proposed binding interactions.

### *In vitro* EfgA:ligand interaction demonstrates direct, specific binding to formaldehyde

To directly test the hypothesis that EfgA senses formaldehyde by direct binding, as suggested by the structural and biophysical modeling evidence above, we used two independent biophysical approaches. Microscale isothermal titration calorimetry (mITC) binding isotherms showed that formaldehyde binding for native EfgA was exothermic (ΔH = –22.65 ± 1.16 kcal/mol), suggesting formaldehyde binding was favorable. Boiled EfgA had a 23-fold decrease in the enthalpy of binding and broadened isotherms, suggesting that the strong interaction of formaldehyde with EfgA required the native 3-dimensional structure, rather than non-specific interactions (Fig 7A, S15A). Microscale thermophoresis (MST) was used to both independently confirm the interaction of EfgA with formaldehyde and permit calculation of its affinity (K_d_=8.01± 3.5 mM) (Fig S16).

**Figure 7.**
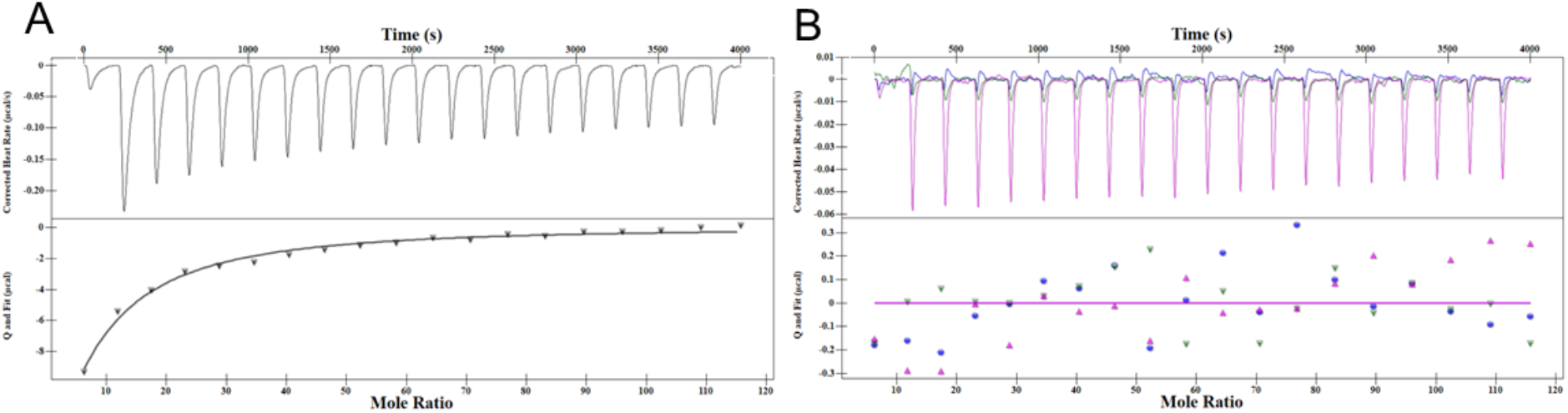
Microscale isothermal calorimetry indicates EfgA binds formaldehyde but not methanol, formate, or acetaldehyde. The binding isotherms represented as heat change (μJ/s) upon injection over time are in the top portion of the split graph, with independent binding modelling on the bottom portion. (A) Binding observed with 50 μM EfgA (black) and 2 μL injections of 25 mM formaldehyde (in PBS). (B) Binding observed with 50 μM EfgA and 2 μL injections of 25 mM methanol (blue), formate (green), acetaldehyde (pink). Data are representative of trends observed in multiple experiments (*n* = 3); additional replicates are shown in Figure S15.

We tested EfgA interactions with two additional categories of alternative ligands that are structurally similar: other C_1_ intermediates (methanol and formate) and a longer aldehyde (acetaldehyde). mITC results indicate no evidence of binding to methanol, formate, or acetaldehyde (Fig 1, Fig 7B, Fig S15D-F). Together, these biochemical data confirm the hypothesis that EfgA specifically binds formaldehyde.

### EfgA homologs from methylotrophs are functionally redundant

Having demonstrated EfgA from *M. extorquens* binds formaldehyde we hypothesized that EfgA homologs found in methylotrophs would have a conserved formaldehyde-sensing function. We attempted to complement a Δ*efgA* strain of *M. extorquens* (an alphaproteobacterium) with *Mfla_1444*, the corresponding gene from *Methylobacillus flagellatus* KT (a betaproteobacterium). Mfla_1444 is an EfgA homolog with 67% identity and 76% similarity and represents a member of the other major clade of EfgA sequences in comparison to the EfgA from *M. extorquens* (Fig 2B). Even the basal, uninduced expression of *Mfla_1444* complemented a Δ*efgA* strain and inhibited formaldehyde growth as well as the native gene (Fig 8). These data suggest that the clade of EfgA homologs found throughout methylotrophs sense formaldehyde and are competent in transmitting that signal to conserved downstream components.

**Figure 8.**
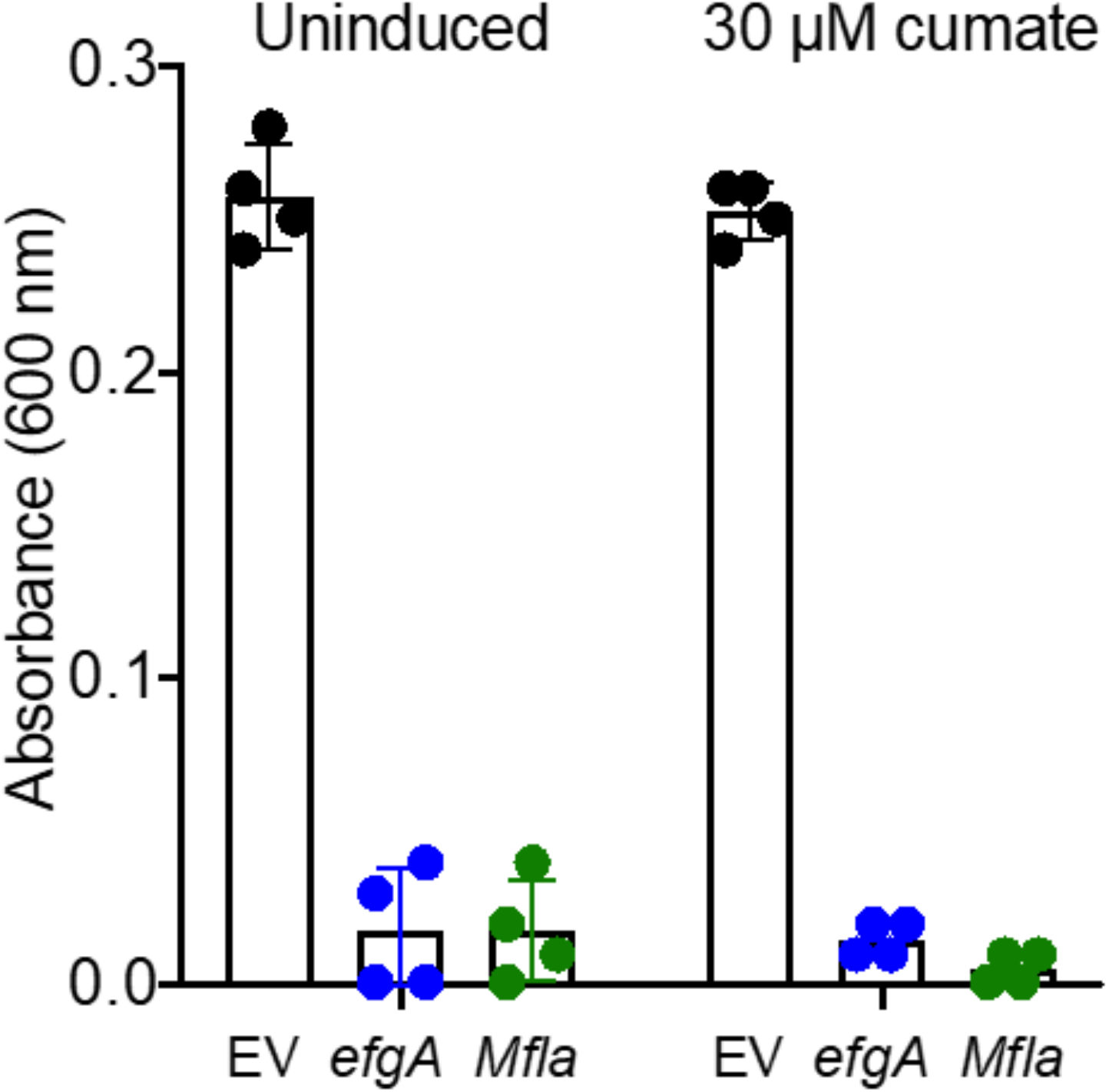
EfgA from *Methylobacillus flagellatus* complements the *M. extorquens* Δ*efgA* mutant. Derivatives of the Δ*efgA* mutant containing pLC290 were grown in liquid MP medium with 8 mM formaldehyde as a sole source of carbon and energy. Strains contained expression plasmids that were either an empty vector control (CM4625, EV, black), expressed EfgA from *M. extorquens* (CM4126, *efgA*, blue), or the homolog from *M. flagellatus* (CM4182, *Mfla_1444*, green). Error bars represent the standard error of the mean for four biological replicates.

### EfgA protects cells from endogenous formaldehyde stress

The direct interaction of EfgA with formaldehyde and its conserved function across methylotrophs led us to question what selective benefit EfgA may provide to cells. The experimental evolution conditions we used had selected for the *removal* of EfgA to allow cells to grow on high concentrations of formaldehyde as a sole carbon source. These conditions are unlikely to be ecologically relevant as they were both extreme and involved exogenous formaldehyde rather than endogenous production as an intermediate. Thus, we hypothesized that the advantage of EfgA regulation may be when methylotrophs experience misbalanced intracellular formaldehyde.

Taking advantage of mutants with partial (Δ*fae*) or complete (Δ*mptG*) lesions in the dH_4_MPT pathway to generate internal formaldehyde stress, we determined that EfgA is beneficial under these conditions. Δ*efgA* Δ*fae* and Δ*efgA* Δ*mptG* strains displayed an exacerbated growth defect compared to the corresponding single mutants with *efgA*^WT^ when 1 mM methanol was added during growth on succinate (Fig 9). These data showed EfgA is beneficial when cells experience endogenous formaldehyde stress. Furthermore, they indicate that the EfgA-mediated response is independent of the dH_4_MPT pathway enzymes and metabolic intermediates.

**Figure 9.**
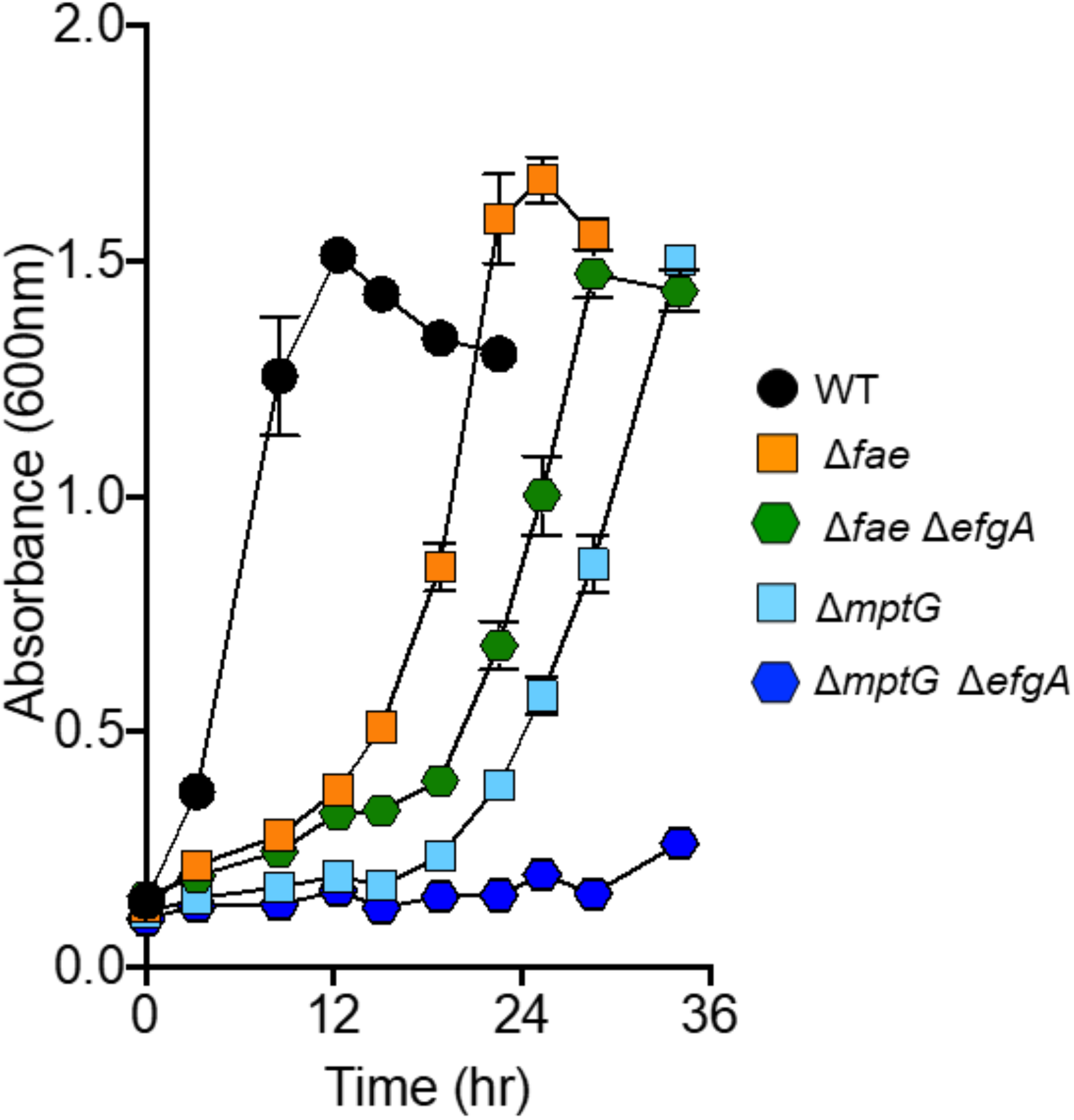
EfgA provides protection from internal formaldehyde in methanol sensitive mutants. Wild-type and mutant strains were grown to early exponential phase in liquid MP medium (succinate) at which point 1 mM methanol was introduced into the medium (t=0 h). Strains represented are wild-type (black circles), Δ*fae* (CM3753, orange squares), Δ*mptG* (CM4765, light blue squares), Δ*efgA* Δ*fae* (CM3421-5, green hexagons), and Δ*efgA* Δ*mptG* mutants (CM3440-13, blue hexagons). Error bars represent the standard error of the mean for three biological replicates.

### EfgA is not linked to phenotypic formaldehyde tolerance

Given that mutations in *efgA* impact genetic resistance to formaldehyde, we questioned whether it was linked to phenotypic formaldehyde tolerance observed between genetically identical cells [33]. *efgA* was not represented amongst the differentially-expressed genes seen in the environmentally-responsive phenotypic variation in formaldehyde tolerance (Table S2). Additionally, neither *efgB* nor any of the other loci identified via mutations during selection for formaldehyde growth had significant changes, other than a single modest change in a secondary locus yet to be investigated (*_2112*, encoding an XRE family transcriptional regulator/shikimate kinase; 1.19-fold change, p-adj=9.28 E-05) in (Table S2). A comparison of the tolerance distributions of wild-type and the Δ*efgA* mutant showed that a distribution of formaldehyde tolerance was maintained within populations of the Δ*efgA* mutant but was shifted toward higher tolerance (Fig 10). Notably, the qualitative shape of the distribution altered with the rate of decline being decreased in the Δ*efgA* mutant (slope: WT=-4.112 log10 cells / mM formaldeyde, Δ*efgA=-1*.482, p-value=0.0004). Further work will be required to determine whether their protein levels or activities may play any role in this response.

**Figure 10.**
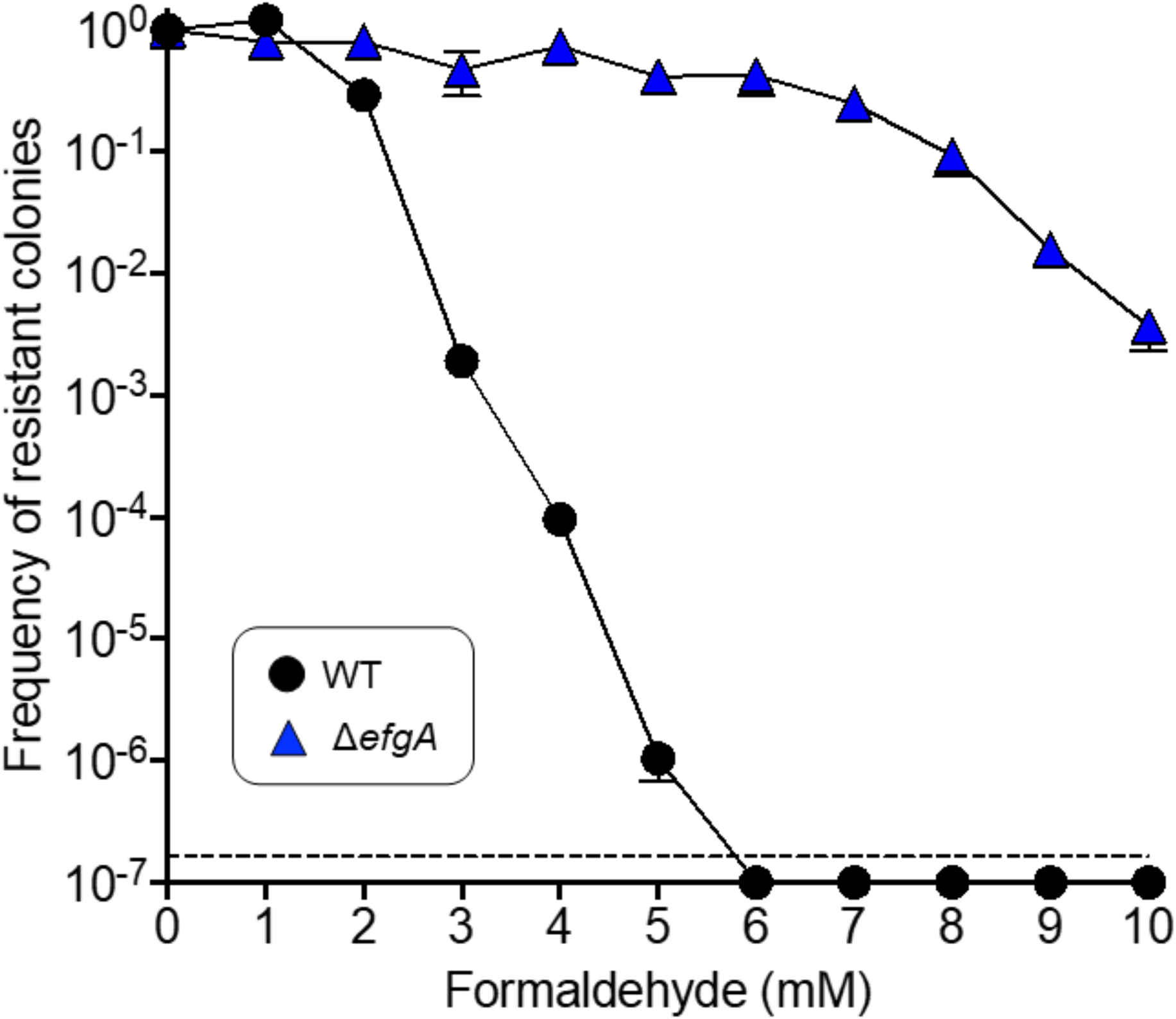
EfgA influences phenotypic formaldehyde tolerance distribution in *M. extorquens*. The distribution of formaldehyde tolerances among individual cells was assessed in WT (black solid circles) and Δ*efgA* (blue triangles) populations. Stationary-phase cultures were plated onto MP-methanol agar medium containing a range of formaldehyde concentrations at 1 mM intervals. The frequency of tolerant cells is expressed as the ratio of the colony-forming units (CFU) enumerated at the given formaldehyde concentration to the CFU enumerated on formaldehyde-free (0 mM) medium. Error bars represent the mean standard deviation of three replicate plates; the horizontal dotted line denotes the limit of detection.

### EfgA can provide protection from formaldehyde in the non-methylotroph *E. coli*

Does EfgA only function in methylotrophs, or might it perhaps provide protection to a heterologous host that is not a methylotroph? Several pieces of evidence led us to hypothesize that this might be possible. First, EfgA senses formaldehyde directly and does not require a functional C_1_ pathway, so these functions should be possible in a different genomic context. Second, we demonstrated that a phylogenetically distinct EfgA homolog can complement a Δ*efgA* mutant, suggesting that any downstream signaling system might be broadly conserved. Third, unlike other DUF336 homologs (e.g., *hbpS, pduO, glcC*), there does not appear to be a conserved genomic context for *efgA* across methylotrophs, suggesting it may have been introduced on its own in the history of those lineages.

Heterologous expression of EfgA in *E. coli* (a gammaproteobacterium) demonstrated EfgA can provide increased formaldehyde resistance in a novel organism. *E. coli* grown in minimal MOPS medium with glucose displays sensitivity to >0.7 mM formaldehyde, but strains expressing EfgA showed a decreased lag time across the formaldehyde concentrations tested (Fig 11). The ability of EfgA to mitigate formaldehyde stress in a new organism, in the absence of native methylotrophic machinery, supported our hypothesis. These data also suggest one of two scenarios exist: EfgA acts independently or EfgA elicits a cellular response via interactions with proteins that are present and sufficiently conserved across non-methylotrophs to enable fortuitous interactions to occur.

**Figure 11.**
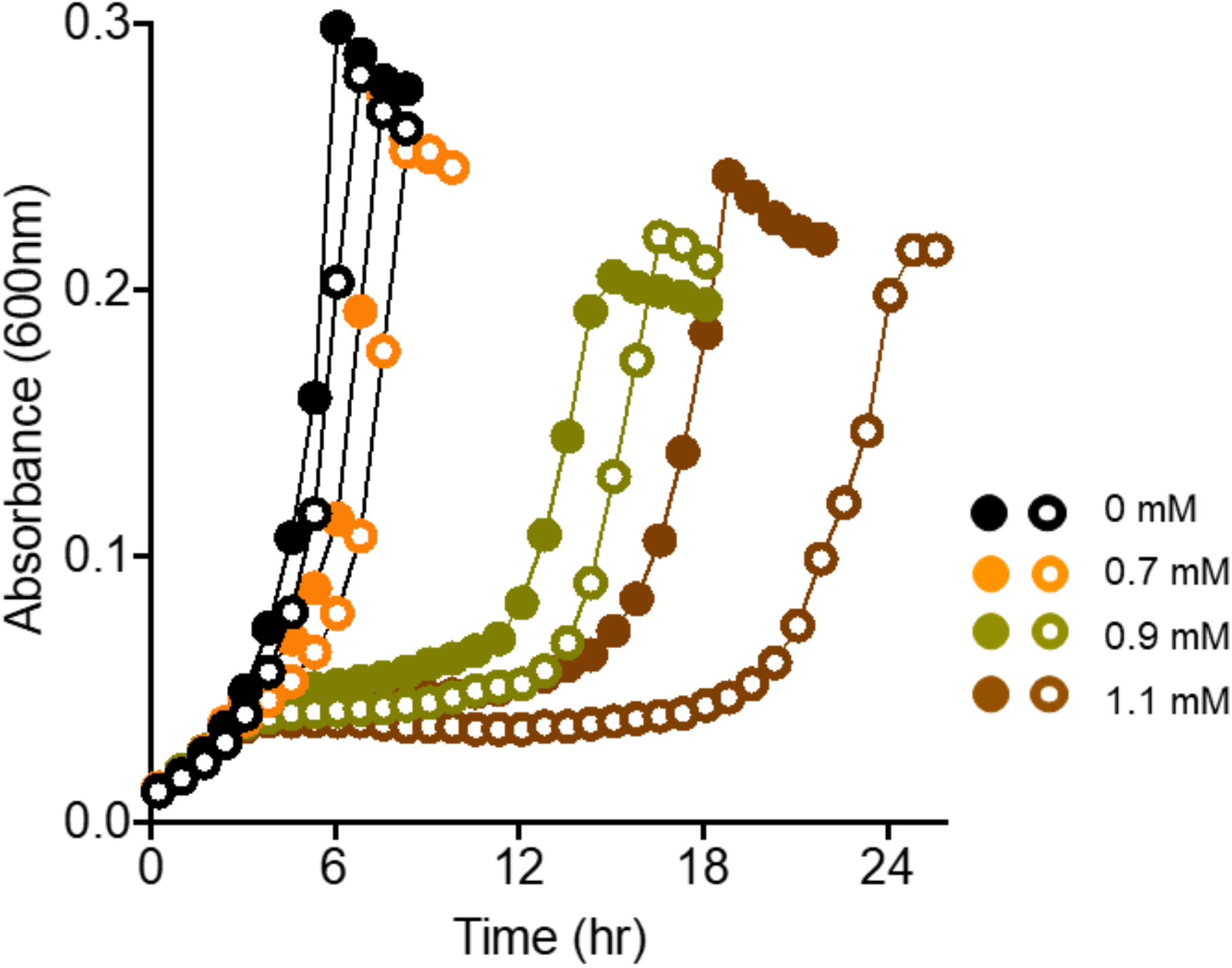
EfgA provides protection to *E. coli* strains during formaldehyde exposure. Growth of wild-type *E. coli* and was quantified in liquid MOPS medium with 2 mM glucose and 0, 0.7, 0.9, 1.0, and 1.1 mM exogenous formaldehyde. Strain WM8637 (filled symbols) harboring a chromosomal insert of *efgA* under the control of P*_rhaS_* or the empty vector control (WM8653, empty symbols) was grown in comparable conditions. *efgA* expression was induced with 0.5 mM rhamnose. Data are representative of trends observed in multiple experiments.

### Alternative loci that permit growth on formaldehyde in the presence of active EfgA include two ribosome-associated proteins

Despite having a functional EfgA, four other loci were targets beneficial, first step mutations that permitted growth on formaldehyde. To glean information about possible downstream effects of EfgA we resequenced the genomes of isolates from the 7 (of 25) populations that rapidly evolved growth on 5 mM formaldehyde but had an *efgA*^WT^ allele. This identified four loci: *potG* (*_4194*, encoding a putative putrescine transporter), *_0925* (encoding a MarR family transcriptional regulator), *prmA* (*_4479*, encoding ribosomal protein L11 methyltransferase), and *def* (*_1636*, encoding peptide deformylase, PDF) (Table 1, S1). An evolved isolate representing each of these four genes was found to have fairly similar fitness (within 10%) to a reconstructed *efgA^evo^* mutant strain during growth on 5 mM formaldehyde (Fig S17). While the role of PotG and the MarR-like regulator remain unclear, the finding that mutations in two different genes encoding ribosome-associated proteins (PrmA and PDF) could protect cells from formaldehyde led us to further explore the unexpected connection between formaldehyde stress and translational events.

### *N*-formylmethionine pathway contributes to formaldehyde resistance

The observation that variants of PDF allowed formaldehyde growth led us to hypothesize that the *N*-formylmethionine (fMet) pathway plays a role in formaldehyde resistance. During translation PDF removes the formyl moiety from the fMet at the N-terminus of a majority of the nascent peptides in bacteria. We were unable to delete *def* in wild-type *M. extorquens*, consistent with the finding that *def* is often individually essential [99–101]. In other bacteria mutants lacking methionyl-tRNA formyltransferase (encoded by *fmt*) do not synthesize fMet and no longer require PDF [100–103]. In a wild-type background, deletions of *fmt* alone or the entire *fmt-def* operon significantly increased sensitivity to formaldehyde (Fig 12). To test whether the lack of fMet precludes the effects of EfgA activity the Δ*fmt* and Δ*fmt-def* alleles were combined with the Δ*efgA* allele. The resulting strains showed that the deletion of *efgA* significantly increased resistance even in the absence of the fMet pathway (Fig 12). These results indicate that EfgA must act, at least in part, independent of the N-terminal protein formylation pathway, but further implicate protein translation/maturation as having a key role in formaldehyde resistance.

**Figure 12.**
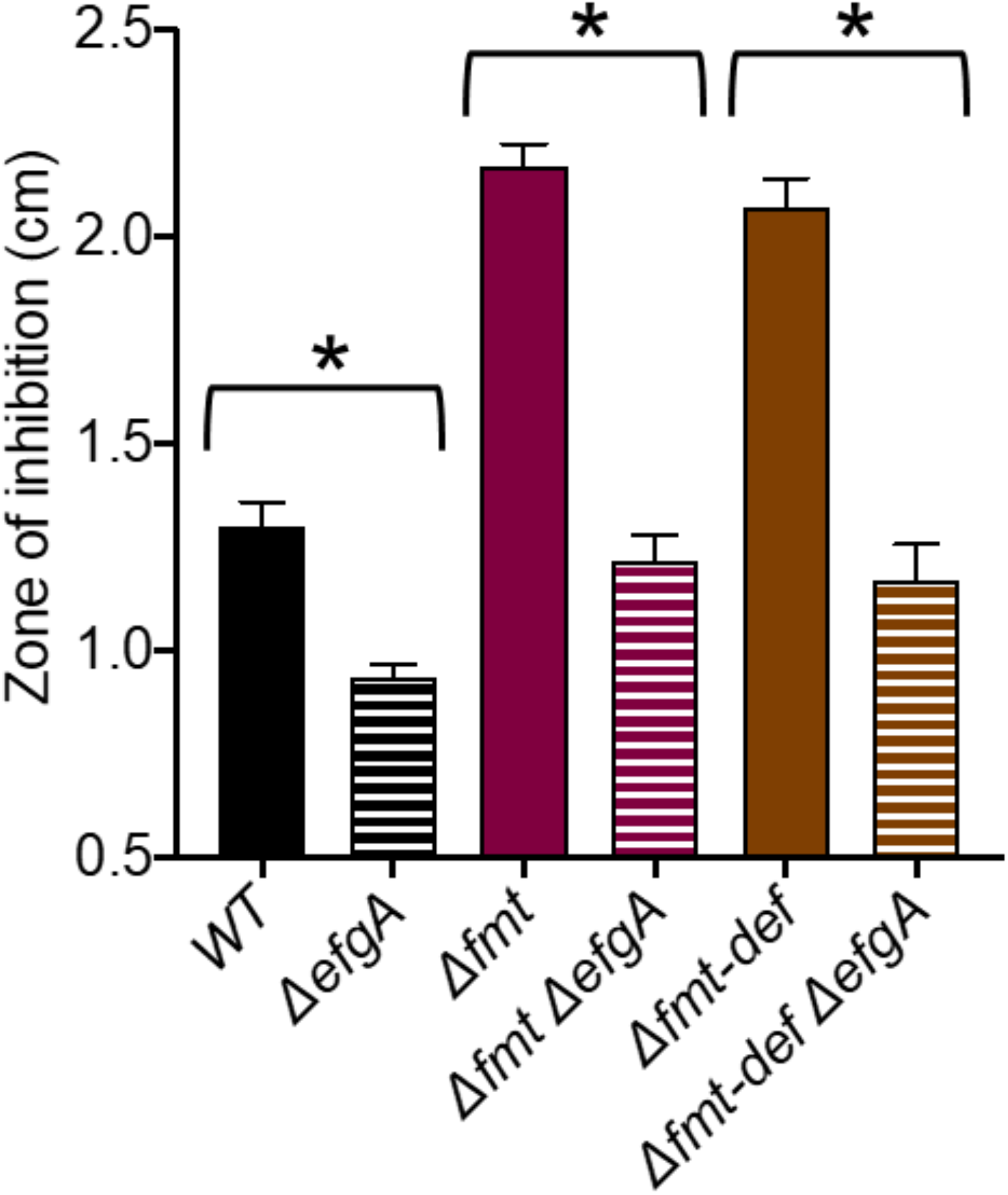
EfgA-mediated formaldehyde resistance is not dependent on PDF. Disc-diffusion assays were performed by placing formaldehyde-impregnated discs upon soft agar overlays of *M. extorquens* on solid MP media (125 mM methanol). The zones of inhibition observed showed that like wild-type, mutants lacking the fMet-mediated protein degradation signal (Δ*fmt*, Δ*fmt-def*) are more formaldehyde-resistant when *efgA* is also deleted. Wild-type and Δ*efgA* mutant were included as experimental controls. Error bars represent the standard error of the mean for three biological replicates. Statistical significance was determined by an unpaired Student’s *t* test (*, p<0.05).

### EfgA halts translation *in vivo* in response to formaldehyde

Multiple findings led us to directly test whether EfgA acts by modulating translation. Translation was assayed *in vivo* by tracking the incorporation of exogenously-provided [^13^CD_3_]-methionine (Met) into cellular proteins of wild-type and an Δ*efgA* mutant by GC/MS. Treating cells with kanamycin, a known translation inhibitor, reduced methionine incorporation by the 360 m timepoint in both genotypes (Fig 13). In contrast, formaldehyde-treated wild-type samples halted translation immediately, showing no detectable increase in methionine incorporation with statistical significance for the full 360 m (Fig 13). For the formaldehyde-treated Δ*efgA* mutant, translation was not halted and increased between all timepoints. The effects of treatments upon translation were largely mirrored in growth; kanamycin inhibition being slow and formaldehyde inhibition in the presence of EfgA being rapid (Fig S18). The addition of formaldehyde to the Δ*efgA* mutant did not induce growth arrest and only led to a modest growth defect. These data indicate that, although formaldehyde does exert some inhibition of translation and growth on its own, the primary effect of excess formaldehyde upon translation and growth is mediated by EfgA.

**Figure 13.**
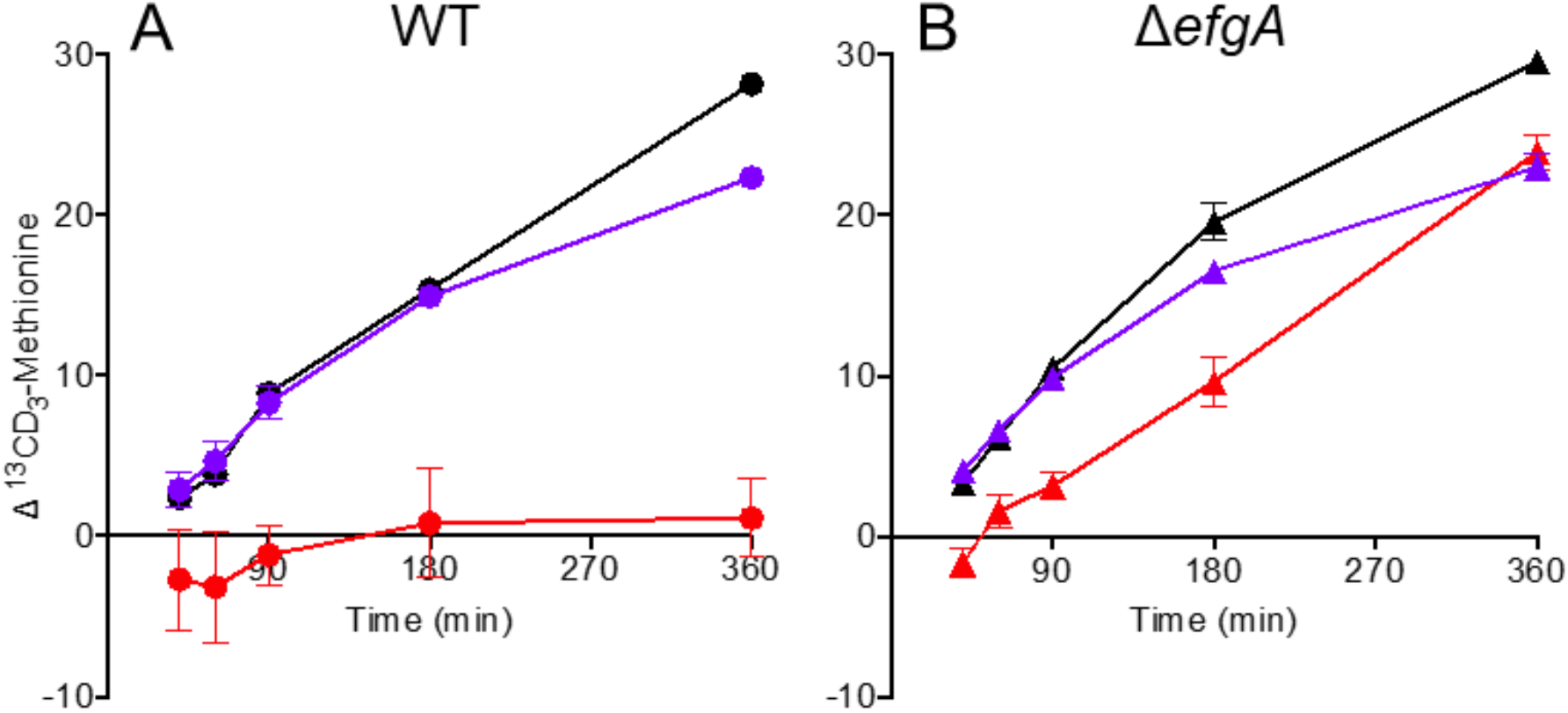
EfgA mediates rapid cessation of translation in response to formaldehyde. Exponential phase cultures of wild-type (circles, A) and Δ*efgA* mutant (triangles, B) strains were treated with kanamycin (purple), exogenous formaldehyde (red) or left untreated (black). After treatment (t >0 min) *in vivo* translation was assayed via [^13^CD_3_]-methionine incorporation. Error bars represent the standard error of the mean for three biological replicates.

## Discussion

Formaldehyde detoxification systems have been identified in all domains of life [104]. Methylotrophs generate formaldehyde at high rate, yet several decades of work with methylotrophs had failed to reveal any proteins that sense and respond to its toxicity. By evolving *M. extorquens* to grow on 20 mM formaldehyde as a sole substrate, a concentration that was found to sterilize a wild-type culture in just two hours [33], we uncovered several genes that could permit growth. Notably, we report here that one of these, now named *efgA* for **<u>e</u>**nhanced **<u>f</u>**ormaldehyde **<u>g</u>**rowth, encodes a formaldehyde sensor that can act through inhibition of protein translation.

We propose that the role of EfgA is to impart protective growth stasis when formaldehyde is transiently elevated to protect from formaldehyde damage. Metabolite-mediated translational arrest has been previously reported for metabolites involved in translation itself, such as amino acids or ppGpp, but not for endogenous stressors [105]. Such a mechanism could limit formaldehyde-induced damage to cellular components, spare protein resources under stress conditions [106], or even reduce further enzymatic production of formaldehyde during high formaldehyde exposure. Nascent peptides emerging from ribosomes may be amongst the most consequential sites of damage and toxicity of formaldehyde crosslinking. Although the mechanism of EfgA inhibition of translation is unclear, the consequent inhibition may be effective because it provides a rapid mechanism for inducing growth arrest. Sustained high external concentrations of formaldehyde that induced permanent growth arrest in wild-type are quite unlikely to be encountered in nature, however, the ability of EfgA to help cells navigate elevated internal formaldehyde stress may be critical in a fluctuating environment. *M. extorquens* are commonly isolated as key members of the leaf microbiome. Although formaldehyde concentrations in plant tissues are relatively low (0.1-10 μmol/g in *Zea mays* and 4.2 μmol/g in *Arabidopsis thaliana*) [107,108], methanol is generated and released in pulses each day [109,110]. This raises the possibility that the beneficial role of EfgA is to mitigate formaldehyde toxicity when formaldehyde production from MDH and usage by the dH_4_MPT pathway become transiently imbalanced. Such imbalances might also arise from metabolic perturbations that can be caused by stressors [111,112], metabolic crosstalk [113], transcriptional bursts [114], or nutrient limitation/shifts [29,115].

Both mITC and MST results confirmed that EfgA binds formaldehyde (Fig 7, S15), but with a relatively high K_d_ of 8 mM (Fig S16). Given that the intracellular concentration of formaldehyde during growth of *M. extorquens* during steady-state growth on methanol has been estimated to be 1 mM [32], a high K_d_ above that concentration would ensure that EfgA occupancy would remain unsaturated during methylotrophic growth, leaving the opportunity for a dynamic response when formaldehyde concentrations rise further. Furthermore, the specificity of EfgA binding of formaldehyde (Fig 7, S15) would render EfgA insensitive to the free C_1_ intermediates upstream and downstream of it, as well as acetaldehyde that would be generated during growth on C_2_ compounds, such as ethanol.

Curiously, the additional electron densities that arose in the formaldehyde-soaked crystals were more consistent with the formaldehyde derivatives, oxydimethanol and formate and not formaldehyde itself. Each of these compounds are derived from formaldehyde and can spontaneously form in aqueous formaldehyde solutions [116]. Aqueous solutions of formaldehyde are composed of formaldehyde and formaldehyde-water derivatives (methylene glycol and oxydimethanol) that exist in equilibrium. Formaldehyde crystal soaks were performed at 10 °C, where the molar ratio of oxydimethanol is nearly at its maximum (0.6 at 8.3°C) [116] and the molar ratio of formaldehyde is ~ 0.3. Therefore, we suspect that the presence of oxydimethanol was an artifact of the experimental conditions but, in fact, its localization is the physiologically relevant site of formaldehyde in the binding pocket. Alternatively, it is possible that EfgA does not sense the unmodified form of formaldehyde directly, but rather an adduct/derivative only formed when formaldehyde is present at sufficiently high concentrations (e.g., oxydimethanol) and which could serve as a proxy for formaldehyde itself. The presence of formate in one of the four monomers may represent an oxidation product formed during crystallization; the lack of interaction seen via mITC when testing formate directly argue against it being the genuine ligand *in vivo*. Taken together, our results validated the protein:ligand interaction as distinct from non-specific binding that might be expected from the ability of formaldehyde to nonspecifically form adducts on and crosslink amino acids residues.

Both X-ray crystallography and *in silico* approaches to dock formaldehyde support the hypothesis that formaldehyde binding demonstrated empirically occurs in a pocket formed via hydrogen bonds with S114, D121 and K57. Fortuitously, a substitution in one of these sites, EfgA^S114N^, was isolated as one of the loss-of-function alleles isolated from one of the evolving populations. Combined with the high degree of conservation at these three residues (Fig S14), these are top candidates for residues involved in ligand binding and will be investigated further in future work.

The discovery that EfgA is a formaldehyde sensor indicates a previously unknown role for DUF336 proteins in sensing small metabolites. Proteins with DUF336 domains are present in single- or multi-domain proteins and their genetic association with gene clusters that encode well-characterized pathways such as glycolate utilization in *E. coli* (GlcG), 1,2-propanediol utilization in *Salmonella enterica* (PduO), and chloroaniline degradation in *Diaphorobacter* sp. PCA039 (OrfU2) has been noted [82–86]. The two best characterized examples are HbpS of *S. reticuli* and PduO of *S. enterica*. HbpS is located extracellularly where it senses and degrades heme and activates a two-component system involved in oxidative stress [88,92,117,118]. PduO is localized to the *pdu* microcompartment its DUF336 domain is fused to an ATP:cob(I)alamin adenosyltransferase domain; its DUF336 domain is not required for the activity of the ATP:Cob(I)alamin adenosyltransferase domain *in vitro* but is required for optimal 1,2-propanediol utilization for unknown reasons [86]. Both HbpS and PduO have been shown to bind heme and cobalamin but their cellular locations are distinct, consistent with their disparate functions. In this light, EfgA represents the third biochemically-characterized bacterial DUF336 protein and is distinct from HbpS in that it is cytoplasmic, senses formaldehyde, and modulates translation. EfgA and HbpS overlap in terms of being sensors; EfgA and PduO may exhibit functional overlap due to the involvement of an aldehyde in propanediol utilization. Thus, our work, which assigns a novel, aldehyde sensing function for a DUF336 protein, helps to define the broader role of DUF336 domains.

The involvement of several genes with known involvement in translation suggests that controlling protein damage may be particularly critical for cells confronted with formaldehyde stress. Two secondary mutations that we have yet to examine also suggest the key role for protein quality control in formaldehyde resistance. For *hrcA^evo^*, the apparent loss-of-function mutation would eliminate HrcA, a heat-inducible transcriptional repressor that negatively regulates heat shock genes [119,120]. The intergenic *_3827/_3828* mutation is upstream of *_3828*, annotated as encoding HdeA, a periplasmic chaperone protein that prevents aggregation of periplasmic proteins [121,122]. These mutations suggest that preventing/repairing protein damage may be important for formaldehyde resistance and imply that unchecked formaldehyde stress leads to protein damage in *M. extorquens*. Though formaldehyde damage is most commonly associated with DNA damage, there is precedent for formaldehyde-induced protein damage [13,123] which was also suggested by our previous work [33].

While the catalytic activity of PDF has long been known, it has only recently been identified as a key player in protein quality control [7,124,125]. Typically, as the fMet of an elongating peptide chain emerges from the ribosome exit tunnel, PDF quickly removes its formyl group, making it a suitable substrate for the downstream processing enzyme. In instances where the elongating peptide is misfolded, fMet is less accessible to PDF and serves as a signal for protein degradation [124]. We isolated *def^evo^* alleles that, through an unknown mechanism, increase formaldehyde resistance, whereas Δ*fmt* and Δ*fmt-def* strains with no fMet cycling decreased formaldehyde resistance. To our knowledge, this is the first indication that fMet modification plays a role in formaldehyde resistance.

The surprising ability for EfgA homologs to influence formaldehyde resistance when introduced between distantly related organisms raises an intriguing possibility that EfgA may directly interact with one or more ribosomal or ribosome-associated components to halt translation. There is precedent for a DUF336 protein to physically interact with ribosome-associated proteins. The homolog in *Saccharomyces cerevisiae*, Ybr137wp, has been shown to be involved in post-translational control, whereby it binds Sgt2 and Get proteins and is involved in the co-translational targeting of tail-anchored proteins into membranes [126]. If EfgA interacts with ribosomal or ribosome-associated proteins, it should be noted that molecular composition of ribosomes and the sequences of the molecular components are amongst most well-conserved aspects of bacteria. The finding that EfgA from *M. extorquens* can provide protection from formaldehyde in *E. coli* may rely upon the fact that its interaction partners were already encoded and expressed there. Having downstream partners already present in genomes could also explain how *efgA* could be acquired by horizontal gene transfer and integrated into the genome as isolated genes without conserved synteny. This would be quite distinct from the rampant exchange of methylotrophy pathways by horizontal gene transfer which appear to have been cointroduced as large, genomically-clustered modules [127]. The ability for EfgA to provide an immediate benefit in dealing with formaldehyde stress may also have more immediate biotechnological benefits. Given that formaldehyde toxicity was also a key challenge in the engineering and evolution of an *E. coli* strain that can grow on methanol as a sole carbon source [128], this raises the possibility that introducing EfgA would increase the cells’ ability to grow while producing formaldehyde as a high flux intermediate. We perhaps should not be surprised that metabolic pathways that generate toxic intermediates need molecular systems to sense their accumulation and mount responses that either eliminate the toxin, increase the ability to repair such damage, or help the cell avoid making the molecules that the toxin damages in the first place.

## Acknowledgements

We thank members of the Marx lab, Lauren D. Palmer, and Andrew J. Borchert for critical reading of this manuscript. We thank Paige K. Swanson for construction of pPS04 and Caleb J. Renshaw for construction of pCJR1.

## References

1. Imlay JA. Cellular defenses against superoxide and hydrogen peroxide. Annu Rev Biochem. 2008;77: 755–776. doi:10.1146/annurev.biochem.77.061606.161055

2. Chen NH, Djoko KY, Veyrier FJ, McEwan AG. Formaldehyde Stress Responses in Bacterial Pathogens. Front Microbiol. 2016;7: 257. doi:10.3389/fmicb.2016.00257

3. Borchert AJ, Ernst DC, Downs DM. Reactive Enamines and Imines *In Vivo*: Lessons from the RidA Paradigm. Trends Biochem Sci. 2019;44: 849–860. doi:10.1016/j.tibs.2019.04.011

4. Downs DM, Ernst DC. From microbiology to cancer biology: the Rid protein family prevents cellular damage caused by endogenously generated reactive nitrogen species. Mol Microbiol. 2015;96: 211–219. doi:10.1111/mmi.12945

5. Niehaus TD, Hillmann KB. Enzyme promiscuity, metabolite damage, and metabolite damage control systems of the tricarboxylic acid cycle. FEBS J. 2020;287: 1343–1358. doi:10.1111/febs.15284

6. Zhang H, Xiong Y, Chen J. DNA-protein cross-link repair: what do we know now? Cell Biosci. 2020;10: 3. doi:10.1186/s13578-019-0366-z

7. Varshavsky A. N-degron and C-degron pathways of protein degradation. Proc Natl Acad Sci U S A. 2019;116: 358–366. doi:10.1073/pnas.1816596116

8. Grove A. Regulation of Metabolic Pathways by MarR Family Transcription Factors. Comput Struct Biotechnol J. 2017;15: 366–371. doi:10.1016/j.csbj.2017.06.001

9. Kamps JJAG, Hopkinson RJ, Schofield CJ, Claridge TDW. How formaldehyde reacts with amino acids. Communications Chemistry. 2019;2: 126. doi:10.1038/s42004-019-0224-2

10. Kawanishi M, Matsuda T, Yagi T. Genotoxicity of formaldehyde: molecular basis of DNA damage and mutation. Front Environ Sci Eng China. 2014;2: 36. doi:10.3389/fenvs.2014.00036

11. Metz B, Kersten GFA, Hoogerhout P, Brugghe HF, Timmermans HAM, de Jong A, et al. Identification of formaldehyde-induced modifications in proteins: reactions with model peptides. J Biol Chem. 2004;279: 6235–6243. doi:10.1074/jbc.M310752200

12. Toews J, Rogalski JC, Clark TJ, Kast J. Mass spectrometric identification of formaldehyde-induced peptide modifications under *in vivo* protein cross-linking conditions. Anal Chim Acta. 2008;618: 168–183. doi:10.1016/j.aca.2008.04.049

13. Ortega-Atienza S, Rubis B, McCarthy C, Zhitkovich A. Formaldehyde Is a Potent Proteotoxic Stressor Causing Rapid Heat Shock Transcription Factor 1 Activation and Lys48-Linked Polyubiquitination of Proteins. Am J Pathol. 2016;186: 2857–2868. doi:10.1016/j.ajpath.2016.06.022

14. Achkor H, Díaz M, Fernández MR, Biosca JA, Parés X, Martínez MC. Enhanced formaldehyde detoxification by overexpression of glutathione-dependent formaldehyde dehydrogenase from *Arabidopsis*. Plant Physiol. 2003;132: 2248–2255. doi:10.1104/pp.103.022277

15. Burgos-Barragan G, Wit N, Meiser J, Dingler FA, Pietzke M, Mulderrig L, et al. Mammals divert endogenous genotoxic formaldehyde into one-carbon metabolism. Nature. 2017;548: 549–554. doi:10.1038/nature23481

16. Pontel LB, Rosado IV, Burgos-Barragan G, Garaycoechea JI, Yu R, Arends MJ, et al. Endogenous Formaldehyde Is a Hematopoietic Stem Cell Genotoxin and Metabolic Carcinogen. Mol Cell. 2015;60: 177–188. doi:10.1016/j.molcel.2015.08.020

17. Marx CJ, Miller JA, Chistoserdova L, Lidstrom ME. Multiple formaldehyde oxidation/detoxification pathways in *Burkholderia fungorum* LB400. J Bacteriol. 2004;186: 2173–2178. doi:10.1128/jb.186.7.2173-2178.2004

18. Denby KJ, Iwig J, Bisson C, Westwood J, Rolfe MD, Sedelnikova SE, et al. The mechanism of a formaldehyde-sensing transcriptional regulator. Sci Rep. 2016;6: 38879. doi:10.1038/srep38879

19. Schoeler M, Caesar R. Dietary lipids, gut microbiota and lipid metabolism. Rev Endocr Metab Disord. 2019;20: 461–472. doi:10.1007/s11154-019-09512-0

20. Onyszkiewicz M, Jaworska K, Ufnal M. Short chain fatty acids and methylamines produced by gut microbiota as mediators and markers in the circulatory system. Exp Biol Med. 2020;245: 166–175. doi:10.1177/1535370219900898

21. McTaggart TL, Beck DAC, Setboonsarng U, Shapiro N, Woyke T, Lidstrom ME, et al. Genomics of Methylotrophy in Gram-Positive Methylamine-Utilizing Bacteria. Microorganisms. 2015;3: 94–112. doi:10.3390/microorganisms3010094

22. Chistoserdova L. Applications of methylotrophs: can single carbon be harnessed for biotechnology? Curr Opin Biotechnol. 2018;50: 189–194. doi:10.1016/j.copbio.2018.01.012

23. Green PN, Ardley JK. Review of the genus *Methylobacterium* and closely related organisms: a proposal that some *Methylobacterium* species be reclassified into a new genus, *Methylorubrum* gen. nov. Int J Syst Evol Microbiol. 2018;68: 2727–2748. doi:10.1099/ijsem.0.002856

24. Chen BJ, Jean GC. Active transport of formaldehyde in methanol-utilizing bacteria. The Chemical Engineering Journal. 1982;25: 9–20. doi:10.1016/0300-9467(82)85017-X

25. Marx CJ, Chistoserdova L, Lidstrom ME. Formaldehyde-detoxifying role of the tetrahydromethanopterin-linked pathway in *Methylobacterium extorquens AM1*. J Bacteriol. 2003;185: 7160–7168. doi:10.1128/jb.185.23.7160-7168.2003

26. Chistoserdova L, Vorholt JA, Thauer RK, Lidstrom ME. C1 transfer enzymes and coenzymes linking methylotrophic bacteria and methanogenic Archaea. Science. 1998;281: 99–102. doi:10.1126/science.281.5373.99

27. Vorholt JA, Marx CJ, Lidstrom ME, Thauer RK. Novel Formaldehyde-Activating Enzyme in Methylobacterium *extorquens* AM1 Required for Growth on Methanol. J Bacteriol. 2000;182: 6645–6650. doi:10.1128/JB.182.23.6645-6650.2000

28. Crowther GJ, Kosály G, Lidstrom ME. Formate as the main branch point for methylotrophic metabolism in *Methylobacterium extorquens* AM1. J Bacteriol. 2008;190: 5057–5062. doi:10.1128/JB.00228-08

29. Skovran E, Crowther GJ, Guo X, Yang S, Lidstrom ME. A systems biology approach uncovers cellular strategies used by *Methylobacterium extorquens* AM1 during the switch from multi-to single-carbon growth. PLoS One. 2010;5: e14091. doi:10.1371/journal.pone.0014091

30. Laukel M, Rossignol M, Borderies G, Völker U, Vorholt JA. Comparison of the proteome of *Methylobacterium extorquens* AM1 grown under methylotrophic and nonmethylotrophic conditions. Proteomics. 2004;4: 1247–1264. doi:10.1002/pmic.200300713

31. Bosch G, Skovran E, Xia Q, Wang T, Taub F, Miller JA, et al. Comprehensive proteomics of *Methylobacterium extorquens* AM1 metabolism under single carbon and nonmethylotrophic conditions. Proteomics. 2008;8: 3494–3505. doi:10.1002/pmic.200800152

32. Martinez-Gomez NC, Good NM, Lidstrom ME. Methenyl-Dephosphotetrahydromethanopterin Is a Regulatory Signal for Acclimation to Changes in Substrate Availability in *Methylobacterium extorquens* AM1. J Bacteriol. 2015;197: 2020–2026. doi:10.1128/JB.02595-14

33. Lee JA, Riazi S, Nemati S, Bazurto JV, Vasdekis AE, Ridenhour BJ, et al. Microbial phenotypic heterogeneity in response to a metabolic toxin: Continuous, dynamically shifting distribution of formaldehyde tolerance in *Methylobacterium extorquens* populations. PLoS Genet. 2019;15: e1008458. doi:10.1371/journal.pgen.1008458

34. Marx CJ. Evolution as an experimental tool in microbiology:’Bacterium, improve thyself!’. Environ Microbiol Rep. 2011. Available: https://dash.harvard.edu/handle/1/34262173

35. Marx CJ. Experimental Evolution of *Methylobacterium*: 15 Years of Planned Experiments and Surprise Findings. Curr Issues Mol Biol. 2019;33: 249–266. doi:10.21775/cimb.033.249

36. Chistoserdova L, Chen S-W, Lapidus A, Lidstrom ME. Methylotrophy in *Methylobacterium extorquens* AM1 from a genomic point of view. J Bacteriol. 2003;185: 2980–2987. doi:10.1128/jb.185.10.2980-2987.2003

37. Marx CJ, Bringel F, Chistoserdova L, Moulin L, Farhan Ul Haque M, Fleischman DE, et al. Complete genome sequences of six strains of the genus *Methylobacterium*. J Bacteriol. 2012;194: 4746–4748. doi:10.1128/JB.01009-12

38. Knief C, Frances L, Vorholt JA. Competitiveness of diverse *Methylobacterium* strains in the phyllosphere of *Arabidopsis thaliana* and identification of representative models, including *M. extorquens* PA1. Microb Ecol. 2010;60: 440–452. doi:10.1007/s00248-010-9725-3

39. Delaney NF, Kaczmarek ME, Ward LM, Swanson PK, Lee M-C, Marx CJ. Development of an optimized medium, strain and high-throughput culturing methods for *Methylobacterium extorquens*. PLoS One. 2013;8: e62957. doi:10.1371/journal.pone.0062957

40. Haldimann A, Wanner BL. Conditional-replication, integration, excision, and retrieval plasmid-host systems for gene structure-function studies of bacteria. J Bacteriol. 2001;183: 6384–6393. doi:10.1128/JB.183.21.6384-6393.2001

41. Lee M-C, Chou H-H, Marx CJ. Asymmetric, bimodal trade-offs during adaptation of *Methylobacterium* to distinct growth substrates. Evolution. 2009;63: 2816–2830. doi:10.1111/j.1558-5646.2009.00757.x

42. Neidhardt FC, Bloch PL, Smith DF. Culture medium for enterobacteria. J Bacteriol. 1974;119: 736–747. doi:10.1128/JB.119.3.736-747.1974

43. Gumerov VM, Ortega DR, Adebali O, Ulrich LE, Zhulin IB. MiST 3.0: an updated microbial signal transduction database with an emphasis on chemosensory systems. Nucleic Acids Res. 2020;48: D459–D464. doi:10.1093/nar/gkz988

44. Altschul SF, Madden TL, Schäffer AA, Zhang J, Zhang Z, Miller W, et al. Gapped BLAST and PSI-BLAST: a new generation of protein database search programs. Nucleic Acids Res. 1997;25: 3389–3402. doi:10.1093/nar/25.17.3389

45. Pruitt KD, Tatusova T, Maglott DR. NCBI Reference Sequence (RefSeq): a curated non-redundant sequence database of genomes, transcripts and proteins. Nucleic Acids Res. 2005;33: D501–4. doi:10.1093/nar/gki025

46. Li W, Godzik A. CD-HIT: a fast program for clustering and comparing large sets of protein or nucleotide sequences. Bioinformatics. 2006;22: 1658–1659. doi:10.1093/bioinformatics/btl158

47. Edgar RC. MUSCLE: multiple sequence alignment with high accuracy and high throughput. Nucleic Acids Res. 2004;32: 1792–1797. doi:10.1093/nar/gkh340

48. Price MN, Dehal PS, Arkin AP. FastTree: computing large minimum evolution trees with profiles instead of a distance matrix. Mol Biol Evol. 2009;26: 1641–1650. doi:10.1093/molbev/msp077

49. Le SQ, Gascuel O. An improved general amino acid replacement matrix. Mol Biol Evol. 2008;25: 1307–1320. doi:10.1093/molbev/msn067

50. Whelan S, Goldman N. A general empirical model of protein evolution derived from multiple protein families using a maximum-likelihood approach. Mol Biol Evol. 2001;18: 691–699. doi:10.1093/oxfordjournals.molbev.a003851

51. Jones DT, Taylor WR, Thornton JM. The rapid generation of mutation data matrices from protein sequences. Comput Appl Biosci. 1992;8: 275–282. doi:10.1093/bioinformatics/8.3.275

52. Letunic I, Bork P. Interactive tree of life (iTOL) v3: an online tool for the display and annotation of phylogenetic and other trees. Nucleic Acids Res. 2016;44: W242–5. doi:10.1093/nar/gkw290

53. Nei M, Kumar S. Molecular Evolution and Phylogenetics. Oxford University Press; 2000. Available: https://play.google.com/store/books/details?id=hcPSag2pn9IC

54. Felsenstein J. Confidence Limits On Phylogenies: An Approach Using the Bootstrap. Evolution. 1985;39: 783–791. doi:10.1111/j.1558-5646.1985.tb00420.x

55. Kumar S, Stecher G, Li M, Knyaz C, Tamura K. MEGA X: Molecular Evolutionary Genetics Analysis across Computing Platforms. Mol Biol Evol. 2018;35: 1547–1549. doi:10.1093/molbev/msy096

56. Marx CJ. Development of a broad-host-range *sacB-based* vector for unmarked allelic exchange. BMC Res Notes. 2008;1: 1. doi:10.1186/1756-0500-1-1

57. Marx CJ, Lidstrom ME. Development of improved versatile broad-host-range vectors for use in methylotrophs and other Gram-negative bacteriaThe GenBank accession numbers for the sequences reported in this paper are AF327711, AF327712, AF327713, AF327714, AF327715, AF327716, AF327717, AF327718, AF327719 and AF327720. Microbiology. 2001;147: 2065–2075. Available: https://www.microbiologyresearch.org/content/journal/micro/10.1099/00221287-147-8-2065

58. Figurski DH, Helinski DR. Replication of an origin-containing derivative of plasmid RK2 dependent on a plasmid function provided in trans. Proc Natl Acad Sci U S A. 1979;76: 1648–1652. doi:10.1073/pnas.76.4.1648

59. Nash T. The colorimetric estimation of formaldehyde by means of the Hantzsch reaction. Biochem J. 1953;55: 416–421. doi:10.1042/bj0550416

60. Otwinowski Z, Minor W. [20] Processing of X-ray diffraction data collected in oscillation mode. Methods in Enzymology. Academic Press; 1997. pp. 307–326. doi:10.1016/S0076-6879(97)76066-X

61. McCoy AJ, Grosse-Kunstleve RW, Adams PD, Winn MD, Storoni LC, Read RJ. Phaser crystallographic software. J Appl Crystallogr. 2007;40: 658–674. doi:10.1107/S0021889807021206

62. Emsley P, Cowtan K. Coot: model-building tools for molecular graphics. Acta Crystallogr D Biol Crystallogr. 2004;60: 2126–2132. doi:10.1107/S0907444904019158

63. Adams PD, Afonine PV, Bunkóczi G, Chen VB, Davis IW, Echols N, et al. PHENIX: a comprehensive Python-based system for macromolecular structure solution. Acta Crystallogr D Biol Crystallogr. 2010;66: 213–221. doi:10.1107/S0907444909052925

64. Davis IW, Leaver-Fay A, Chen VB, Block JN, Kapral GJ, Wang X, et al. MolProbity: all-atom contacts and structure validation for proteins and nucleic acids. Nucleic Acids Res. 2007;35: W375–83. doi:10.1093/nar/gkm216

65. Van Der Spoel D, Lindahl E, Hess B, Groenhof G, Mark AE, Berendsen HJC. GROMACS: fast, flexible, and free. J Comput Chem. 2005;26: 1701–1718. doi:10.1002/jcc.20291

66. Trott O, Olson AJ. AutoDock Vina: improving the speed and accuracy of docking with a new scoring function, efficient optimization, and multithreading. J Comput Chem. 2010;31: 455–461. doi:10.1002/jcc.21334

67. Hornak V, Abel R, Okur A, Strockbine B, Roitberg A, Simmerling C. Comparison of multiple Amber force fields and development of improved protein backbone parameters. Proteins. 2006;65: 712–725. doi:10.1002/prot.21123

68. Berendsen HJC, Postma JPM, van Gunsteren WF, DiNola A, Haak JR. Molecular dynamics with coupling to an external bath. J Chem Phys. 1984;81: 3684–3690. doi:10.1063/1.448118

69. Parrinello M, Rahman A. Polymorphic transitions in single crystals: A new molecular dynamics method. J Appl Phys. 1981;52: 7182–7190. doi:10.1063/1.328693

70. Bussi G, Donadio D, Parrinello M. Canonical sampling through velocity rescaling. J Chem Phys. 2007;126: 014101. doi:10.1063/1.2408420

71. Darden T, York D, Pedersen L. Particle mesh Ewald: An N·log(N) method for Ewald sums in large systems. J Chem Phys. 1993;98: 10089–10092. doi:10.1063/1.464397

72. Hess B, Bekker H, Berendsen HJC, Fraaije JG. LINCS: a linear constraint solver for molecular simulations. J Comput Chem. 1997;18: 1463–1472. Available: https://onlinelibrary.wiley.com/doi/abs/10.1002/(SICI)1096-987X(199709)18:12%3C1463::AID-JCC4%3E3.0.CO;2-H

73. Humphrey W, Dalke A, Schulten K. VMD: visual molecular dynamics. J Mol Graph. 1996;14: 33–8, 27–8. doi:10.1016/0263-7855(96)00018-5

74. Patel JS, Quates CJ, Johnson EL, Ytreberg FM. Expanding the watch list for potential Ebola virus antibody escape mutations. PLoS One. 2019;14: e0211093. doi:10.1371/journal.pone.0211093

75. Yang J, Naik N, Patel JS, Wylie CS, Gu W, Huang J, et al. Predicting the viability of beta-lactamase: How folding and binding free energies correlate with beta-lactamase fitness. PLoS One. 2020;15: e0233509. doi:10.1371/journal.pone.0233509

76. Miller CR, Johnson EL, Burke AZ, Martin KP, Miura TA, Wichman HA, et al. Initiating a watch list for Ebola virus antibody escape mutations. PeerJ. 2016;4: e1674. doi:10.7717/peerj.1674

77. Schymkowitz J, Borg J, Stricher F, Nys R, Rousseau F, Serrano L. The FoldX web server: an online force field. Nucleic Acids Res. 2005;33: W382–8. doi:10.1093/nar/gki387

78. Delano, L W. The PyMOL Molecular Graphics System. http://www.pymol.org. 2002 [cited 13 Sep 2020]. Available: https://ci.nii.ac.jp/naid/10020095229/

79. Laemmli UK. Cleavage of Structural Proteins during the Assembly of the Head of Bacteriophage T4. Nature. 1970;227: 680–685. doi:10.1038/227680a0

80. Bradford MM. A rapid and sensitive method for the quantitation of microgram quantities of protein utilizing the principle of protein-dye binding. Anal Biochem. 1976;72: 248–254. doi:10.1016/0003-2697(76)90527-3

81. Zamboni N, Fendt S-M, Rühl M, Sauer U. (13)C-based metabolic flux analysis. Nat Protoc. 2009;4: 878–892. doi:10.1038/nprot.2009.58

82. Johnson CL, Pechonick E, Park SD, Havemann GD, Leal NA, Bobik TA. Functional genomic, biochemical, and genetic characterization of the *Salmonella pduO* gene, an ATP:cob(I)alamin adenosyltransferase gene. J Bacteriol. 2001;183: 1577–1584. doi:10.1128/JB.183.5.1577-1584.2001

83. Chowdhury C, Sinha S, Chun S, Yeates TO, Bobik TA. Diverse bacterial microcompartment organelles. Microbiol Mol Biol Rev. 2014;78: 438–468. doi:10.1128/MMBR.00009-14

84. Axen SD, Erbilgin O, Kerfeld CA. A taxonomy of bacterial microcompartment loci constructed by a novel scoring method. PLoS Comput Biol. 2014;10: e1003898. doi:10.1371/journal.pcbi.1003898

85. Zhang T, Ren H-F, Liu Y, Zhu B-L, Liu Z-P. A novel degradation pathway of chloroaniline in Diaphorobacter sp. PCA039 entails initial hydroxylation. World J Microbiol Biotechnol. 2010;26: 665–673. Available: https://idp.springer.com/authorize/casa?redirect_uri= https://link.springer.com/article/10.1007/s11274-009-0221-1&casa_token=I0i1UVru00kAAAAA:C9CnKeesGDMAuildgQ4z499DFVqOOOu36NTij-41QgogZkklxJ7HPaF-wB-539nUqi0PaGcarY05eiuwNw

86. Ortiz de Orué Lucana D, Hickey N, Hensel M, Klare JP, Geremia S, Tiufiakova T, et al. The Crystal Structure of the C-Terminal Domain of the *Salmonella enterica* PduO Protein: An Old Fold with a New Heme-Binding Mode. Front Microbiol. 2016;7: 1010. doi:10.3389/fmicb.2016.01010

87. Zarzycki J, Erbilgin O, Kerfeld CA. Bioinformatic characterization of glycyl radical enzyme-associated bacterial microcompartments. Appl Environ Microbiol. 2015;81: 8315–8329. doi:10.1128/AEM.02587-15

88. Bogel G, Schrempf H, Ortiz de Orué Lucana D. The heme-binding protein HbpS regulates the activity of the *Streptomyces reticuli* iron-sensing histidine kinase SenS in a redox-dependent manner. Amino Acids. 2009;37: 681–691. doi:10.1007/s00726-008-0188-5

89. Zou P, Groves MR, Viale-Bouroncle SD, Ortiz de Orué Lucana D. Crystallization and preliminary characterization of a novel haem-binding protein of *Streptomyces reticuli*. Acta Crystallogr Sect F Struct Biol Cryst Commun. 2008;64: 386–390. doi:10.1107/S1744309108008348

90. Groves MR, Lucana DOO. Adaptation to oxidative stress by Gram-positive bacteria: the redox sensing system HbpS-SenS-SenR from *Streptomyces reticuli*. Appl Microbiol Microb Biotechnol. 2010;1: 33–42. Available: https://www.researchgate.net/profile/Dario_Ortiz_de_Orue_Lucana/publication/266890113_Adaptation_to_oxidative_stress_by_Gram-positive_bacteria_The_redox_sensing_system_HbpS-SenS-SenR_from_Streptomyces_reticuli/links/544e32940cf29473161a4168/Adaptation-to-oxidative-stress-by-Gram-positive-bacteria-The-redox-sensing-system-HbpS-SenS-SenR-from-Streptomyces-reticuli.pdf

91. de Orué Lucana DO, Groves MR. The three-component signalling system HbpS--SenS--SenR as an example of a redox sensing pathway in bacteria. Amino Acids. 2009;37: 479–486. Available: https://idp.springer.com/authorize/casa?redirect_uri= https://link.springer.com/article/10.1007/s00726-009-0260-9&casa_token=KHKyI9Fuxw0AAAAA:g02raRt-YD65w_Hhy7XyYlr0Re9GaC0sOm6z6yj3dJ_T4JkV1IdadCV00wVY3vncP7lkJDMJ1DC9RMicAA

92. de Orué Lucana DO, Fedosov SN, Wedderhoff I. The extracellular heme-binding protein HbpS from the soil bacterium *Streptomyces reticuli* is an aquo-cobalamin binder. Journal of Biological. 2014. Available: http://www.jbc.org/content/289/49/34214.short

93. Ortiz de Orué Lucana D, Groves MR. The three-component signalling system HbpS-SenS-SenR as an example of a redox sensing pathway in bacteria. Amino Acids. 2009;37: 479–486. doi:10.1007/s00726-009-0260-9

94. de Orué Lucana DO, Bogel G, Zou P, Groves MR. The Oligomeric Assembly of the Novel Haem-Degrading Protein HbpS Is Essential for Interaction with Its Cognate Two-Component Sensor Kinase. J Mol Biol. 2009;386: 1108–1122. doi:10.1016/j.jmb.2009.01.017

95. Wedderhoff I, Kursula I, Groves MR, Ortiz de Orué Lucana D. Iron binding at specific sites within the octameric HbpS protects streptomycetes from iron-mediated oxidative stress. PLoS One. 2013;8: e71579. doi:10.1371/journal.pone.0071579

96. Donovan GT, Norton JP, Bower JM, Mulvey MA. Adenylate cyclase and the cyclic AMP receptor protein modulate stress resistance and virulence capacity of uropathogenic *Escherichia coli*. Infect Immun. 2013;81: 249–258. doi:10.1128/IAI.00796-12

97. McDonough KA, Rodriguez A. The myriad roles of cyclic AMP in microbial pathogens: from signal to sword. Nat Rev Microbiol. 2011;10: 27–38. doi:10.1038/nrmicro2688

98. Görke B, Stülke J. Carbon catabolite repression in bacteria: many ways to make the most out of nutrients. Nat Rev Microbiol. 2008;6: 613–624. doi:10.1038/nrmicro1932

99. Chan PF, O’Dwyer KM, Palmer LM, Ambrad JD, Ingraham KA, So C, et al. Characterization of a novel fucose-regulated promoter (PfcsK) suitable for gene essentiality and antibacterial mode-of-action studies in Streptococcus pneumoniae. J Bacteriol. 2003;185: 2051–2058. doi:10.1128/jb.185.6.2051-2058.2003

100. Mazel D, Pochet S, Marlière P. Genetic characterization of polypeptide deformylase, a distinctive enzyme of eubacterial translation. EMBO J. 1994;13: 914–923. Available: https://www.ncbi.nlm.nih.gov/pubmed/8112305

101. Teo JWP, Thayalan P, Beer D, Yap ASL, Nanjundappa M, Ngew X, et al. Peptide deformylase inhibitors as potent antimycobacterial agents. Antimicrob Agents Chemother. 2006;50: 3665–3673. doi:10.1128/AAC.00555-06

102. Margolis PS, Hackbarth CJ, Young DC, Wang W, Chen D, Yuan Z, et al. Peptide deformylase in Staphylococcus aureus: resistance to inhibition is mediated by mutations in the formyltransferase gene. Antimicrob Agents Chemother. 2000;44: 1825–1831. doi:10.1128/aac.44.7.1825-1831.2000

103. Cai Y, Chandrangsu P, Gaballa A, Helmann JD. Lack of formylated methionyl-tRNA has pleiotropic effects on *Bacillus subtilis*. Microbiology. 2017;163: 185–196. doi:10.1099/mic.0.000413

104. Tola YH, Fujitani Y, Tani A. Bacteria with natural chemotaxis towards methanol revealed by chemotaxis fishing technique. Biosci Biotechnol Biochem. 2019;83: 2163–2171. doi:10.1080/09168451.2019.1637715

105. Seip B, Innis CA. How Widespread is Metabolite Sensing by Ribosome-Arresting Nascent Peptides? J Mol Biol. 2016;428: 2217–2227. doi:10.1016/j.jmb.2016.04.019

106. Zhang Q, Li R, Li J, Shi H. Optimal Allocation of Bacterial Protein Resources under Nonlethal Protein Maturation Stress. Biophys J. 2018;115: 896–910. doi:10.1016/j.bpj.2018.07.021

107. Trézl L, Hullán L, Szarvas T, Csiba A, Szende B. Determination of endogenous formaldehyde in plants (fruits) bound to L-arginine and its relation to the folate cycle, photosynthesis and apoptosis. Acta Biol Hung. 1998;49: 253–263. Available: https://www.ncbi.nlm.nih.gov/pubmed/10526968

108. Li Z, Xu Y, Zhu H, Qian Y. Imaging of formaldehyde in plants with a ratiometric fluorescent probe. Chem Sci. 2017;8: 5616–5621. doi:10.1039/c7sc00373k

109. Nemecek-Marshall M, MacDonald RC, Franzen JJ, Wojciechowski CL, Fall R. Methanol Emission from Leaves (Enzymatic Detection of Gas-Phase Methanol and Relation of Methanol Fluxes to Stomatal Conductance and Leaf Development). Plant Physiol. 1995;108: 1359–1368. doi:10.1104/pp.108.4.1359

110. Hüve K, Christ MM, Kleist E, Uerlings R, Niinemets U, Walter A, et al. Simultaneous growth and emission measurements demonstrate an interactive control of methanol release by leaf expansion and stomata. J Exp Bot. 2007;58: 1783–1793. doi:10.1093/jxb/erm038

111. Jozefczuk S, Klie S, Catchpole G, Szymanski J, Cuadros-Inostroza A, Steinhauser D, et al. Metabolomic and transcriptomic stress response of *Escherichia coli*. Mol Syst Biol. 2010;6: 364. doi:10.1038/msb.2010.18

112. Bhat SV, Booth SC, Vantomme EAN, Afroj S, Yost CK, Dahms TES. Oxidative stress and metabolic perturbations in *Escherichia coli* exposed to sublethal levels of 2,4-dichlorophenoxyacetic acid. Chemosphere. 2015;135: 453–461. doi:10.1016/j.chemosphere.2014.12.035

113. Bazurto JV, Dearth SP, Tague ED, Campagna SR, Downs DM. Untargeted metabolomics confirms and extends the understanding of the impact of aminoimidazole carboxamide ribotide (AICAR) in the metabolic network of *Salmonella enterica*. Microb Cell Fact. 2017;5: 74–87. doi:10.15698/mic2018.02.613

114. Lenstra TL, Rodriguez J, Chen H, Larson DR. Transcription Dynamics in Living Cells. Annu Rev Biophys. 2016;45: 25–47. doi:10.1146/annurev-biophys-062215-010838

115. Yukihira D, Fujimura Y, Wariishi H, Miura D. Bacterial metabolism in immediate response to nutritional perturbation with temporal and network view of metabolites. Mol Biosyst. 2015;11: 2473–2482. doi:10.1039/c5mb00182j

116. Tsanas C, Stenby EH, Yan W. Calculation of simultaneous chemical and phase equilibrium by the method of Lagrange multipliers. Chem Eng Sci. 2017;174: 112–126. doi:10.1016/j.ces.2017.08.033

117. Lucana DO de O, de Orué Lucana DO, Zou P, Nierhaus M, Schrempf H. Identification of a novel two-component system SenS/SenR modulating the production of the catalase-peroxidase CpeB and the haem-binding protein HbpS in *Streptomyces reticuli*. Microbiology. 2005. pp. 3603–3614. doi:10.1099/mic.0.28298-0

118. Lucana DO de O, Schaa T, Schrempf H. The novel extracellular *Streptomyces reticuli* haem-binding protein HbpS influences the production of the catalase-peroxidase CpeB. Microbiology. 2004;150: 2575–2585. doi:10.1099/mic.0.27091-0

119. Minder AC, Fischer HM, Hennecke H, Narberhaus F. Role of HrcA and CIRCE in the heat shock regulatory network of *Bradyrhizobium japonicum*. J Bacteriol. 2000;182: 14–22. doi:10.1128/jb.182.1.14-22.2000

120. Schulz A, Schumann W. *hrcA*, the first gene of the *Bacillus subtilis dnaK* operon encodes a negative regulator of class I heat shock genes. J Bacteriol. 1996;178: 1088–1093. doi:10.1128/jb.178.4.1088-1093.1996

121. Masuda N, Church GM. Regulatory network of acid resistance genes in *Escherichia coli*. Mol Microbiol. 2003;48: 699–712. doi:10.1046/j.1365-2958.2003.03477.x

122. Gajiwala KS, Burley SK. HDEA, a periplasmic protein that supports acid resistance in pathogenic enteric bacteria. J Mol Biol. 2000;295: 605–612. doi:10.1006/jmbi.1999.3347

123. Patterson JA, He H, Folz JS, Li Q, Wilson MA, Fiehn O, et al. Thioproline formation as a driver of formaldehyde toxicity in *Escherichia coli*. Biochem J. 2020;477: 1745–1757. doi:10.1042/BCJ20200198

124. Piatkov KI, Vu TTM, Hwang C-S, Varshavsky A. Formyl-methionine as a degradation signal at the N-termini of bacterial proteins. Microb Cell Fact. 2015;2: 376–393. doi:10.15698/mic2015.10.231

125. Kim J-M, Seok O-H, Ju S, Heo J-E, Yeom J, Kim D-S, et al. Formyl-methionine as an N-degron of a eukaryotic N-end rule pathway. Science. 2018;362. doi:10.1126/science.aat0174

126. Krysztofinska EM, Evans NJ, Thapaliya A, Murray JW, Morgan RML, Martinez-Lumbreras S, et al. Structure and Interactions of the TPR Domain of Sgt2 with Yeast Chaperones and Ybr137wp. Front Mol Biosci. 2017;4: 68. doi:10.3389/fmolb.2017.00068

127. Kalyuzhnaya MG, Korotkova N, Crowther G, Marx CJ, Lidstrom ME, Chistoserdova L. Analysis of gene islands involved in methanopterin-linked C1 transfer reactions reveals new functions and provides evolutionary insights. J Bacteriol. 2005;187: 4607–4614. doi:10.1128/JB.187.13.4607-4614.2005

128. Chen FY-H, Jung H-W, Tsuei C-Y, Liao JC. Converting *Escherichia coli* to a Synthetic Methylotroph Growing Solely on Methanol. Cell. 2020;182: 933–946.e14. doi:10.1016/j.cell.2020.07.010

129. Sievers F, Higgins DG. Clustal Omega, accurate alignment of very large numbers of sequences. Methods Mol Biol. 2014;1079: 105–116. doi:10.1007/978-1-62703-646-7_6

130. Rice P, Longden I, Bleasby A. EMBOSS: the European Molecular Biology Open Software Suite. Trends Genet. 2000;16: 276–277. doi:10.1016/s0168-9525(00)02024-2

131. Nayak DD, Marx CJ. Genetic and phenotypic comparison of facultative methylotrophy between *Methylobacterium extorquens* strains PA1 and AM1. PLoS One. 2014;9: e107887. doi:10.1371/journal.pone.0107887

