## Supplemental Figures for "EfgA is a conserved formaldehyde sensor that halts bacterial translation in response to elevated formaldehyde"

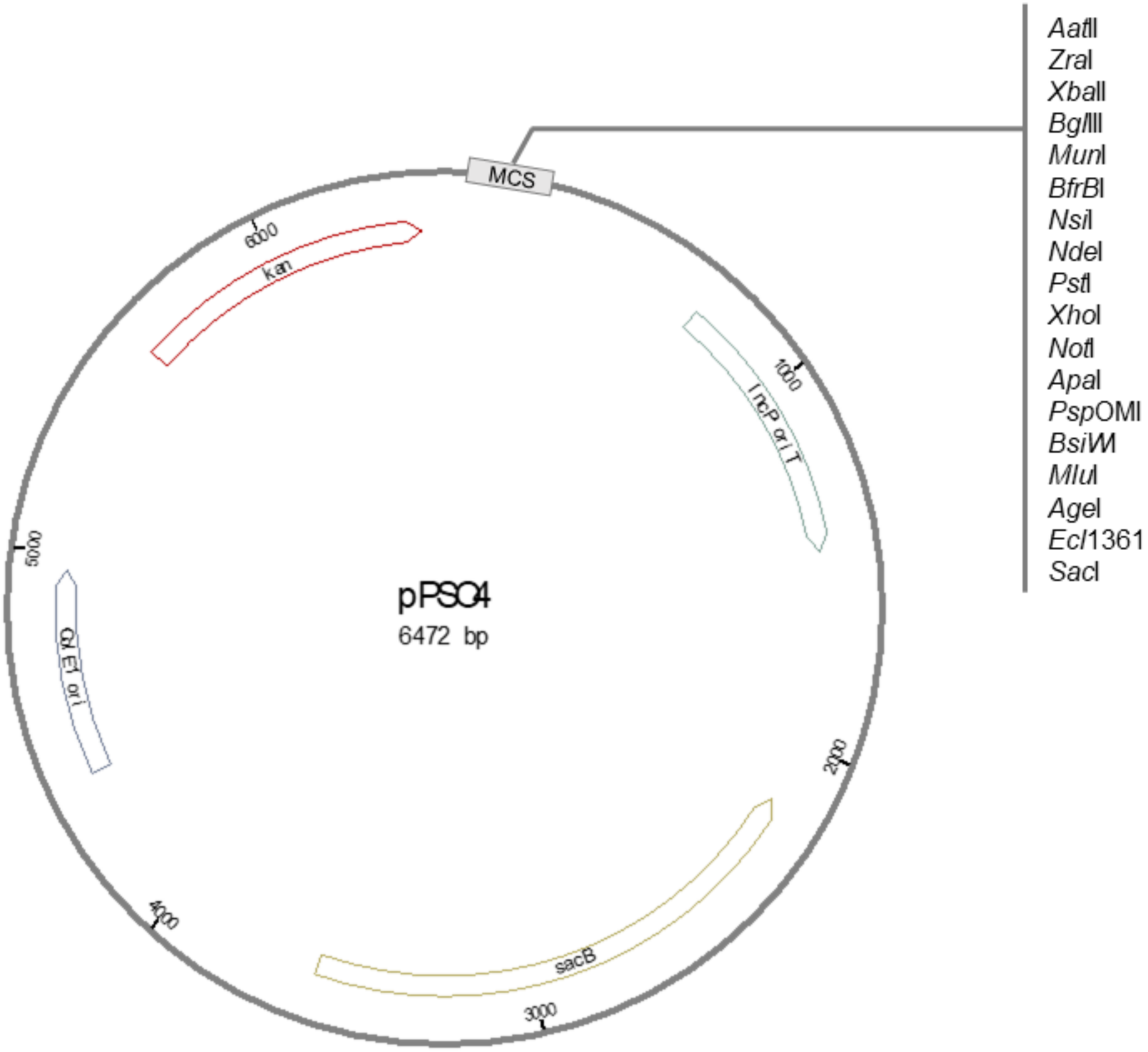


**Figure S1. Selective regime for the evolution of *M. extorquens*.**
*M. extorquens* was experimentally evolved in Hypho liquid media for 150 generations. Initial growth conditions relied on methanol as a sole source of carbon and energy. At each transfer (~ 6 generations), methanol concentrations were decreased and formaldehyde was introduced into the growth media at increasing concentrations. By generation 35, formaldehyde was the only carbon/energy source present. Formaldehyde concentrations continued to be gradually increased until generation 60 when it reached 20 mM. Selective pressure was sustained at 20 mM formaldehyde until the experiment was completed at 150 generations.

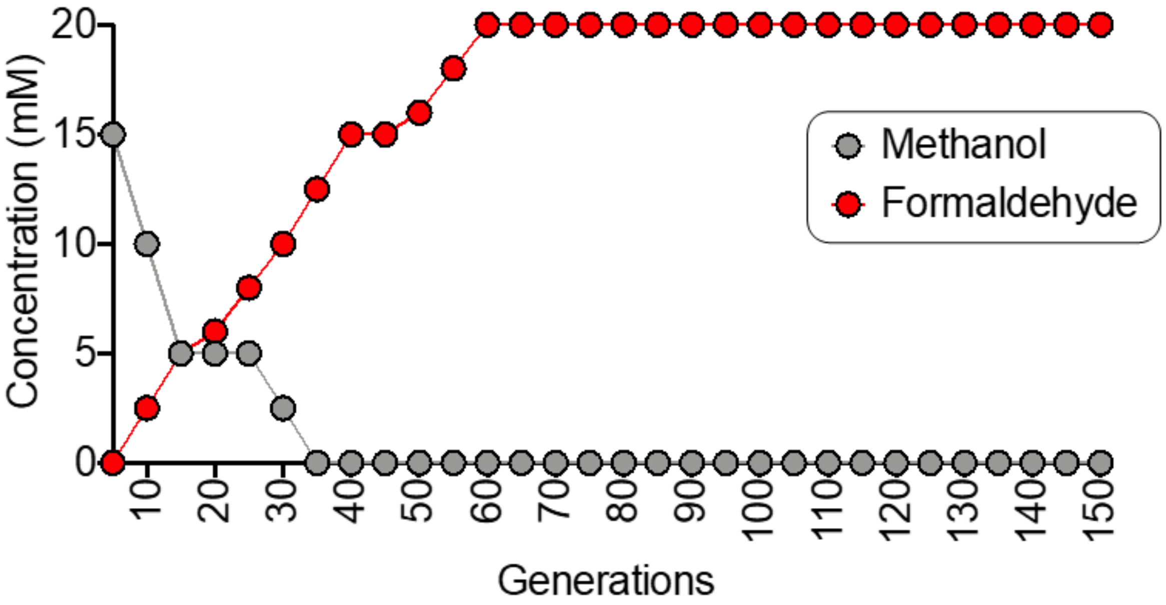


**Figure S2. Broad-host-range allelic exchange vector pPS04.** Plasmid map of pPS04 [GenBank: pending] showing the key features including *kan* (encodes kanamycin resistance), multiple cloning site (MCS) containing a number of single-cutting restriction sites, IncP *oriT* (origin of conjugal transfer), *sacB* (encodes levansucrase for sucrose sensitivity), ColE1 *ori* (high-copy origin of replication for *E. coli*).

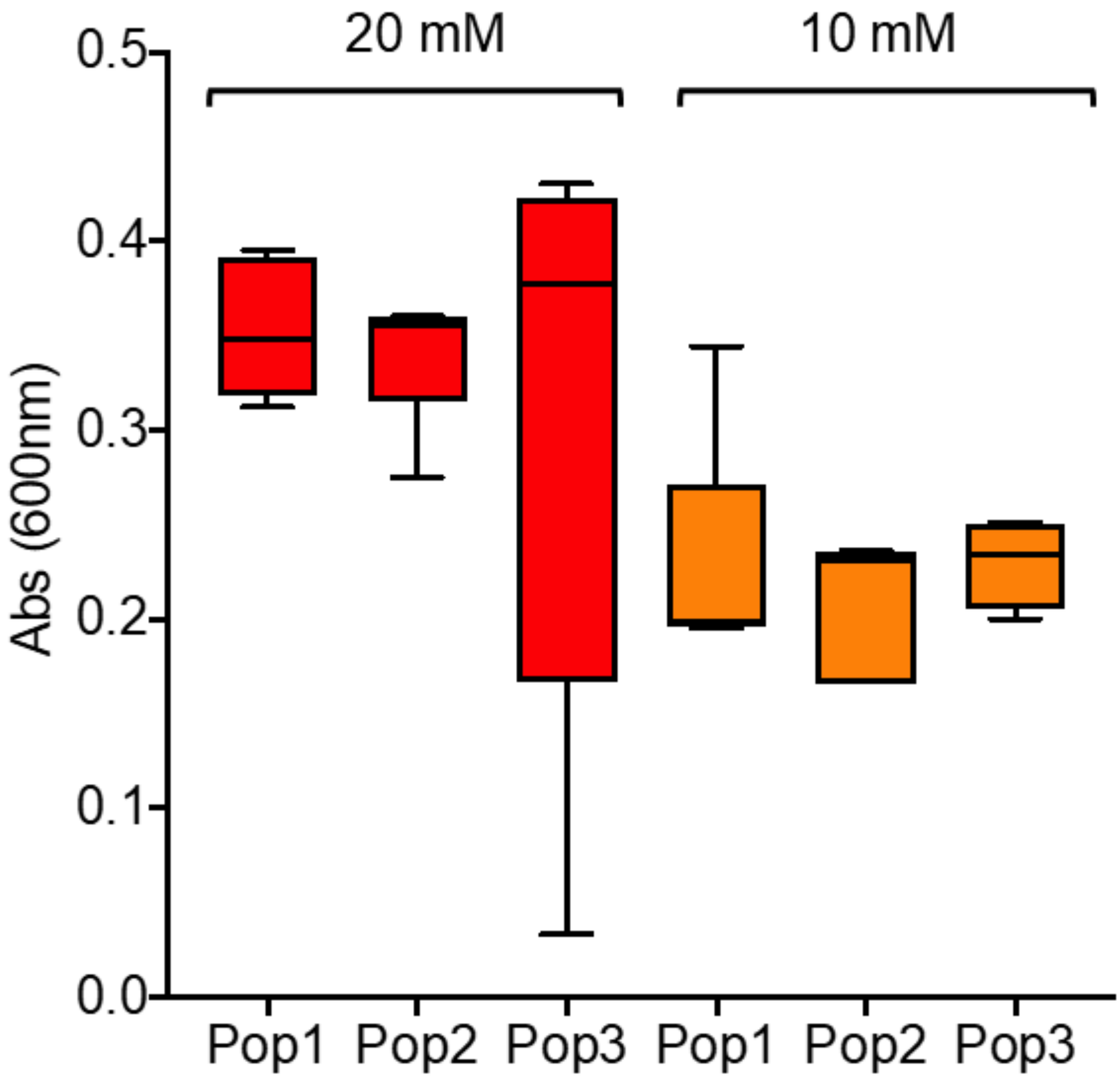


**Figure S3. Individual isolates from formaldehyde-evolved populations grow on formaldehyde up to 20 mM.**
Five isolates from each of the final formaldehyde-evolved populations (CM3031-CM3045, Table 1) were grown in minimal Hypho medium with 20 mM or 10 mM formaldehyde for 48 h. Plot whiskers indicate the minimum and maximum values.

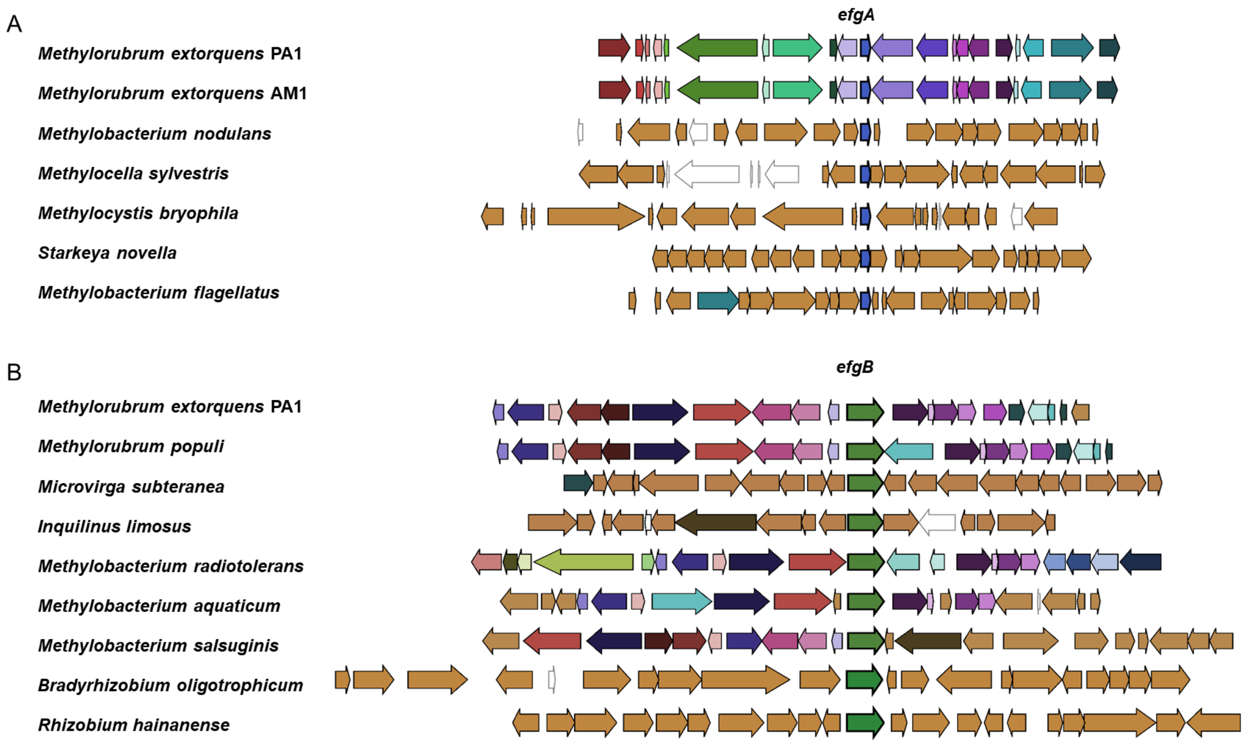


**Figure S4. Genomic context of *efgA* and *efgB* in other organisms.**
Gene neighborhoods from fully assembled genomes are aligned by A) *efgA* (blue) or B) *efgB* (green) and include 10 flanking genes on either side. Regions of homology are indicated by identical colors among the genomes within each panel. In both panels, regions lacking homology are shown in tan.

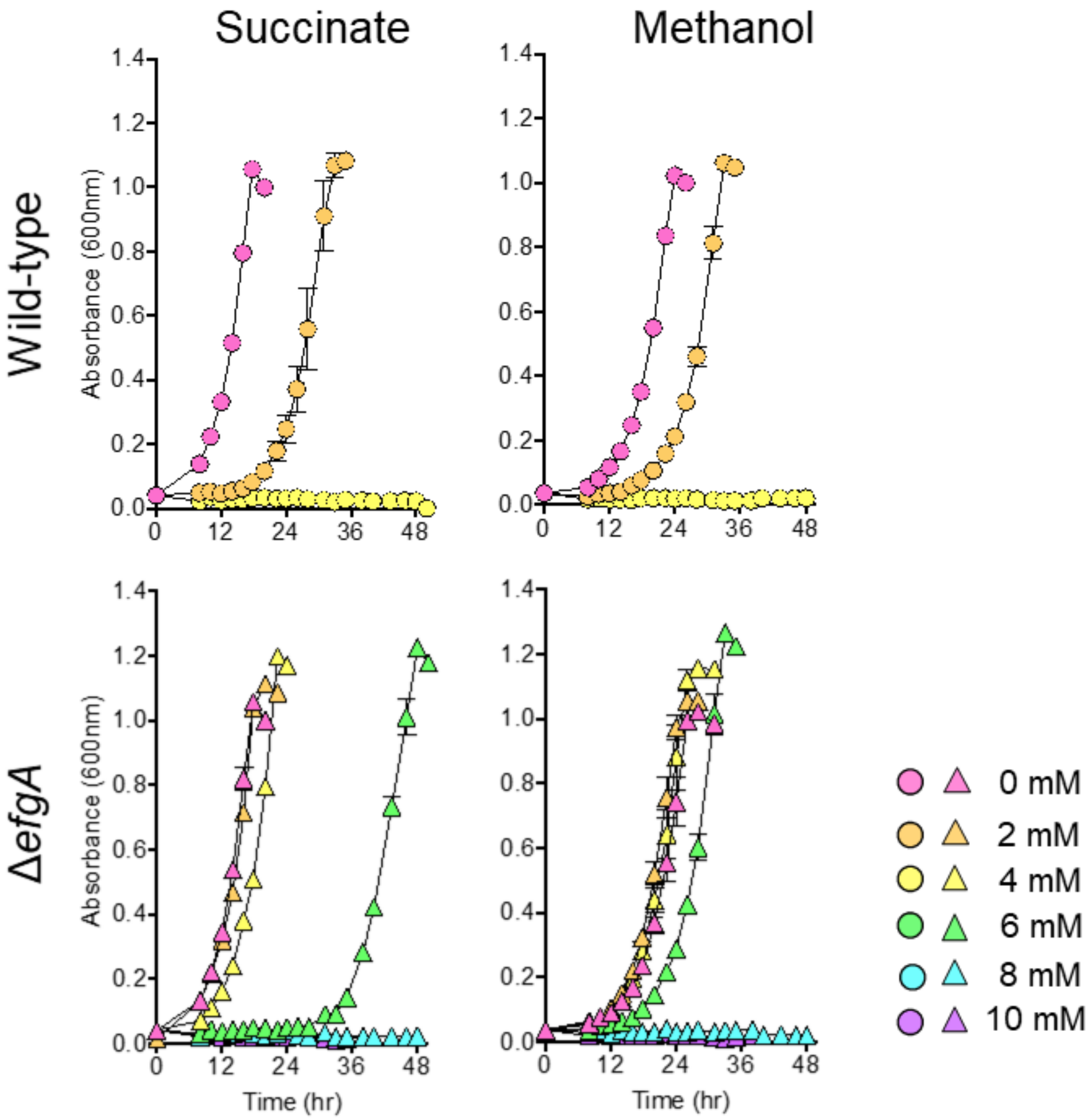


**Figure S5. The absence of *efgA* increases formaldehyde resistance during growth on alternative compounds.**
Wild-type (CM2730, circles, upper panels) and the Δ*efgA* mutant (CM3745, triangles, lower panels) were grown in liquid MP medium with succinate (left panels) or methanol (right panels) provided as the primary carbon source. Additionally, 0, 2, 4, 6, 8, or 10 mM exogenous formaldehyde was provided as a stressor. Error bars represent the standard error of the mean of three biological replicates.

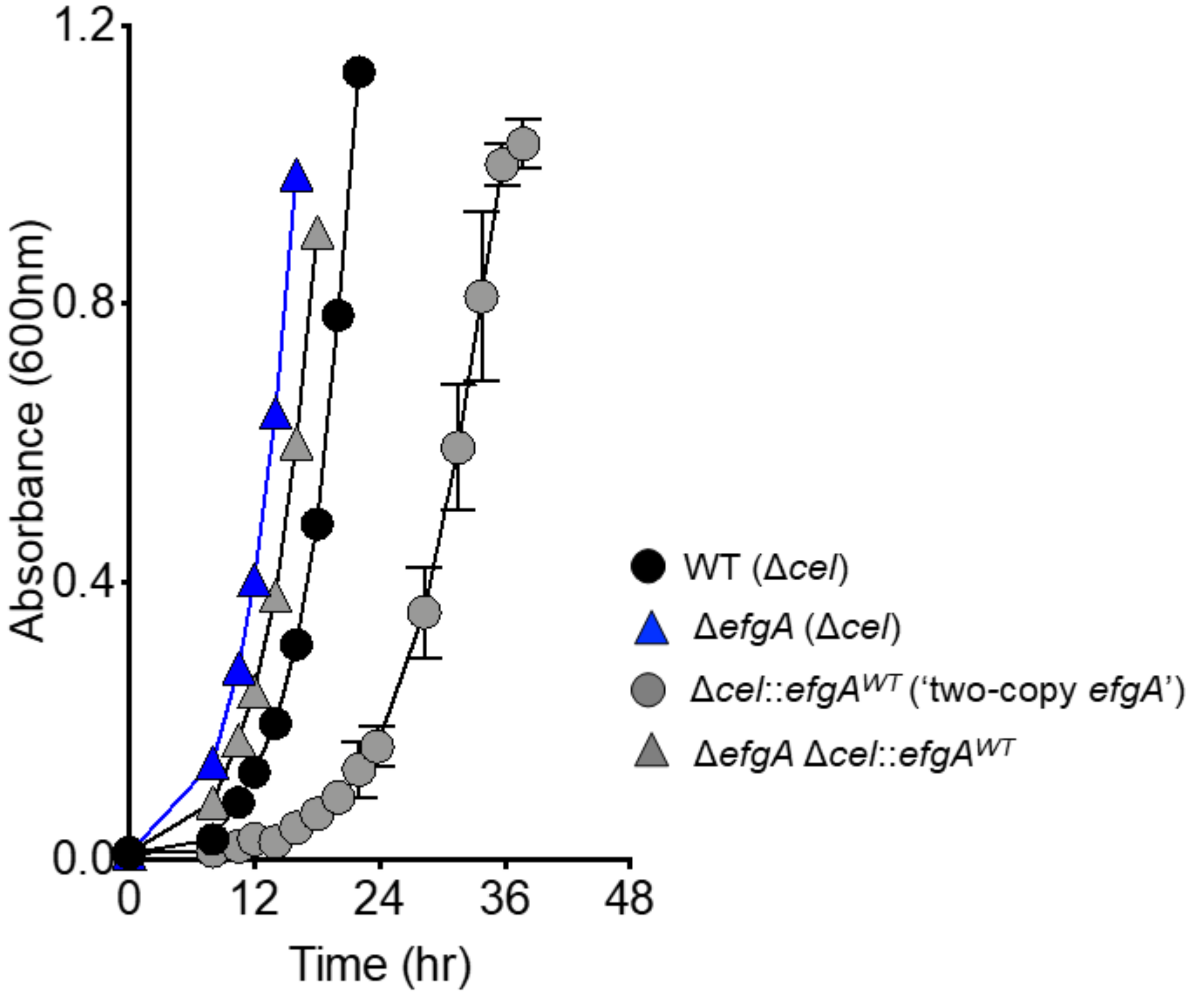


**Figure S6. Increasing the *efgA* gene dose sensitizes *M. extorquens* to formaldehyde.** The wild-type (black circles), Δ*efgA* mutant (blue triangles), two-copy *efgA* mutant (gray circles), and Δ*efgA*+chromosomal *efgA* complement (gray triangles) were grown in liquid MP medium with 3.5 mM succinate and 2 mM formaldehyde. Error bars represent the standard error of mean of three biological replicates. All strains were derived from WT, which is Δ*cel*; the second copy of *efgA* was introduced at the Δ*cel* locus.

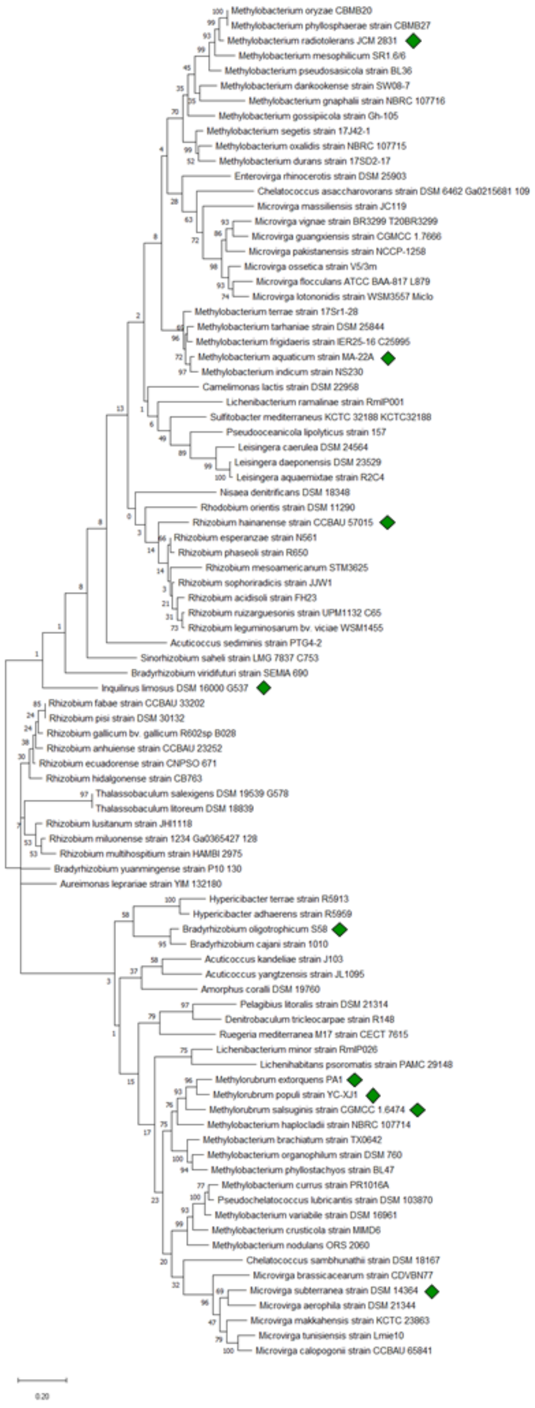


**Figure S7. Phylogenetic analysis of the EfgB indicates close homologs are in other members of the Rhizobiales with diverse physiologies.** The evolutionary relationship of *efgB* was compared to genes with 65-90 % identity via maximum likelihood. Bootstrap values are shown at nodes and branch lengths reflect the indicate substitutions per nucleotide. The green diamonds represent members whose genomic context is illustrated in Fig S4.

The phylogenetic data is available at TreeBASE (http://purl.org/phylo/treebase/phylows/study/TB2:S27073).

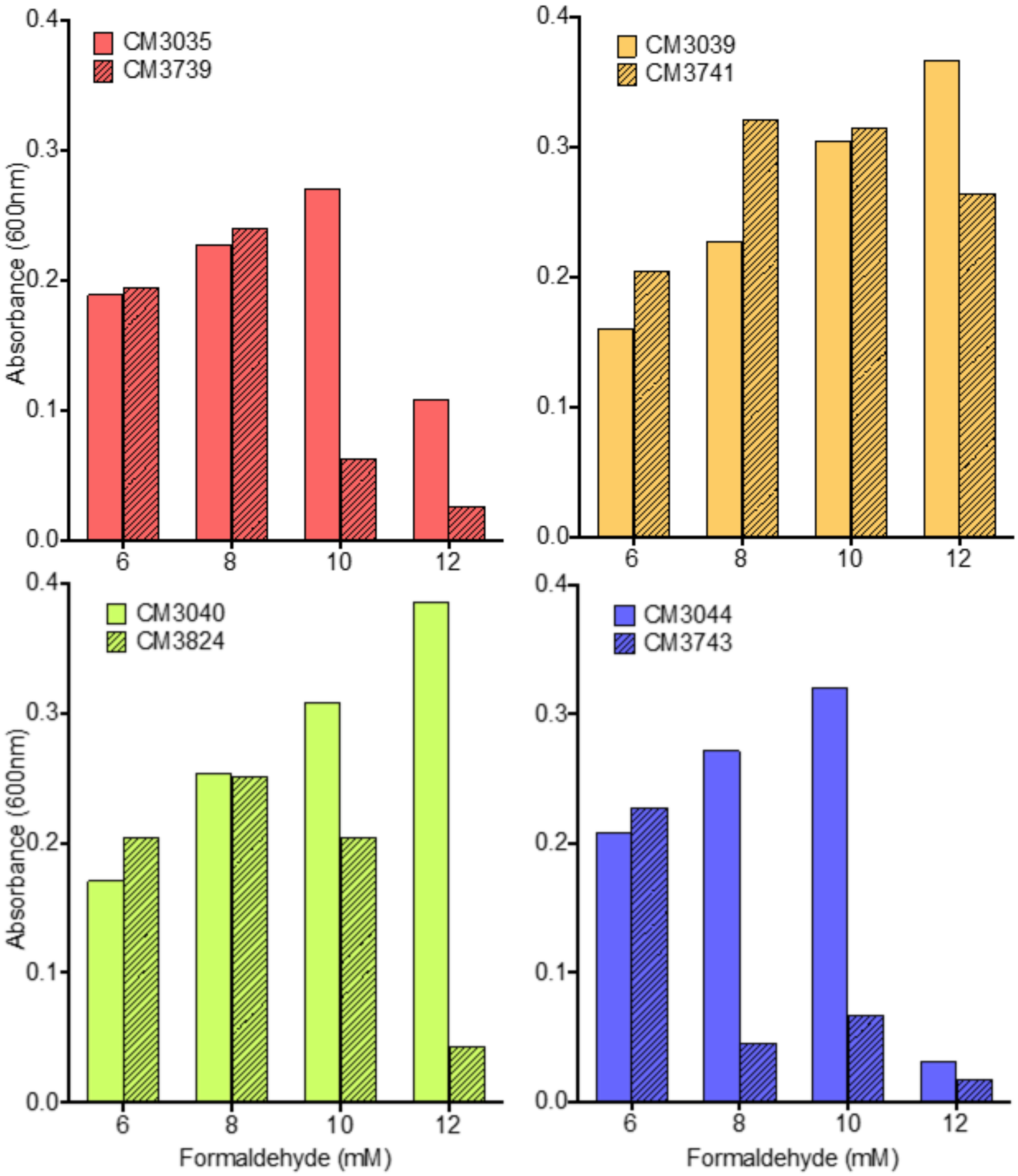


**Figure S8. *efgB* activity is needed for enhanced formaldehyde resistance in evolved isolates.** Formaldehyde growth of evolved isolates CM3035 (pink), CM3039 (orange), CM3040 (green), and CM3044 (violet) was measured in liquid MP medium with 6, 8, 10, or 12 mM exogenous formaldehyde provided as a sole source of carbon and energy. Replacing evolved beneficial *efgB* alleles with Δ*efgB* resulted in otherwise isogenic strains CM3739, CM3741, CM3824, and CM3743 (hatched bars) failed to utilize formaldehyde at higher concentrations. Data are representative of trends observed in multiple experiments.

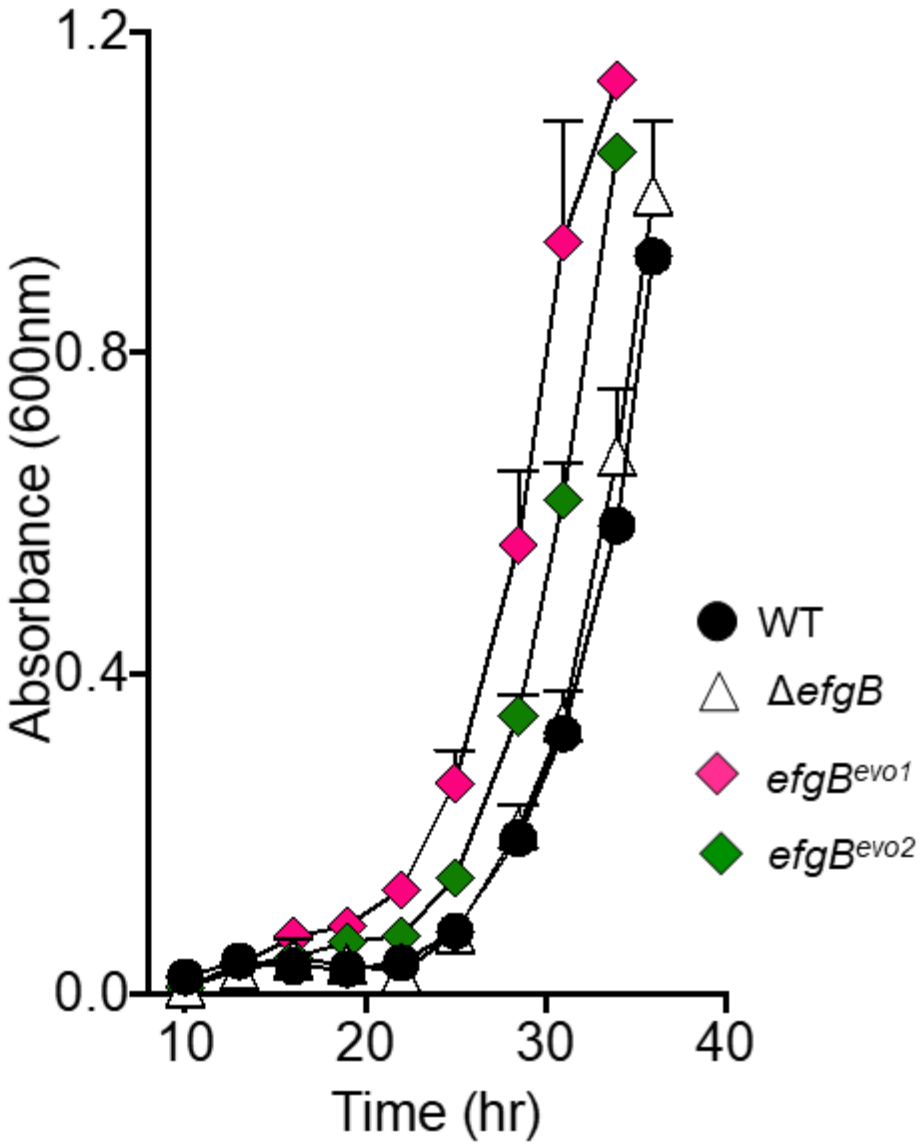


**Figure S9.** ***efgB***$^{evo}$ **alleles increase resistance to low concentrations of formaldehyde even when EfgA is functional.**
Growth of wild-type (CM2730, circles), Δ*efgB* (CM3737, triangles) and *efgB*$^{evo1}$ (CM3783, pink diamonds) and *efgB*$^{evo2}$ mutant (CM3837, green diamonds) was quantified in liquid MP medium (methanol) containing 2 mM formaldehyde. Error bars represent the standard error of the mean for three biological replicates.

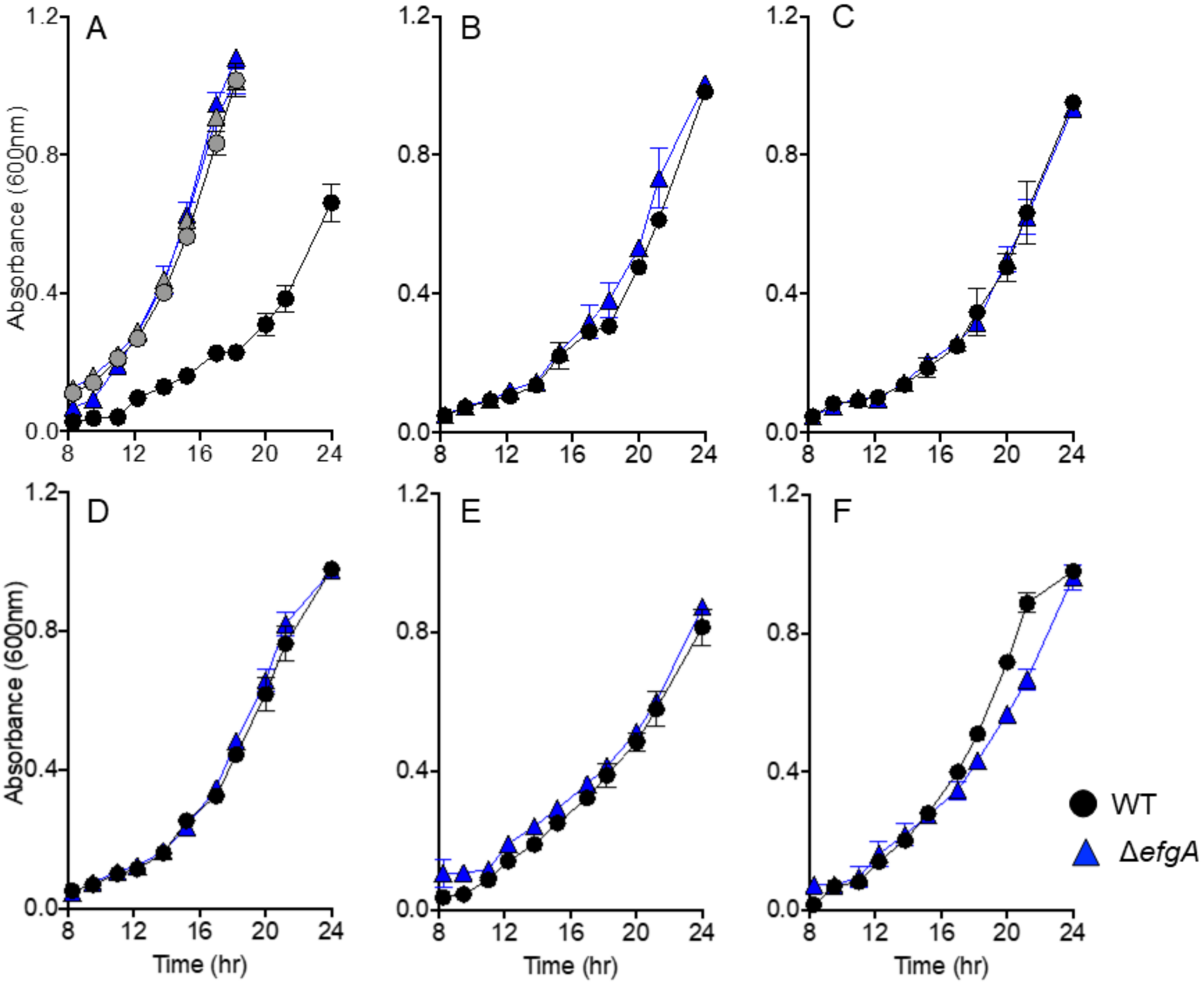


**Figure S10. The Δ*efgA* mutant does not have additional resistance to alternative aldehydes.** Growth of wild-type (CM2730, circles) and the Δ*efgA* mutant (CM3745, triangles) was quantified in liquid MP medium (succinate) containing no aldehydes (panel A, gray symbols). Additionally, growth of wild-type (black circles) and the Δ*efgA* mutant (blue triangles) was quantified in the same medium with the addition of (A) 2 mM formaldehyde, (B) 1.25 mM acetaldehyde, (C) 2.5 mM butyraldehyde, (D) 2.5 mM propionaldehyde, (E) 1.25 mM glyoxal, and (F) 0.157 mM glutaraldehyde. Error bars represent the standard error of mean of three biological replicates.

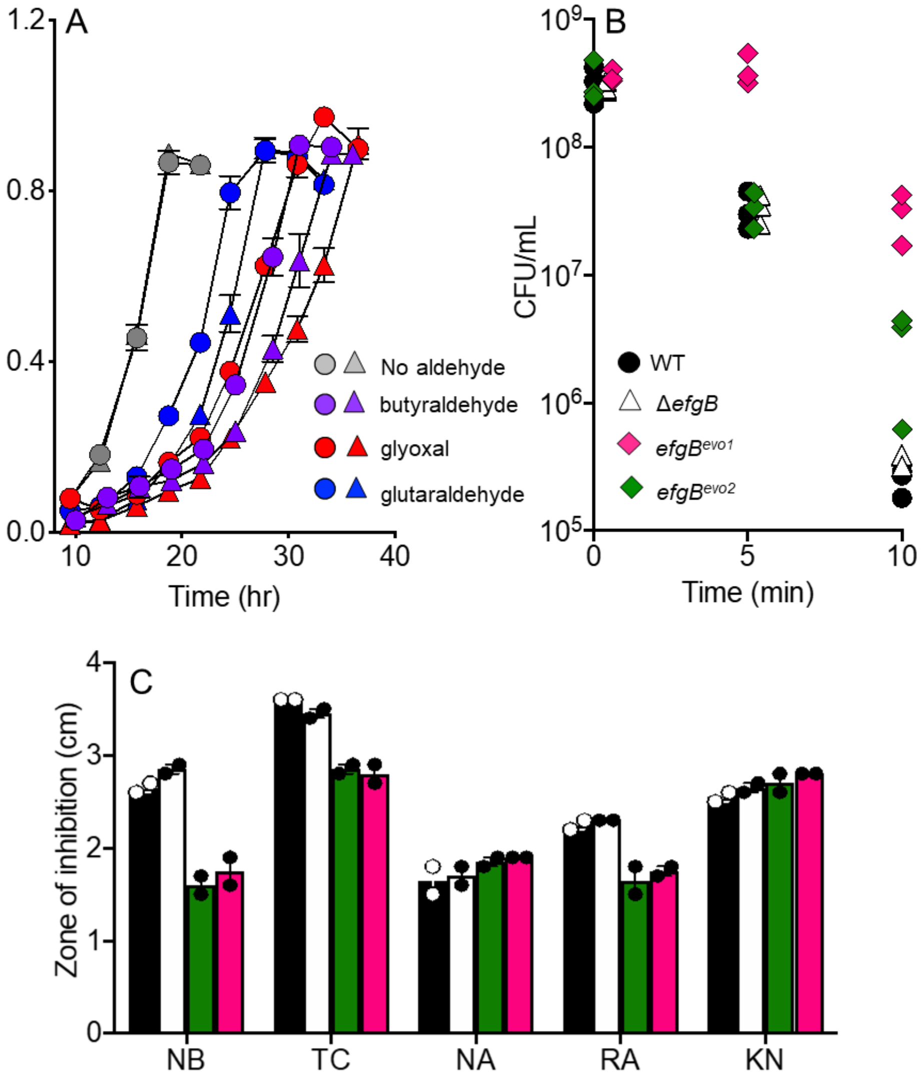


**Figure S11. *efgB^evo* alleles confer resistance to a variety of stressors.**
(A) Growth of wild-type (CM2730, circles) and the Δ*efgB* mutant (CM3737, triangles) were quantified in methanol medium with the addition of no aldehydes (gray symbols), 2.5 mM butyraldehyde (purple symbols), 1.25 mM glyoxal (red symbols), and 0.157 mM glutaraldehyde (blue symbols).
(B) Viability of wild-type (CM2730, circles), Δ*efgB* (CM3737, triangles) and *efgB^evo1* (CM3783, pink diamonds) and *efgB^evo2* mutant (CM3837, green diamonds) was assayed when culture tubes were submerged in a 55 °C water bath for 0, 5, or 10 m.
Error bars represent the standard error of the mean for three biological replicates.
(C) Disc-diffusion assays were performed by placing antibiotic-impregnated discs upon soft agar overlays of *M. extorquens* on solid MP media (15 mM succinate). The zones of inhibition showed that *efgB^evo1* (CM3783, pink) and *efgB^evo2* mutants (CM3837, green) are more resistant to multiple antibiotics than the wild-type (CM2730, black) and the Δ*efgB* mutant (CM3737, white). Abbreviations: NB, novobiocin; TC, tetracycline; NA, nalidixic acid; RA, rifampicin; KN, kanamycin.

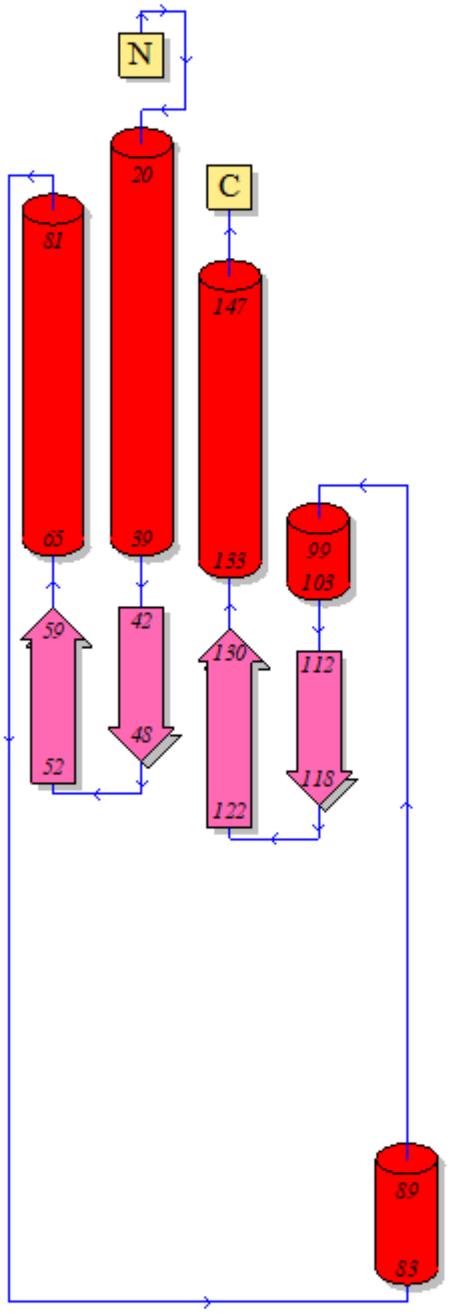


**Figure S12. Protein topology of EfgA.**
The secondary structural elements of EfgA and their relative positions are shown. The peptide chain begins at the N-terminus ('N') and proceeds through the C-terminus ('C'); the directionality is indicated by the small blue arrows. Cylinders represent α-helices and the wide arrows represent the β-strands. Residue numbers that begin and end each element are noted.

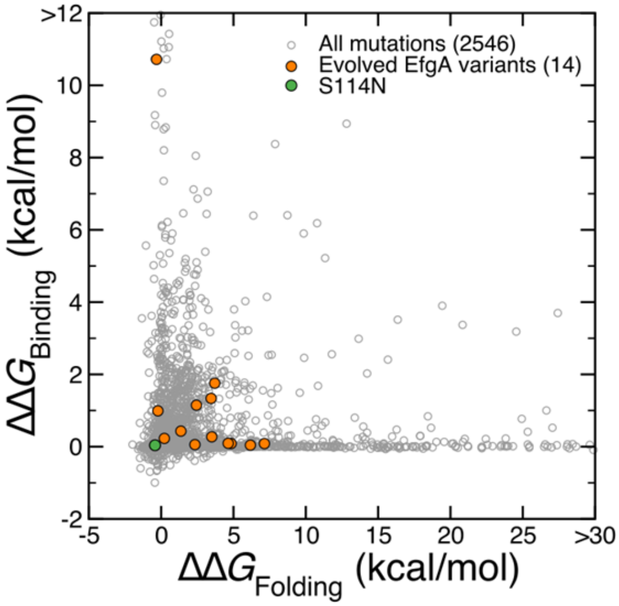


| Mutation | ΔΔG Folding (kcal/mol) | ΔΔG Binding (kcal/mol) |
|---|---|---|
| H2Q | -0.22545 | 0.98970 |
| T4I | 1.36372 | 0.43060 |
| T24P | 4.83795 | 0.08607 |
| G49D | -0.32094 | 10.71873 |
| T60I | 2.33365 | 0.05671 |
| F64S | 3.70297 | 1.75553 |
| G65D | 3.45529 | 1.33689 |
| G65S | 2.43884 | 1.14712 |
| S75P | 4.63705 | 0.09160 |
| L91H | 3.50223 | 0.27134 |
| M107I | 0.21980 | 0.22777 |
| G109C | 7.14074 | 0.08397 |
| A110P | 6.17139 | 0.04177 |
| S114N | -0.42634 | 0.03889 |

**Figure S13. Evolved *efgA* alleles have a variety of predicted folding and binding stabilities.**

We used our MD+FoldX approach [76] to predict the effect of all possible 19 mutations at each amino acid site on monomers or tetramer formation. (A) The distribution of ΔΔG values associated to monomer folding and tetramer binding for all possible nonsynonymous mutations (2546) of *efgA* are shown as grey circles. Orange circles indicate the location of the experimentally observed mutations (14) within the distribution. The green circle indicates the location of the S114N mutation. (B) A table of the 14 experimentally observed amino acid substitutions with their ΔΔG folding and ΔΔG binding values listed (in kcal/mol). Of these, 10 mutations increased the folding free energy of the monomer, suggesting they decreased the monomer stability and one was predicted to significantly increase binding free energy associated to tetramer formation, suggesting it destabilized the oligomeric assembly.

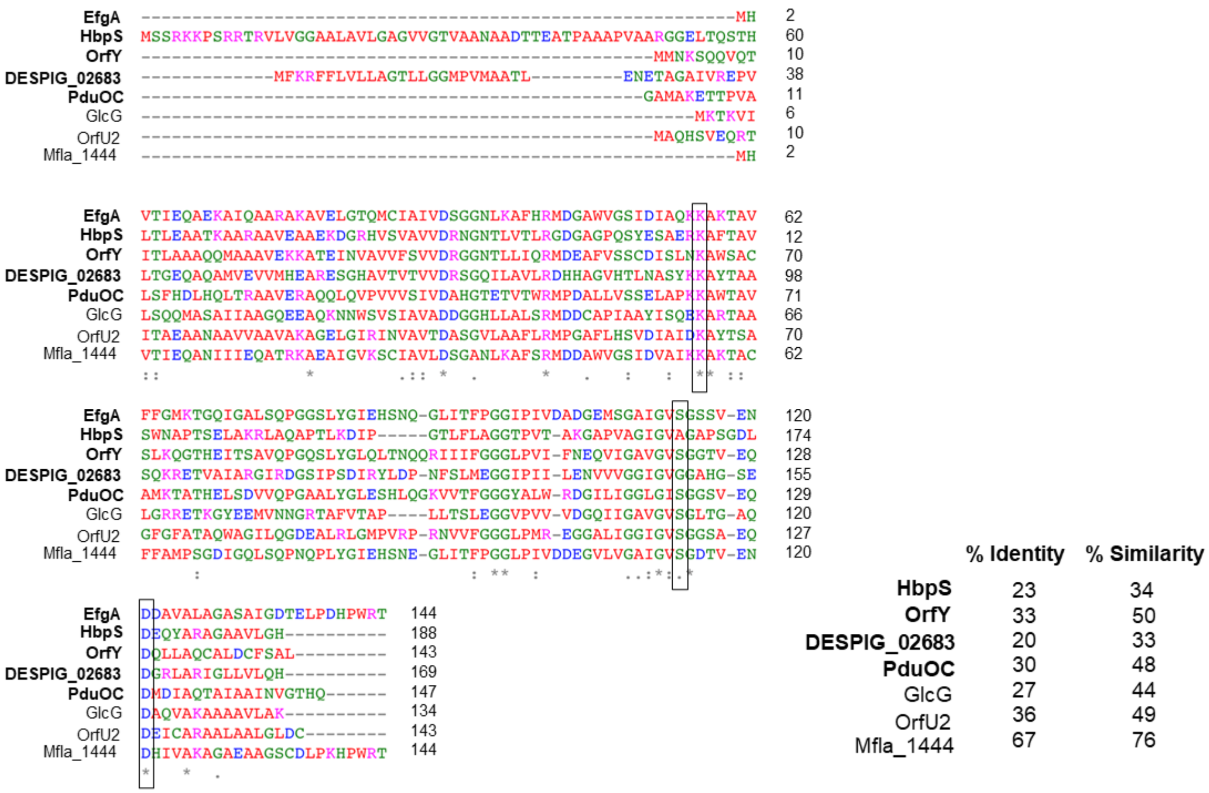


**Figure S14. Formaldehyde-binding residues are largely conserved in DUF336 homologs.** A Clustal Omega [129] alignment was performed with EfgA, structural DUF336 homologs (bold) and additional homologs referenced in the text. Conservation of residues is indicated when identical (*), strongly similar (:), or weakly similar (.). Small-hydrophobic residues (less Y) are in red (AVFPMILW), acidic residues are in blue (DE), basic residues are in magenta (RHK), and hydroxly + sulfhydryl + amine + G residues are in green (STYHCNGQ). Boxes indicate residues that correspond to those implicated in formaldehyde binding in EfgA (K57, S114, and D121). Pairwise comparisons between EfgA and each homolog were performed with EMBOSS Needle [130] to ascertain % identities and % similarities. For PduO, only the C-terminal DUF336 domain (PDB:5CX7) was included.

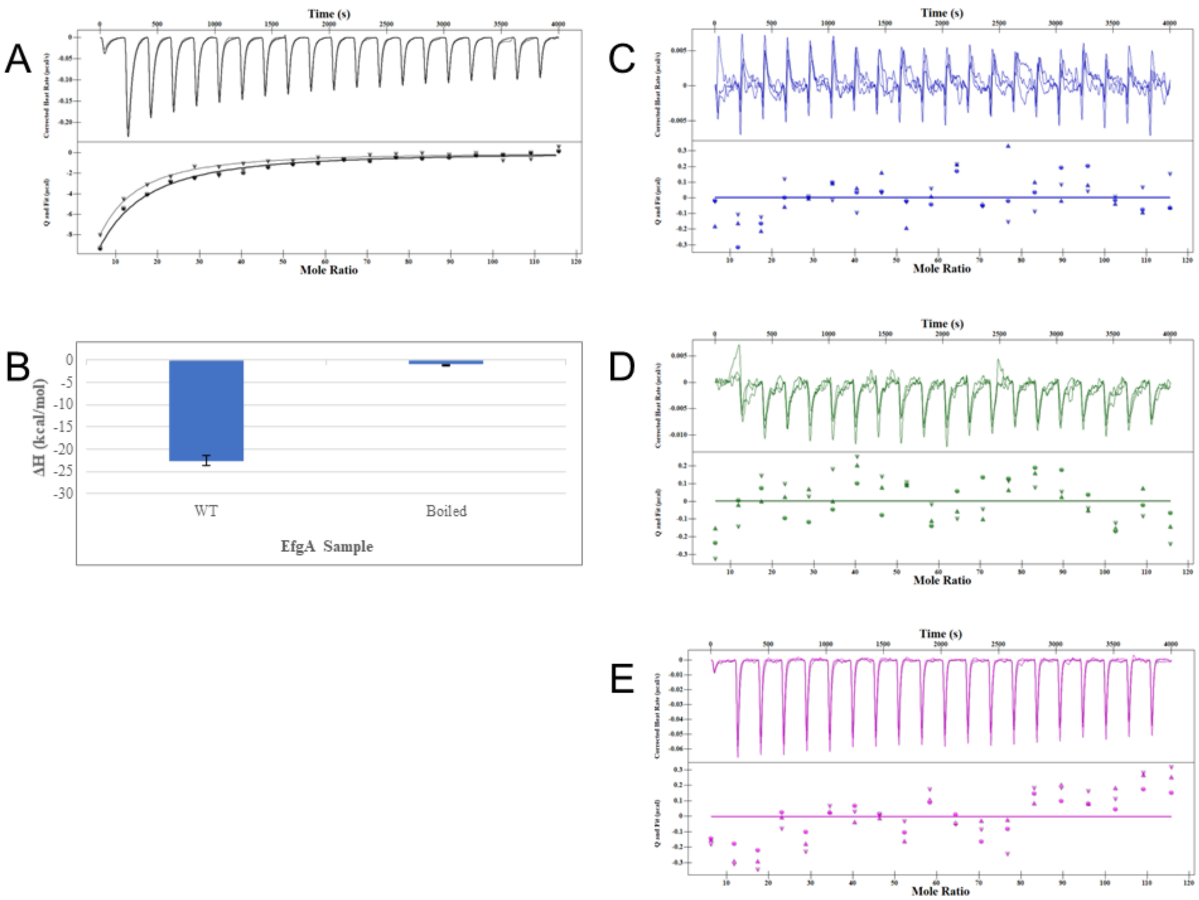


**Figure S15. Replicates of microscale isothermal calorimetry assays of EfgA:ligand binding.** The binding isotherms represented as heat change (µJ/s) upon injection over time are in the top portion of the split graphs, with independent binding modelling on the bottom portion. Binding observed with 50 µM EfgA (A) and 2 µL injections of 25 mM formaldehyde (in PBS). Binding observed with 50 µM EfgA and 2 µL injections of methanol (C), formate (D), acetaldehyde (E). Data are experimental replicates (*n* = 3) performed with protein from three independent purifications.

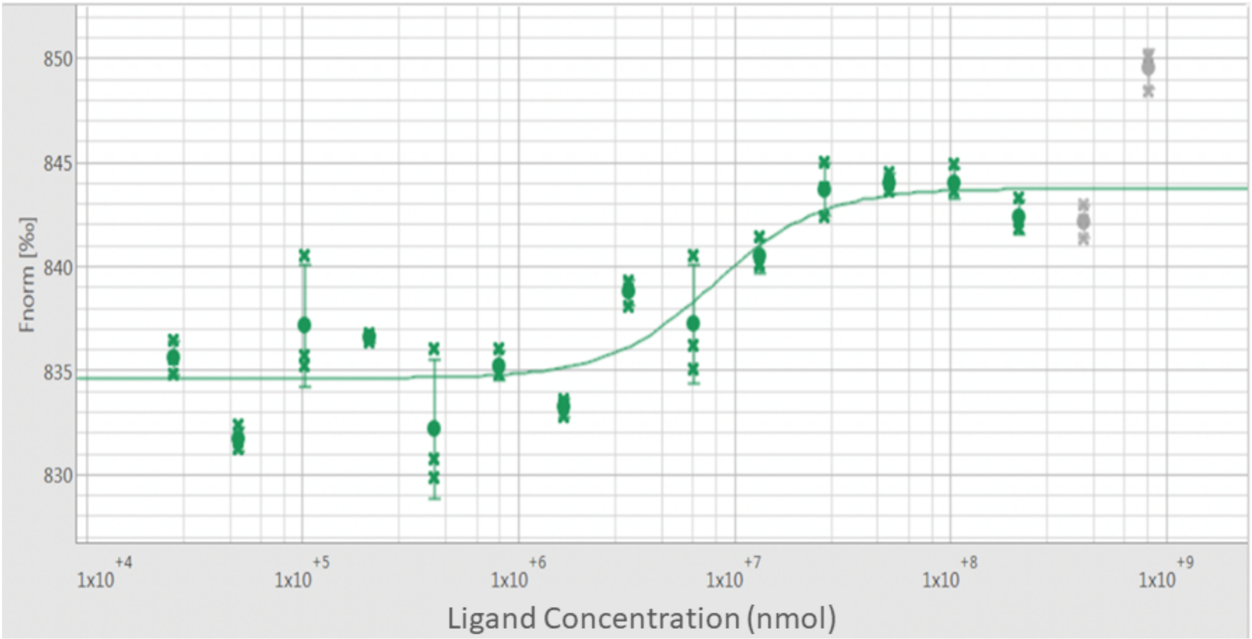


**Figure S16: Microscale thermophoresis indicates EfgA binds formaldehyde.** The dose response curve of EfgA to formaldehyde is represented by the difference in normalized fluorescence ($F_{norm}$ [‰]) for analysis of thermophoresis across formaldehyde concentrations with 20 nM EfgA. The $K_d$ is fitted to 8.01± 3.5 mM. Data represent (*n* = 3) MST measurements.

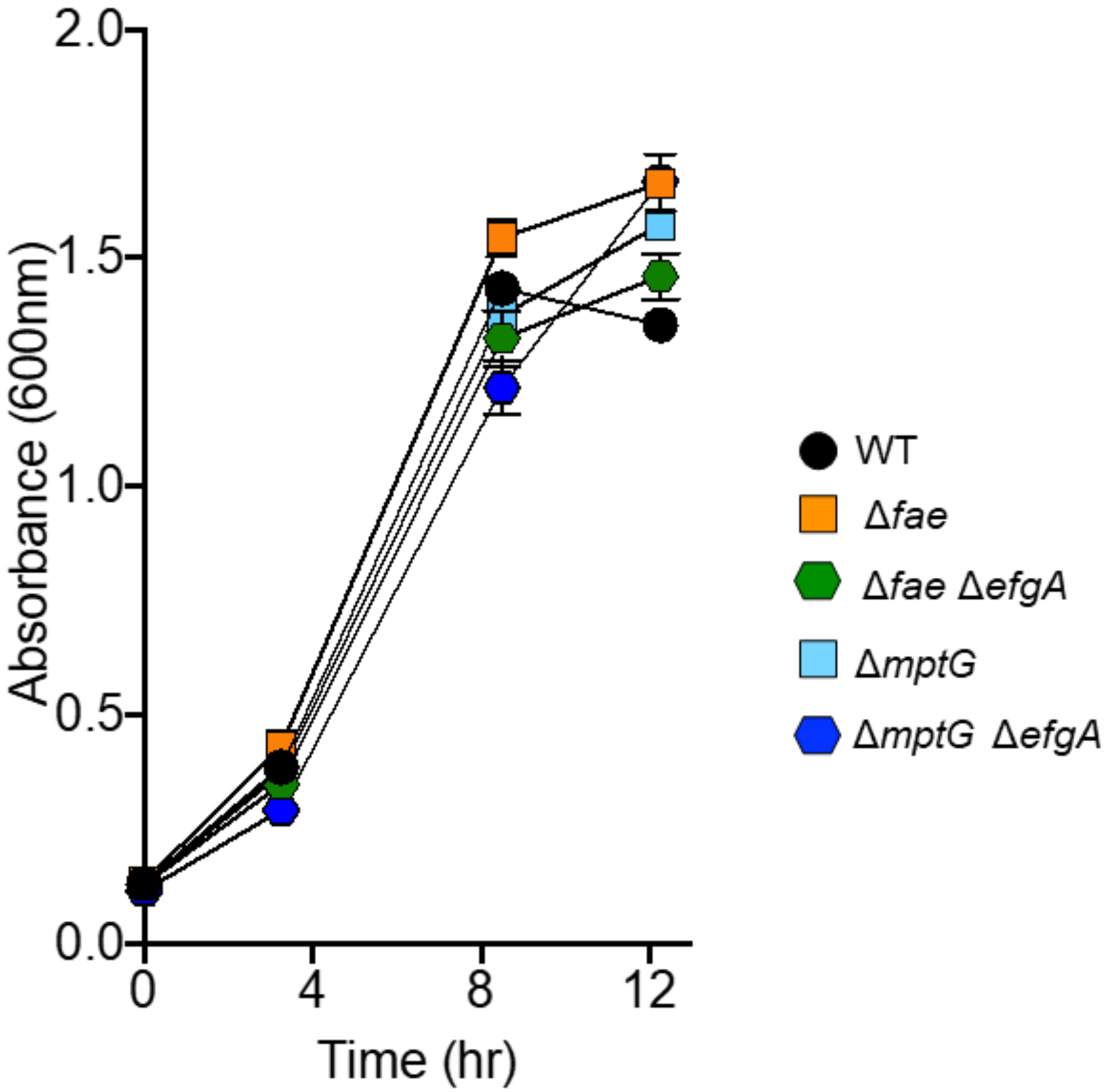


**Figure S17. Growth of methanol sensitive mutants is comparable to wild-type during growth on succinate.**

Wild-type and mutant strains were grown in liquid MP medium (succinate). Strains represented are wild-type (black circles), Δ*fae* (CM3753, orange squares), Δ*mptG* (CM4765, light blue squares), Δ*efgA* Δ*fae* (CM3421-5, green hexagons), and Δ*efgA* Δ*mptG* mutants (CM3440-13, blue hexagons). Error bars represent the standard error of the mean for three biological replicates.

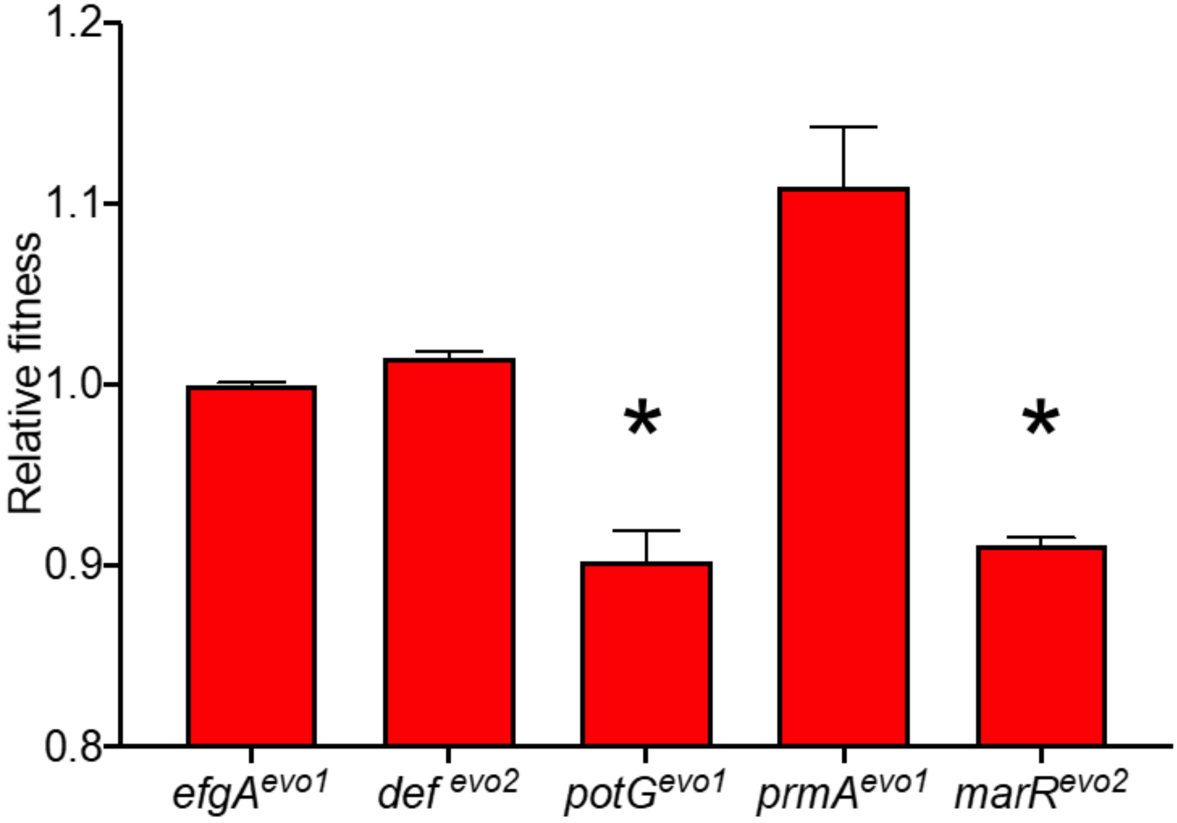


**Figure S18. Relative fitness of *efgA* and alleles at other loci.** Strains with evolved beneficial alleles that independently conferred formaldehyde growth were assessed for fitness in media containing 5 mM formaldehyde. Relative fitness values were determined via competition experiments against a common fluorescently-tagged reference strain. Fitness values for each strain, relative to the *efgA*$^{evo1}$ mutant, are presented as bars representing mean +/- SEM (n=3 biological replicates). Statistical significance was determined by an unpaired Student's *t* test (*, p<0.05).

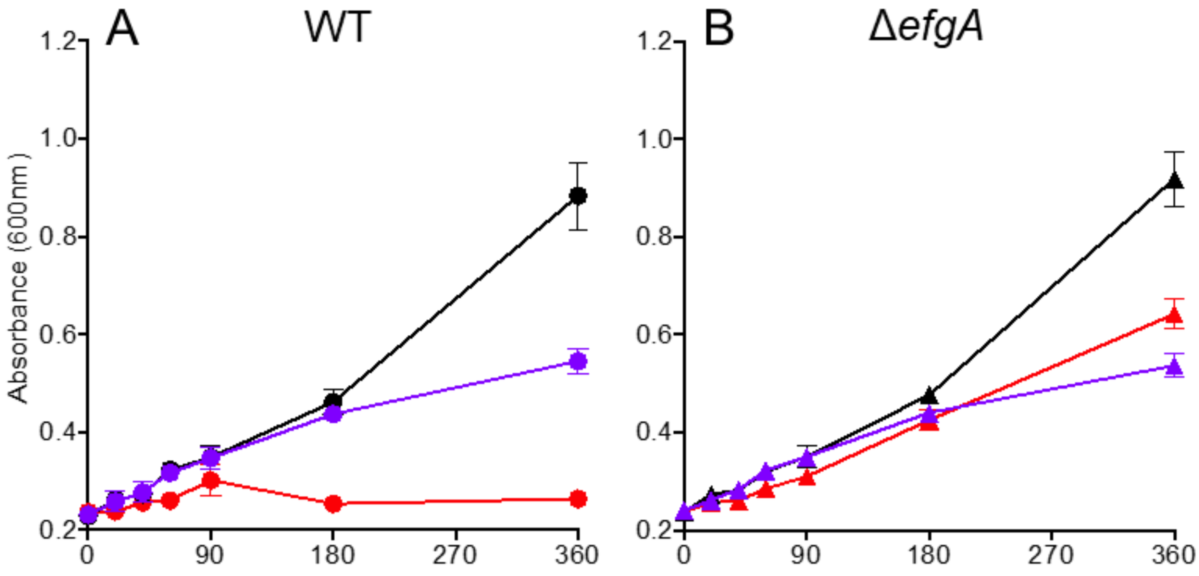


**Figure S19. EfgA causes rapid growth arrest in response to formaldehyde.** Exponential phase cultures of wild-type (circles, A) and Δ*efgA* mutant (triangles, B) strains were treated with kanamycin (purple), exogenous formaldehyde (red) or left untreated (black). Growth in response to treatment (t >0 min) was monitored. Error bars represent the standard error of the mean for three biological replicates.
